# Expression, subcellular localization, and phosphorylation of MK5 in adult cardiac ventricular fibroblasts

**DOI:** 10.1101/2020.07.24.219790

**Authors:** Pramod Sahadevan, Sherin A. Nawaito, Joëlle Trépanier, Sabrina Benamar, Fatiha Sahmi, Gabriel Theberge-Julien, Louis R. Villeneuve, Matthias Gaestel, Jean-Claude Tardif, Bruce G. Allen

**Author notes:** Address correspondence to: Bruce G. Allen, Montreal Heart Institute, 5000 Belanger St., Montréal, Québec, Canada, H1T 1C8.

## Abstract

MAP kinase-activated protein kinase-5 (MK5) plays an important role in cardiac fibroblast function. Although p38 MAPK and atypical MAPKs and ERK3 and ERK4 have been identified as activators of MK5, the kinases that activate MK5 remain controversial. Here we examined the expression, subcellular distribution, and regulation of MK5 in cardiac ventricular myofibroblasts and myocytes. The copy numbers for MK5 and ERK4 mRNA were comparable in myocytes and myofibroblasts, whereas that of ERK3 was much higher in myofibroblasts. Interestingly, MK5 and ERK3 immunoreactivity was detected in myofibroblasts but not myocytes whereas ERK4 immunoreactivity was detected in myocytes: treating in myocytes with a proteasome inhibitor or hypertrophic agonists failed to rescue MK5 immunoreactivity. In myofibroblasts, MK5 and ERK3 immunoreactivity was predominantly nuclear and cytosolic, respectively. In serum-starved cardiac myofibroblasts, phosphothreonine-182 MK5 (pT182-MK5) immunoreactivity was predominantly nuclear but increased in intensity and relocated to the cytoplasm in response to serum, sorbitol, angiotensin II, TGFβ, or H_2_O_2_ and this was prevented by inhibition of p38*α*/*β*. Phos-tag SDS-PAGE revealed multiple slower migrating bands of MK5 immunoreactivity, indicating phosphorylation of MK5 at multiple sites. Phos-tag PAGE also revealed MK5 phosphorylation was increased with fibroblast activation and in hearts exposed to a chronic increase in afterload. MK5 and ERK3 co-immunoprecipitated and proximity ligation assays revealed ERK3 and MK5 in close proximity in myofibroblast cytoplasmic compartment. Furthermore, p38*α*/*β* inhibition decreased the abundance of MK5 immunoreactivity in ERK3 immunoprecipitates. Finally, deleting MK5 did not reduce the abundance of ERK3 immunoreactivity. These observations suggest that p38*α* and/or p38*β* are the primary mediators of T182-MK5 phosphorylation and hence MK5 activation in cardiac myofibroblasts.

## Introduction

Mitogen-activated protein kinases (MAPKs) are protein serine/threonine kinases that are involved in various signal transduction pathways in all eukaryotic cells including plants, fungi and animals ^1, 2^. Activated MAPKs regulate a wide range of functions through phosphorylation of several substrates, including other protein kinases such as the MAPK-activated protein kinases (MAPKAPKs) ^3^. MAP kinase-activated protein kinase-5 (MK5), a member of MAPKAPKs, was first identified and characterized as p38-regulated/activated kinase (PRAK) ^4, 5^. Subsequent studies have reported various functionally diverse proteins including the atypical MAPKs ERK3/4 as interacting partners and/or activators of MK5 ^6–8^. A rich source of MK5 transcripts is the left ventricular myocardium ^9^. However, in addition to the molecular interactions for the activation, the *in vivo* consequence of such interactions to evoke various functions of MK5 remain poorly understood.

Both the heart and ventricular myocytes in male MK5 haploinsufficient mice showed significantly less hypertrophy in response to a chronic increase in afterload induced by constriction of the transverse aorta (TAC). In addition, the TAC-induced increase in cardiac collagen 1-*α*_1_ mRNA is significantly attenuated in MK5 haplodeficient mice relative MK5^+/+^ littermates ^10^. Furthermore, following myocardial infarction induced by ligation of the left anterior descending coronary artery, scar size, and collagen content were reduced in MK5-haploinsufficient mice compared with wildtype littermates ^11^. MK5 deficiency or haplodeficiency altered the abundance of numerous transcripts for proteins involved in extracellular matrix homeostasis. Reducing MK5 expression in cardiac myofibroblasts increases the abundance of collagen 1-*α*_1_ mRNA but attenuates the angiotensin-II (AngII)- and TGFβ-induced increase in collagen secretion and impairs both migration and contraction ^11, 12^. Hence, MK5 plays an important role in mediating fibroblast function.

The regulation of MK5 activity, mediated by phosphorylation at threonine-182 in the activation loop, remains controversial, with p38*α*/*β*, ERK3, and ERK4 having been implicated. However, PKA may also activate MK5 via phosphorylation at S115 ^13, 14^. ERK3 immunoreactivity, but not that of ERK4 or p38*α*, was detected in MK5 immunoprecipitates from mouse heart ^9^. Finally, MK5 immunoreactivity is detected in fibroblasts but not in myocytes ^10^. The present study was to examine MK5 phosphorylation in adult cardiac ventricular fibroblasts.

## Materials and Methods

### Materials

Medium 199 and fungizone were from Sigma-Aldrich, fetal bovine serum (FBS) and trypsin were from Gibco Laboratories (Life Technologies, Inc.), and penicillin-streptomycin solution was from Multicell. All culture plates were from Sarstedt, Inc. PCR Primers were from Invitrogen. SDS-polyacrylamide gel electrophoresis reagents, nitrocellulose, and Bradford protein assay reagents were from Bio-Rad Laboratories. Antibodies against MK5 were from Santa Cruz Biotechnology (PRAK A-7) and Cell Signaling Technology (MAPKAPK-5 #D70A10). Antibodies against phosphothreonine182-MK5 were from Abcam (pT182-MK5, ab138668). ERK3 antibodies were from Santa Cruz Biotechnology (ERK3 D23) and Thermo Fisher Scientific (ERK3 5E1). Antibodies to ERK4 were from Santa Cruz Biotechnology (ERK4 N20). Horseradish peroxidase (HRP)-conjugated secondary antibodies were from Jackson ImmunoResearch Laboratories. Proximity Ligation Assay kits from Sigma-Aldrich. Lambda protein phosphatase and alkaline phosphatase were from Calbiochem (#539514-20KV) and Sigma-Aldrich (#P6774-2KU), respectively. Collagen-coated hydrogel-bound (8 kPa) polystyrene 35mm (PS35-COL-8) and 6-well (SW6-COL-8) plates were from Matrigen. Phos-tag acrylamide was from Wako Chemicals (#AAL-107). All other reagents were of analytical grade or best grade available. Milli Q-grade (≥18.2 MΩ/cm) water was used throughout these studies.

### Mice

The line of constitutive MK5 knockout mice employed in these studies have been described previously ^10, 15^ and were on a mixed 129/Ola x C57BL background. Male 12- to 13-week-old wild type (MK5^+/+^) and MK5-deficient (MK5^-/-^) littermate mice were used. All animal experiments were approved by the institutional ethics committee and performed according to the guidelines of the Canadian Council on Animal Care.

### Isolation of Cardiac Ventricular Fibroblasts

Cardiac fibroblasts were isolated from 12–13 week-old MK5^+/+^ and MK5^-/-^ mice and young adult male Sprague-Dawley rats (151-175 g) as described earlier ^10^. Briefly, mice were sacrificed by cervical dislocation. The heart was excised and placed in sterile phosphate-buffered saline (PBS; 137 mM NaCl, 2.7 mM KCl, 4.2 mM Na_2_HPO_4_·H_2_O, 1.8 mM KH_2_PO_4_, pH 7.4) at 37°C. Hearts were trimmed and the atria were removed. The ventricular tissue was then macerated using scissors and subjected to a series of digestions in dissociation medium (116.4 mM NaCl, 23.4 mM HEPES, 0.94 mM NaH_2_PO_4_.H_2_O, 5.4 mM KCl, 5.5 mM dextrose, 0.4 mM MgSO_4_, 1 mM CaCl_2_, 1 mg/ml BSA, 0.5 mg/ml collagenase type IA, 1 mg/ml trypsin, and 20 μg/ml pancreatin, pH 7.4). Digestion was aided by gentle shaking on an orbital shaker maintained at 37°C. Digests were centrifuged at 1500 rpm for 5 minutes and the pellet was re-suspended in 4 ml of media M199 supplemented with 10% fetal bovine serum (FBS) and 2% penicillin-streptomycin. The cell suspension was plated onto two ‘complaint’ 35 mm collagen coated soft hydrogel-bound polystyrene plates (hydrogel stiffness 8 kPa) or ‘non-compliant’ 35 mm uncoated standard culture dishes and incubated in a humidified incubator at 37°C in a 5% CO_2_ atmosphere to obtain fibroblasts and activated fibroblasts, or myofibroblasts, respectively. The medium was changed after 150 min, to remove unattached cells and debris, and every 48 hours thereafter. Cultures from passages 0, 1, 2, and 3 were used at 80% confluency. Cells were washed with PBS and starved with serum-free M199 media for 24 h prior to treatment with 10% serum, 0.4 M sorbitol, 1 μM Ang-II, 1 μM TGF-β or 25 μM H_2_O_2_.

### Isolation of Rat and Mouse Cardiac Ventricular Myocytes

Cardiac ventricular myocytes were isolated from 12–14 weeks-old MK5^+/+^ mice and young adult male Sprague-Dawley rats (151-175 g) as described previously ^10^ and used immediately. Briefly, rats and mice were anesthetized with an intraperitoneal injection of a combination of sodium pentobarbital (0.55 mg/kg body weight) and heparin (1.0 U/kg body weight). The heart was exposed via sternotomy, rapidly excised, and immersed in ice-cold Tyrode solution (140 mM NaCl, 5.5 mM KCl, 1 mM MgCl_2_, 0.3 mM KH_2_PO_4_, 10 mM dextrose, 5 mM HEPES, 2 mM CaCl_2_, adjusted to pH 7.3 at room temperature with NaOH). The heart was mounted on an isolated heart perfusion system (Harvard Apparatus) via cannulation of the ascending aorta and the coronary arteries were perfused with modified Tyrode solution (140 mM NaCl, 5.5 mM KCl, 1 mM MgCl_2_, 0.3 mM NaH_2_PO_4_, 5 mM HEPES and 10 mM dextrose adjusted to pH 7.3 at room temperature with NaOH) containing 200 μM CaCl_2_. After 3 minutes of perfusion, the buffer was changed to a nominally Ca^2+^-free Tyrode solution and the perfusion continued for an additional 6 min. The heart was then enzymatically digested by recirculation with Ca^2+^-free Tyrode solution containing 0.5 mg/ml collagenase type II (perfusion times: 25 min for mice, 40 min for rat). Solutions were constantly aerated with carbogen gas (95% O_2_/5% CO_2_). Solutions and cells were maintained at 37°C throughout the isolation process. When the myocardial tissue had softened, the left ventricle (LV) was dissected away from the remainder of the heart and minced into small pieces in Kruftbrühe medium (100 mM potassium glutamate, 10 mM potassium aspartate, 25 mM KCl, 10 mM KH_2_PO_4_, 2 mM MgSO_4_, 20 mM taurine, 5 mM creatine, 0.5 mM EGTA, 20 mM glucose, 5 mM HEPES, and 0.1% (w/v) BSA, adjusted to pH 7.5 at room temperature with KOH). To promote cell dissociation, the solution and tissue fragments were gently triturated using a transfer pipette and the suspension was filtered through a 200-μm nylon mesh. The cell suspension was maintained at room temperature for 10 min to allow for gravitational separation of cardiomyocytes from non-cardiomyocytes. The supernatant (non-myocytes) was discarded or used for isolation of cardiac fibroblasts. The cell pellet, containing LV myocytes, was resuspended in Joklik’s minimal essential medium (25 mM NaHCO_2_, 1.2 mM MgSO_4_, 1 mM DL-carnitine, 1 mM CaCl_2_, adjusted to pH 7.5 with NaOH). Calcium-tolerant, rod-shaped ventricular cardiomyocytes (75–90% of all cells) were used for subsequent experiments.

### Cardiomyocyte Stimulation

After calcium reintroduction, viable rod-shaped cardiomyocytes (10^5^ cells) were plated onto laminin-coated culture dishes. Cardiomyocytes were incubated with angiotensin II (1 μM), endothelin-1 (50 nM), or norepinephrine (1 μM) for the following times: 0 min, 5 min, 10 min, 15 min, 30 min, 1 h, 2 h, 4 h, and 6 h. During this time, cells were maintained in a humidified incubator at 37°C under 5% CO_2_.

### Proteasome Inhibition

To examine the effects of proteasome inhibition, Ca^2+^-tolerant, rod-shaped ventricular rat cardiomyocytes were incubated with or without the proteasome inhibitor, MG132 (Sigma-Aldrich, St Louis, MO), at a final concentration of 10 μM or 20 μM for 1 and 24 h. Myocytes were lysed in ice-cold LB1 lysis buffer comprising 50 mM Tris, 20 mM β-glycerophosphate, 20 mM NaF, 5 mM EDTA, 10 mM EGTA, 1.0% TX-100, 1 mM NaVO_4_, 1 μM microcystin LR, 5 mM DTT, 10 μg/ml leupeptin, 0.5 mM PMSF, and 10 mM benzamidine (pH 7.5 at 4°C). Cell homogenates were centrifuged for 30 min at 40,000 ×g and 4°C and the supernatants reserved for immunoblot assay.

### MK5 and ERK3 Knockdown

As MK5-deficient and -haplodeficient mice experience a chronic reduction in MK5 levels, compensatory effects may have occurred. Hence, we also characterized fibroblasts isolated from MK5^+/+^ mice following a knockdown of MK5 using small inhibitory RNA (siRNA): referred to herein as MK5-kd. Additionally, as ERK3^-/-^ mice die within 24 h of birth, we also characterized fibroblasts from C57BL/6mice following a knockdown of ERK3 using siRNA: referred to herein as ERK3-kd. These experiments were performed using mice with the same genetic background. Cells (8 x 10^4^/well) were seeded into 12-well plates. Twenty-four h post sub-culture, the media was replaced with Opti-MEM containing Ambion Silencer-Select MK5 siRNA (5 pmol; catalog number: 4390771 ID: s69588 and s69586) or ERK3 siRNA (5 pmol; catalog number: 4390771 ID: s78451) and Lipofectamine 2000 (2 µl) (Invitrogen) and cells incubated for 19 h. Following an additional incubation in M199 with 10% FBS for 12 h, the cells were incubated in M199 containing with or without 10% FCS for subsequent experiments.

### Transverse Aortic Constriction

Transverse aortic constriction (TAC) was done as described previously ^16^ in adult (12-13 weeks) male mice anesthetized with isoflurane gas plus buprenorphine (0.05 mg/kg, intraperitoneal injection). A 7-0 silk suture was used to constrict the transverse aorta, between the left and right carotid arteries, around a 27 Gauge piece of stainless-steel tubing. The suture was knotted, and the steel tube removed. The result was a constriction of the transverse aorta of approximately 60%. Sham animals underwent the identical surgical procedure, but the aorta was not constricted. Eight-weeks after surgery, the hearts removed, snap-frozen in liquid nitrogen-chilled 2-methyl butane, and stored at −80°C. Pentobarbital, rather than isoflurane, was always used prior to sacrifice as isoflurane activates p38 in the mouse heart (data not shown).

### RNA Extraction and Quantitative PCR

Total RNA was extracted from cardiac fibroblasts and myocytes using RNeasy® Micro kits (Qiagen Inc.) following the manufacturer’s instructions. First-strand cDNA was synthesized from 20 μl reaction volume containing 500 ng of RNA, 100ng of random primers, 1x First Strand buffer (50 mM Tris-HCl pH 8.3, 75 mM KCl, 3 mM MgCl_2_), 0.5 mM dNTP, 10 mM DTT, 40 U RNaseOUT recombinant ribonuclease inhibitor, and 200 U of M-MLV reverse transcriptase using the RT^2^ First Strand cDNA Synthesis Kit from (Invitrogen) according to the manufacturer’s protocol. To detect mRNA variants of MK5, SYBER green-based quantitative real-time PCR (qPCR) was performed using ABI StepOnePlus instrument as described previously ^9^. Samples were assayed in triplicate with glyceraldehyde-3-phosphate dehydrogenase (GAPDH) as control. The expression relative to a reference sample was calculated using the ΔΔCT method. The forward and reverse primers used are listed in the **Table 1**.

**Table 1.**
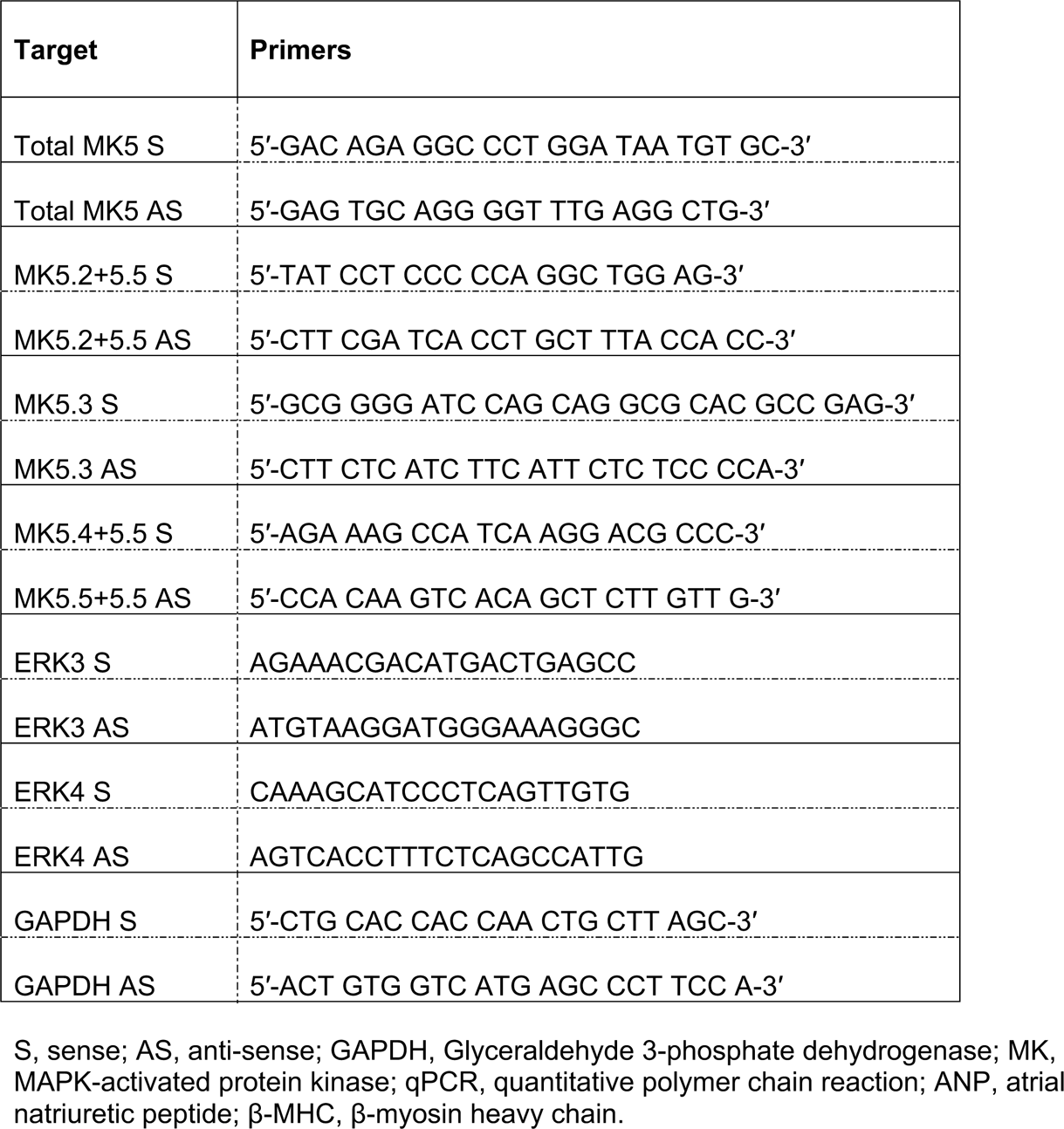
Primers used during real-time qPCR and / or ddPCR

Droplet digital PCR (ddPCR) was performed using a QX200 Droplet Digital PCR system from Bio-Rad Laboratories (Hercules, CA, USA). Reactions mixtures comprised cDNA equivalent to 10 ng total RNA along with 100 nM of forward and reverse primers (MK5, ERK3, and ERK4). After the preparation of 25 μl of reaction mix using QX200 ddPCR Eva Green Supermix (Bio-Rad), 20 μl was loaded into a droplet generation cartridge (Bio-Rad Laboratories, Hercules, USA) along with 70 μl of droplet generation oil for EvaGreen. The resulting ddPCR droplets were transferred to a clean 96-well plate and the plate was sealed with a foil lid using a thermal plate sealer from Bio-Rad Laboratories prior to PCR amplification using a Bio-Rad C1000 touch thermal cycler with deep wells. The amplification parameters were: 95°C for 5 min to activate the polymerase, followed by 40 cycles of 95°C for 30 s and 60°C for 1 min, ending with 95°C for 10 min to deactivate the polymerase. After amplification, the plate was transferred to a Bio-Rad droplet reader where raw fluorescence amplitude data was extracted from the Quantasoft software for downstream analysis. Each sample was run in duplicate.

### Electrophoresis and Immunoblot Assay

Three different electrophoresis procedures were followed. One involving standard 10% SDS-PAGE gels, second involving the in-gel combination of Phos-tag™ with Mn^2+^ (8% acrylamide gels supplemented with 50 µM Phos-tag reagent and 0.1 mM MnCl_2_ solution (Phos-tag gel)) and the third involving combination of Mn^2+^ Phos-tag gel with EDTA (Phos-tag gels supplemented with 10 mM EDTA (Phos-tag + EDTA gel)). For the standard SDS-PAGE, cells were lysed in ice-cold LB1 lysis buffer comprising 50 mM Tris (pH 7.5 at 4°C), 20 mM β-glycerophosphate, 20 mM NaF, 5 mM EDTA, 10 mM EGTA, 1.0% TX-100, 1mM Na_3_VO_4_, 1 μM microcystin LR, 5 mM DTT, 10 μg/ml leupeptin, 0.5 mM PMSF, and 10 mM benzamidine. For Phos-tag SDS-PAGE, cell and heart lysates were prepared in ice-cold EDTA-free lysis buffer comprising 50 mM Tris-HCl (pH 7.4), 150 mM NaCl, 1% NP-40, 0.5% sodium deoxycholate, 0.1% SDS with protease and phosphatase inhibitors. For phosphatase treatment, fibroblasts and myofibroblasts were lysed in ice-cold EDTA-free lysis buffer comprising 50 mM Tris-HCl (pH 7.4), 150 mM NaCl, 1% NP-40 with protease inhibitors. The running buffer for both standard SDS-PAGE and Phos-tag SDS-PAGE was 25 mM Tris base, 192 mM glycine, and 0.1% w/v SDS. Standard SDS-PAGE was at 100 volt/gel for 90 min. However, importantly, for Phos-tag SDS-PAGE, gels were run at 12 mA/gel for 2 h. Protein loading on Phos-tag SDS-PAGE ± EDTA was exactly the same. EDTA chelates the Mn^2+^, preventing the formation of a Phos-tag-Mg^2+^ complex in the gel that would otherwise interact with phosphoryl groups and retard the motility of phosphorylated forms of MK5. Following electrophoresis, proteins were transferred electrophoretically onto nitrocellulose membranes for 90 min at 100 volts in a buffer containing 10 mM Na-CAPS (pH 11.0) and 10% methanol or 60 min at 100 volts in a buffer containing 25 mM Tris base, 192 mM glycine and 20% methanol. Before transfer, Phos-tag SDS-PAGE gels were washed twice with Na-CAPS transfer buffer containing 10 mM EDTA for 10 min and once with Na-CAPS transfer buffer alone for 10 min with gentle agitation on an orbital shaker. Following the transfer, membranes were blocked for 1 h at room temperature in TBS containing 0.1% (v/v) Tween-20 (TBST) and 5% (w/v) non-fat dried milk. Membranes were then incubated with the indicated primary antibodies diluted 1:1000 in TBST containing 1% (w/v) BSA for 16 h at 4°C, washed 3 times in TBST, and incubated with the appropriate horseradish peroxidase-conjugated secondary antibody (Jackson ImmunoResearch Laboratories, Inc., West Grove, PA, USA) diluted 1:10,000 in TBST containing 5% (w/v) nonfat dried milk. Immunoreactive bands were visualized using Western Lightning Plus ECL reagent (PerkinElmer BioSignal Inc., Montreal, Canada) and Kodak BioMax Light film. Films were digitized using a Bio-Rad GS-800 densitometer and band intensities quantified using Bio-Rad Quantity One software.

### Two-Dimensional IEF–SDS-PAGE Immunoblotting

Myofibroblast and heart lysates were precipitated and resuspended in rehydration buffer (see below). Briefly, ice-cold trichloroacetic acid (TCA) was added to a final concentration of 10% (w/v) using a 100% (w/v) TCA stock solution, samples were vortexed, incubated on ice for 60 min, and then centrifuged for 15 min at 18,300 x g and 4°C. The protein pellet was washed twice with ice-cold acetone and dried in SpeedVac for 5 min without heat. The protein pellets were solubilized in rehydration buffer (8 M Urea, 2% CHAPS, 25 mM DTT, 0.2% Bio-Lyte 3/10 ampholyte solution, bromophenol blue for faint color) at room temperature. Samples were resolved on 7 cm pH 3-10 immobilized pH-gradient (IPG) strips (Bio-Rad), under mineral oil, using a Bio-Rad Protean IEF cell with the following protocol: active rehydration at 50 V for 12 h, 250 V for 15 min, a linear gradient of 250-to-10,000 V over 3 h, 10,000 V for 4 h, followed by 500 V until the samples were removed from the IEF cell. After the run, the excess mineral oil was removed by gently touching the edges of the IPG strips to tissue paper. The strips were incubated in equilibration buffer I (0.375 M Tris pH 8.8, 6 M urea, 2% SDS, 20% glycerol, and 130 mM DTT) and then equilibration buffer II (0.375 M Tris pH 8.8, 6 M urea, 2% SDS, 20% glycerol, and 135 mM iodoacetamide) for 10 min each with gentle shaking on an orbital mixer. Strips were rinsed free of excess equilibration buffer by dipping 3-4 times in SDS-PAGE running buffer, positioned above a 1.5-mm 10% acrylamide gel, and overlaid with molten agar (1% agarose) in SDS-PAGE running buffer. Once the agar had set, the second dimension SDS-PAGE gels were run, transferred to nitrocellulose, and MK5 immunoreactivity revealed by immunoblotting as described above.

### Immunoprecipitation Assays

Prior to immunoprecipitation, 2 μg of anti-MK5 antibody or anti-ERK3 antibodies was precoupled to 10 μl of a 30 mg/ml suspension of Dynabeads magnetic beads (Invitrogen) by incubating for 2 hours at 4°C with constant mixing followed by three washes with PBST, using a magnet rack to immobilize the beads, to remove any uncoupled antibody. Cardiac fibroblast lysates (1 mg) were added to the antibody-coated beads and incubated overnight at 4°C with constant mixing on a clinical rotator. Immunoprecipitates were washed three times with PBST, suspended in 20 μl of 2X Laemmli sample buffer, and heated at 70°C for 90 s. The beads were immobilized using a magnetic rack and the supernatants immediately loaded onto 10% acrylamide SDS-PAGE gels.

### Immunocytofluorescence Microscopy

Cardiac fibroblasts were cultured in 35 mm FluoroDish cell culture dishes. The cells were rinsed with PBS, fixed for 20 min at room temperature using PBS containing 2% (v/v) formaldehyde, and washed again with PBS. After permeabilization and blocking with PBS containing 0.2% (v/v) Triton X-100 and 2% (v/v) normal donkey serum, cells were incubated overnight at 4°C with primary antibodies (1:100). The next morning, cells were washed three times with PBS and incubated in the dark with the appropriate Alexa Fluor 555-conjugated secondary antibody (1:500, Life technologies) and DAPI (1:1000) for 1 h at room temperature, rinsed three times with PBS, and mounted on glass slides using 15 μl of DABCO/glycerol. Fluorescence images were acquired using a Zeiss LSM-510 confocal fluorescence microscope. Confocal z-stacks were deconvolved (Huygens Professional, Scientific Volume Imaging) using experimentally derived point-spread functions and 3-dimensional images rendered using Volocity Visualization software (Quorum Technologies Inc).

### Proximity Ligation Assay

Fibroblasts were fixed with 4% paraformaldehyde (20 min), permeabilized with 0.2% Triton X-100 (1 h) and blocked with 1% (w/v) BSA (10 min) at room temperature. Incubation with primary antibodies was overnight at 4°C. Incubation with kit-specific secondary antibodies (anti-rabbit MINUS Sigma Cat. DUO92005, anti-goat PLUS Sigma Cat. DUO92003), ligation, amplification, detection, mounting and wash buffers were as specified by the manufacturer’s instructions (Olink Bioscience, Sweden) using the Duolink *in situ* PLA probes and the Duolink In Situ Detection Reagent FarRed (Sigma Cat. no. DUO92013). Amplification was for 60 min. Slides were mounted with Duolink mounting medium containing DAPI and bound Cy5-conjugated oligonucleotide probes were detected using a Zeiss LSM 510 confocal microscope (63 × oil objective/1.4; Cy5 λ _ex_ 633 nm, λ _em_ 633 nm; DAPI λ _ex_ 405 nm, λ _em_ 461 nm).

### Statistical Analysis

Results are reported as the mean ± standard error (SEM). Statistical differences between two groups were performed using unpaired, two-tailed Student’s t-tests. A value of *P* less than 0.05 was considered statistically significant. Statistical analyses were performed using GraphPad Prism version 8.3 for the Mac OSX.

## Results

### The abundance of MK5, ERK3, and ERK4 mRNA differs between cardiac myofibroblasts and myocytes

In addition to p38, ERK3 and ERK4 have been identified as potential activators of MK5. Previous work from our laboratory identified 4 novel splice variants of MK5 mRNA (MK5.2-MK5.5) in murine ventricular myocardium ^9^ and showed the presence of MK5 mRNA in adult ventricular myocytes and fibroblasts ^17^. In this study, we examined the copy number of MK5, ERK3, and ERK4 mRNA in cardiac myofibroblasts and myocytes using droplet digital PCR. The primers used to detect the mRNA for total MK5 (all known splice variants), ERK3, and ERK4 are listed in Table I. Digital PCR revealed the copy number of MK5 transcripts was similar in myocytes and myofibroblasts. In contrast, the copy number of ERK3 mRNA was significantly elevated in myofibroblasts, whereas ERK4 mRNA was more abundant in myocytes (**Figure 1A**).

**Figure 1.**
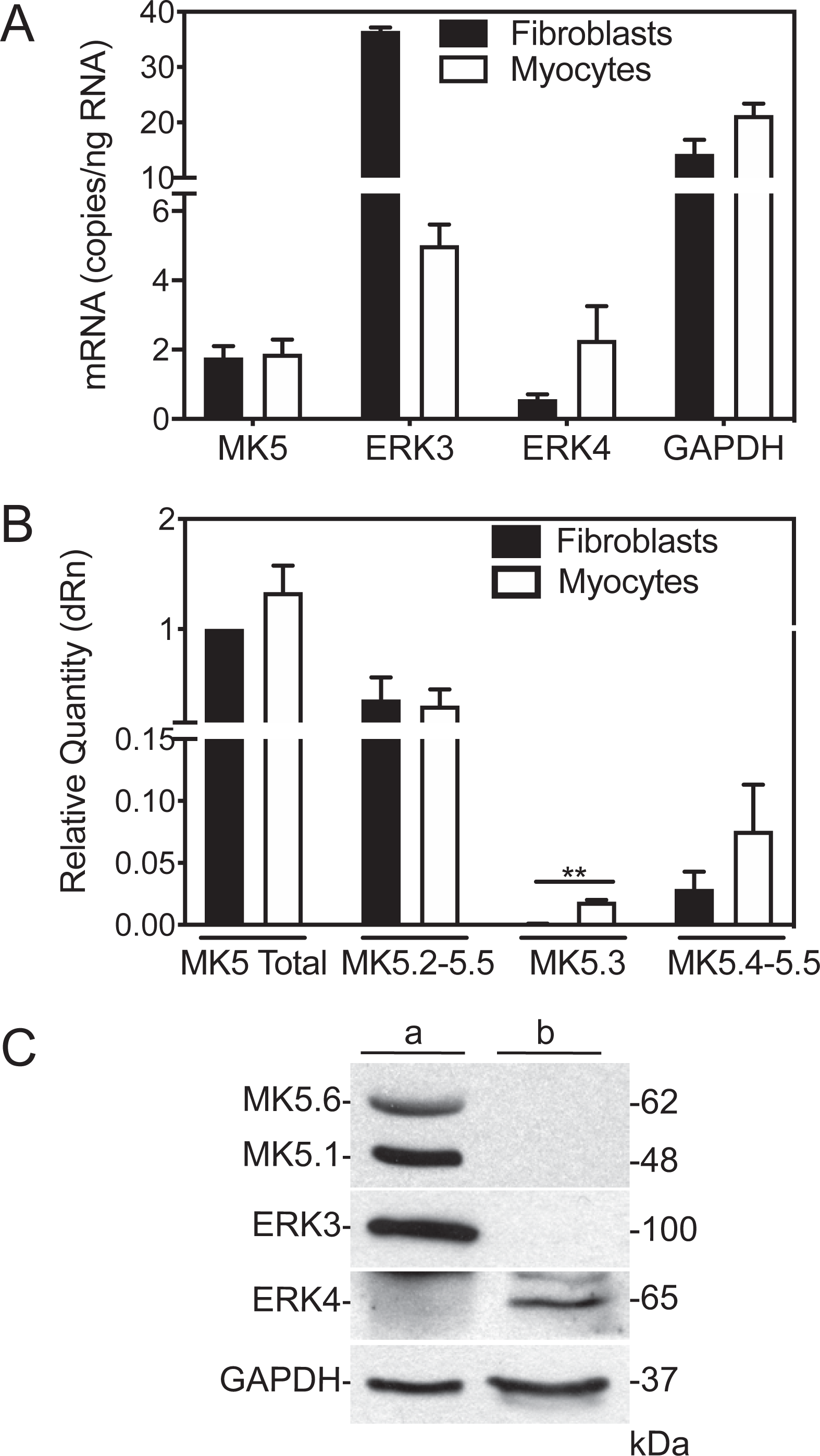
MK5, ERK3, and ERK4 mRNA and immunoreactivity in adult cardiac ventricular myofibroblasts and myocytes. **(A)** The abundance of MK5, ERK3, ERK4, and GAPDH mRNA was measured by droplet digital PCR in RNA isolated from MK5^+/+^ adult mouse cardiac ventricular myofibroblasts and myocytes. Values in the graph represent mean ± SEM of three independent experiments. Each experiment was performed with myofibroblasts and myocytes isolated from a separate mouse. **(B)** MK5 variant mRNA levels were quantified by qPCR in RNA isolated from MK5^+/+^ cardiac myofibroblasts and myocytes and are normalized to the abundance of GAPDH mRNA. Values in the graph represent mean ± SEM of three independent experiments. Each experiment was performed with myofibroblasts isolated from a separate mouse. **(C)** Immunoblot assay of MK5, ERK3, and ERK4 in myofibroblasts and myocytes. Cardiac myofibroblasts and myocytes lysates (20 μg/lane) were prepared, resolved by SDS-PAGE, and transferred to nitrocellulose. Two MK5-immunoreactive bands (MK5.1, 48-kDa; MK5.6, 62-kDa) were detected in myofibroblast lysates (*lane a*), whereas no immunoreactivity was detected in myocyte lysates (*lane b*). Membranes were reprobed for ERK3, ERK4, and GAPDH. ERK3-immunoreactive band (100 kDa) was detected in myofibroblast lysates (*lane a*) but not myocyte lysates (*lane b*). ERK4 (65-kDa) immunoreactivity was detected in myocyte lysates (*lane b*) but not myofibroblast lysates (*lane a*). GAPDH immunoreactivity was used as a loading control. Numbers at the *right* indicate molecular mass (in kDa). These results are qualitatively similar to those obtained from three separate cell preparations.

### The abundance of transcripts for MK5 splice variants is similar in cardiac myofibroblasts and myocytes

We next sought to determine the relative quantity of each MK5 variant transcript (MK5.1-MK5.5) in cardiac ventricular myofibroblasts and myocytes using qPCR. Primers (**Table I**) for total MK5 (i.e., all 5 variants), MK5.2+MK5.5, MK5.3, and MK5.4+MK5.5 were as described previously ^9^. The abundance of splice variant mRNAs was expressed relative to the abundance of total MK5 mRNA (**Figure 1B**). MK5 splice variants were detected in both myofibroblasts and myocytes. MK5.1 was the most abundant variant, whereas MK5.3 was the lowest in both myofibroblasts and myocytes. The variants MK5.2+MK5.5 represented 35.9% of the total pool of MK5 mRNA in cardiac myofibroblasts and 22.9% in myocytes. MK5.3 was 0.1% and 1.4% for myofibroblasts and myocytes, respectively. The abundance of MK5.4+MK5.5 was 2.9% in myofibroblasts and 5.7% in myocytes.

### The abundance of MK5, ERK3, and ERK4 immunoreactivity differs between myofibroblasts and myocytes

We next examined the abundance of MK5, ERK3, and ERK4 immunoreactivity in myofibroblasts and myocytes. Cardiac ventricular myocytes and myofibroblasts were isolated from adult mice, lysates prepared, and resolved by SDS-PAGE. As shown previously, two bands of MK5 immunoreactivity were detected, one at 48-kDa and another at 62-kDa in cardiac fibroblasts, corresponding to MK5.1 and MK5.6 respectively, whereas MK5 immunoreactivity was not detected in myocytes (**Figure 1C**) ^10^. Similarly, ERK3 immunoreactivity was detected in the myofibroblasts but not in myocytes. In contrast, ERK4 immunoreactivity was only detected in myocytes (**Figure 1C**). These results suggest that, although MK5, ERK3, and ERK4 transcripts are present in both cardiac fibroblasts and myocytes, they are translated differentially in these cells. Taken together, these observations suggest ERK3, ERK4, and MK5 play cell-specific roles in the heart and that ERK4 may be a member of a novel signaling cascade in cardiac myocytes that does not involve MK5 as a substrate.

### Proteasome inhibition did not rescue MK5 immunoreactivity in cardiac myocytes

Previous studies have suggested that MK5 might play multiple roles in cardiac pathophysiology ^9–12^. As proteasome-meditated degradation is one of the important mechanisms for regulation *in vivo* protein levels, we next sought to determine if the absence of MK5 immunoreactivity in myocytes was a result of rapid, proteasome-meditated degradation. To address this, adult cardiac ventricular myocytes were treated with the proteasome inhibitor MG132 for 1 (**Figure 2A**) and 24 h (**Figure 2B**), lysed, and subjected to immunoblot assays. MG132 treatment failed to rescue MK5 immunoreactivity. Hence, the absence of MK5 immunoreactivity in cardiac myocytes may reflect the inability of these cells to translate MK5 transcripts.

**Figure 2.**
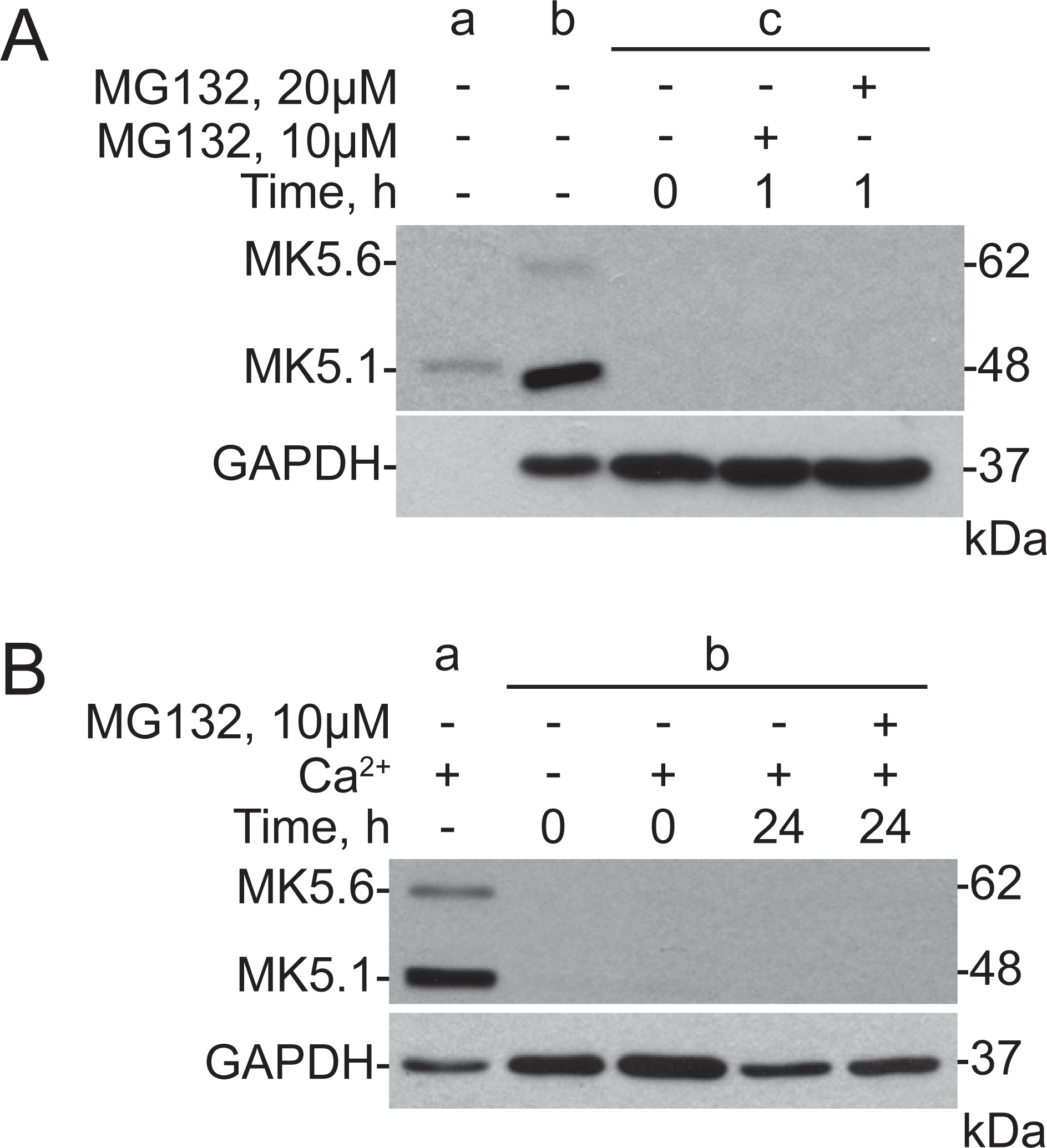
Proteasome inhibition did not rescue MK5 immunoreactivity in cardiomyocytes. **(A)** Ventricular myocytes, isolated from adult rat hearts, were treated with 10 or 20 μM MG132 for 1 h. Total cell lysates were then prepared for immunoblot assay. Lysates from human embryonic kidney 293 cells transfected with pIRES-MK5.1-V5 (40 ng) and rat cardiac myofibroblasts (20 μg) were applied to lane a and lane b, respectively. No MK5 immunoreactivity was detected in myocyte lysates with or without MG132 treatment (lanes labeled c). GAPDH immunoreactivity was used as a loading control. **(B)** Ventricular myocytes, isolated from adult rat hearts, were treated with or without 10 μM MG132 for 24 h. Total cell lysates were then prepared for immunoblot assay. MK5-immunoreactive bands of 48- and 62-kDa were detected in cardiac myofibroblast lysates (20 μg/lane) from rat (*lane a*) whereas no MK5 immunoreactivity was detected in myocyte lysates (*lanes labeled b*). To maintain myocytes for 24 h, they were cultured in media containing CaCl_2_. Freshly isolated myocyte preparations were divided into 4 conditions and one condition was lysed without addition of CaCl_2_-containing culture media. GAPDH immunoreactivity was used as a loading control. These results are qualitatively similar in 3 separate myocyte preparations.

### Angiotensin II, endothelin 1, or norepinephrine failed to induce MK5 translation in cardiac ventricular myocytes

We next asked if MK5 mRNA would be translated in cardiac myocytes in response to pro-hypertrophic agonists. Angiotensin-II (AngII), endothelin-1 (ET-1), and norepinephrine (NE) are G protein-coupled receptor agonists that promote cardiac hypertrophy ^18, 19^. Adult rat ventricular myocytes were incubated with Ang-II, ET-1, or NE for various times up to 6 h, lysed, and the presence of MK5 protein immunoreactivity was assessed by immunoblot assay. No MK5 immunoreactive bands were detected in myocyte lysates as a result of these treatments (**Figure 3A-C**). Myocytes are isolated in nominally Ca^2+^-free buffer, following which Ca^2+^ is added in a stepwise manner. The presence or absence of Ca^2+^ in the culture media did not alter the inability to detect MK5 immunoreactivity. Hence, pro-hypertrophic stimuli failed to induce the translation of MK5 transcripts in adult ventricular myocytes and further studies focused on MK5 in cardiac ventricular fibroblasts.

**Figure 3.**
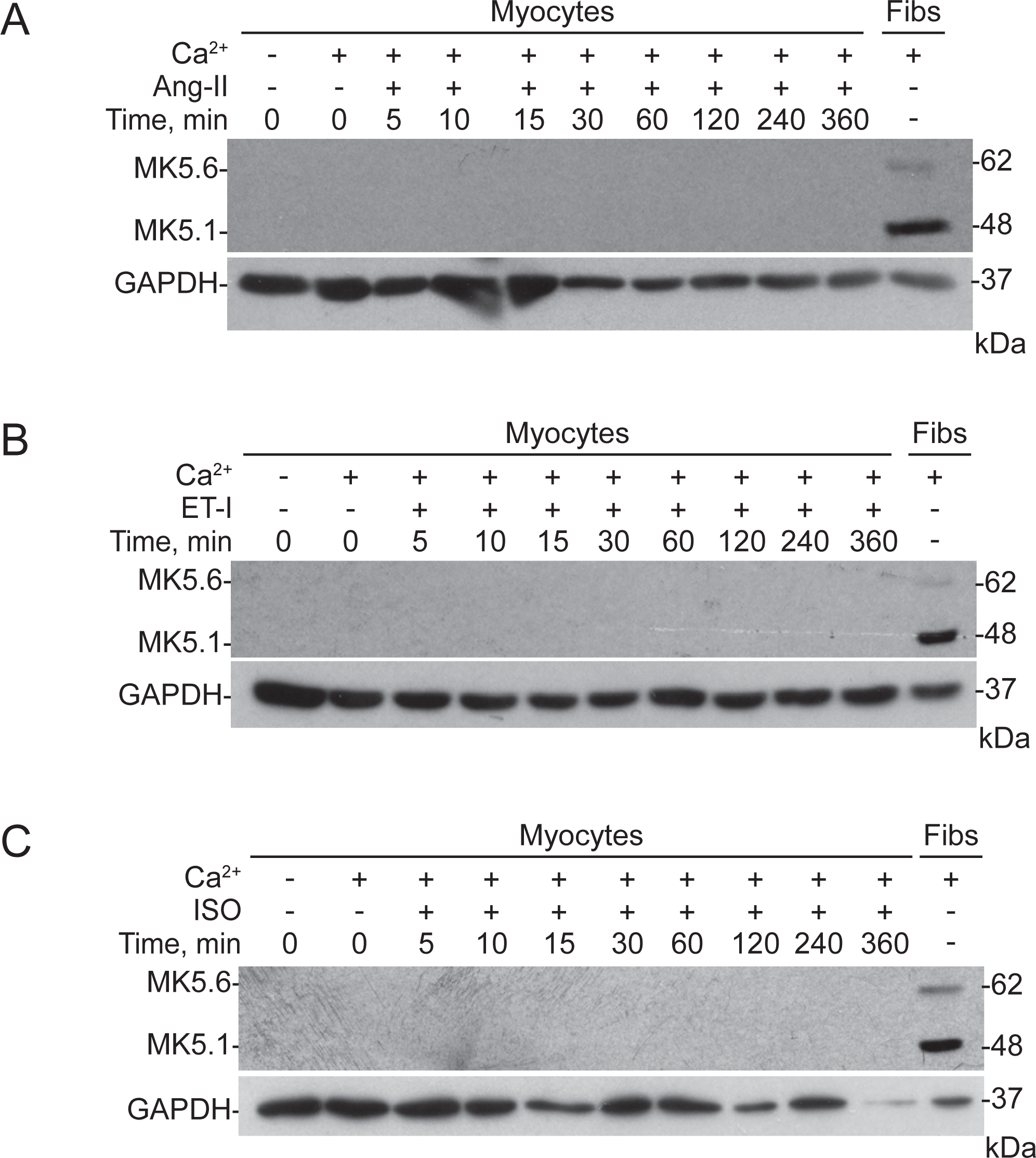
Incubation of rat cardiac ventricular myocytes with prohypertrophic agonists angiotensin II, endothelin 1, or norepinephrine did not induce translation of MK5 mRNA. Ventricular fibroblasts and myocytes were isolated from young adult rat hearts. Myocytes were stimulated with **(A)** angiotensin II (1 μM), **(B)** endothelin-1 (50 nM), and **(C)** norepinephrine (1 μM) for 0 - 360 min, as indicated. Fibroblasts (Fibs) were isolated and maintained in culture until passage 3. Lysates (25 μg/lane) were prepared, resolved by SDS-PAGE, and transferred to nitrocellulose. MK5-immunoreactive bands of 48- and 62-kDa were detected in fibroblast lysates, whereas no MK5 immunoreactivity was detected in myocyte lysates. GAPDH was used as a loading control. The images shown are representative immunoblots. The results were qualitatively similar in three independent experiments: each experiment was performed using cells isolated from a separate animal. Numbers at the right indicate the position of molecular mass markers (in kDa).

### Subcellular localization of ERK3 and MK5 in cardiac myofibroblasts

Knowing the subcellular localization of ERK3 and MK5 in fibroblasts should help to understand their function under physiological and pathophysiological conditions. Several reports demonstrated that MK5 is primarily located in the nucleus, whereas ERK3 is located in both nucleus and the cytoplasm ^20–26^. Consistent with previous reports ^21, 23, 25^, confocal immunofluorescence microscopy revealed MK5 immunoreactivity to be primarily in the nucleus but also in the perinuclear region in myofibroblasts (**Figures 4A,B**). In contrast, ERK3 immunoreactivity was predominantly in the cytoplasm (**Figures 5A,B**). We next investigated whether there was an effect of passage number on the subcellular distribution of MK5 and ERK3 immunoreactivity in myofibroblasts. However, no changes in the subcellular localization of ERK3 and MK5 immunofluorescence was observed in myofibroblasts from passage numbers 0-3 (**Figures 4A,B & 5A,B**). Hence, in serum-starved myofibroblasts, ERK3 and MK5 were located in different subcellular compartments.

**Figure 4.**
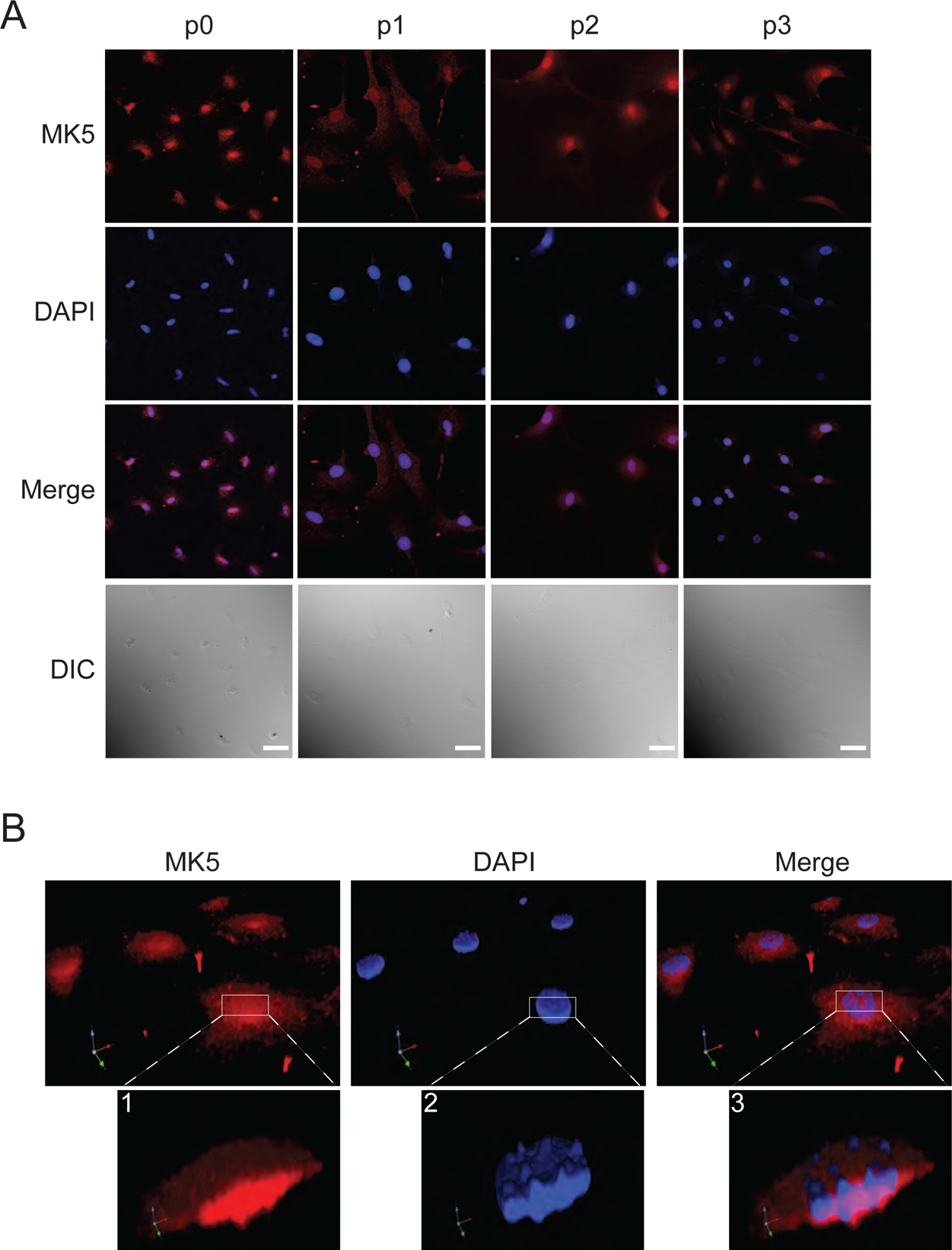
Subcellular distribution of MK5 immunoreactivity in mouse cardiac ventricular myofibroblasts. **(A)** Cardiac myofibroblasts (continuously serum-fed) from passage numbers (p) 0-3 were fixed and decorated with an anti-MK5 antibody followed by an Alexa Fluor 555-coupled secondary (anti-rabbit) antibody. MK5 immunofluorescence is pseudo colored red. Nuclei were visualized using the DNA-binding fluorescent dye, DAPI (pseudo colored blue). Merged refers to an overlay of MK5 and DAPI fluorescence. Myofibroblasts were visualized by differential interference contrast (DIC) microscopy. Results shown are representative of three qualitatively similar experiments performed using different fibroblast preparations. **(B)** Three-dimensional rendering (Velocity 3D image Analysis Software) of a deconvolved (Huygens Professional, Scientific Volume Imaging) confocal z-stack showing MK5 immunofluorescence (*red*) is predominantly located in the nucleus in cardiac myofibroblasts. DNA was visualized by staining with DAPI (*blue*). The lower panels of Figure 4B show higher magnification images of the rendered volumes defined by the white boxes in the upper panels.

**Figure 5.**
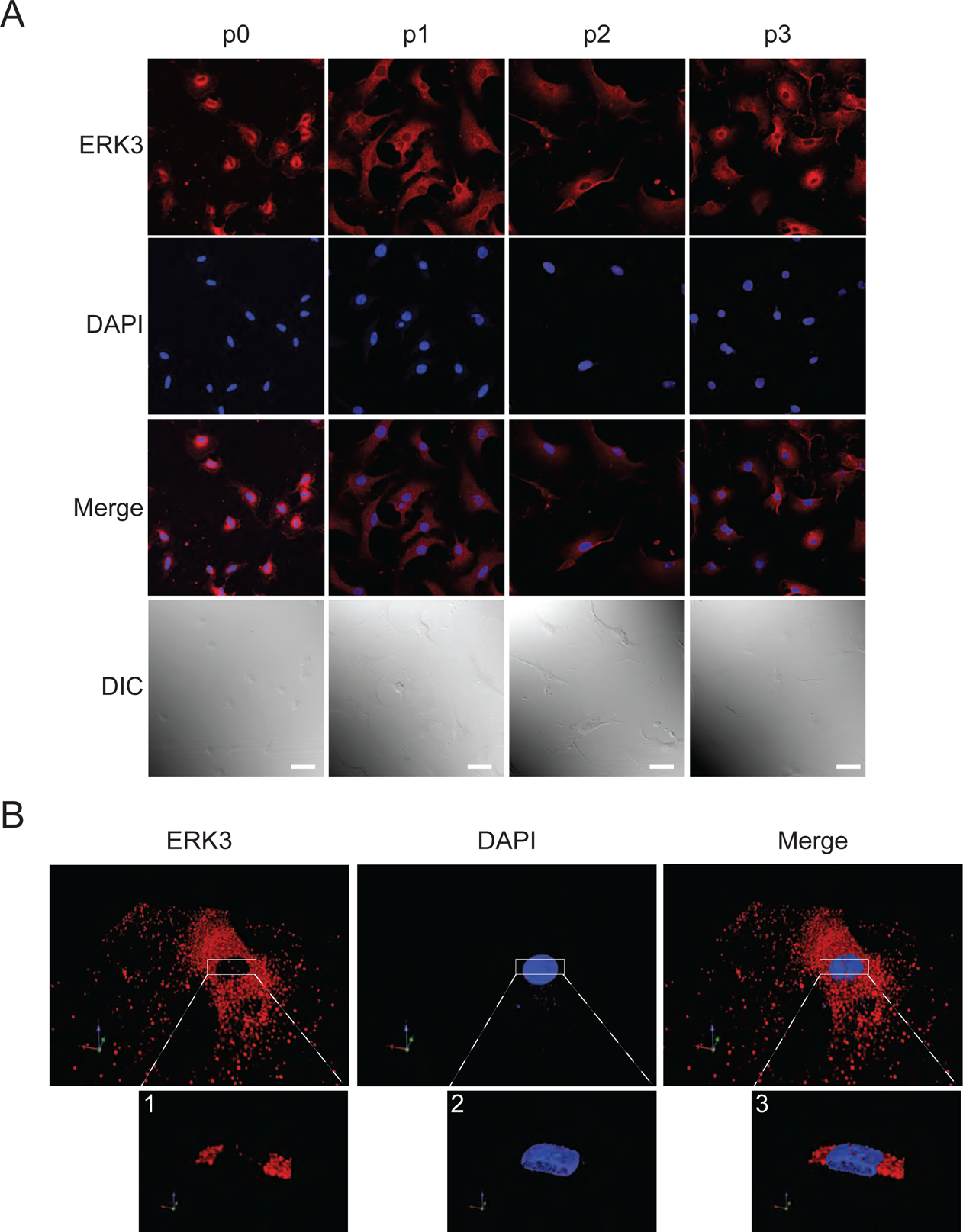
Subcellular distribution of ERK3 immunoreactivity in mouse cardiac ventricular myofibroblasts. **(A)** Cardiac myofibroblasts (continuously serum-fed) from passage numbers (p) 0 - 3 were fixed and decorated with an anti-ERK3 antibody followed by an Alexa Fluor 555-coupled secondary (anti-rabbit) antibody. ERK3 immunofluorescence is pseudo colored red. Nuclei were visualized using the DNA-binding fluorescent dye, DAPI (pseudo colored blue). Merged refers to an overlay of ERK3 and DAPI fluorescence. Myofibroblasts were visualized by differential interference contrast (DIC) microscopy. **(B)** Three-dimensional rendering (Velocity 3D image Analysis Software) of a deconvolved (Huygens Professional, Scientific Volume Imaging) confocal z-stack showing ERK3 immunofluorescence (red) is primarily located in the cytoplasm. DNA was visualized by staining with DAPI (blue). The lower panels of Figure 5B show higher magnification images of the rendered volumes defined by the white boxes in the upper panels. Results shown are representative of three qualitatively similar experiments performed using different fibroblast preparations.

Activation of MK5 involves phosphorylation at threonine-182 (T182) in its activation loop. In serum-starved cardiac myofibroblasts, phospho-threonine-182 MK5 (pT182-MK5) immunofluorescence was predominantly nuclear (**Figure 6A, time 0**). However, following mitogen (e.g., serum) stimulation, pT182-MK5 immunofluorescence was observed in the cytoplasm (**Figure 6A**). As shown in **Figure 6B**, in actively dividing (serum was never withdrawn) myofibroblasts, pT182-MK5 immunofluorescence appeared to be associated with the cytoskeleton and pseudopodia. No pT182-MK5 immunofluorescence was detected in actively dividing myofibroblasts isolated from MK5^-/-^ mice (**Figure 6C**). ERK3 immunofluorescence localized to membrane ruffles and/or lamellipodia following serum stimulation (**Figure 6D**). We then examined the requirements for p38 and ERK3 activity on the ability of serum to increase Interestingly pT182-MK5 immunofluorescence. Pretreating fibroblasts with an inhibitor of p38*α*/*β* activity, SB203580 (20 μM), blocked the serum-dependent increase in pT182-MK5 immunofluorescence induced by serum (**Figure 7A**). Note that p38*γ* and p38*δ* do not phosphorylate MK5.1 ^9, 27^. An siRNA approach was employed to interfere with ERK3 activity. Immunoblot assays using two different siRNA concentrations (a, b) confirmed the knockdown was effective (**Figure 7B**). Further, the acute knockdown of ERK3 resulted in a diffuse cytosolic distribution of pT182-MK5 immunoreactivity but failed to attenuate the increase in intensity evoked by serum stimulation (**Figure 7A**). We next examined the effect of a diverse array of stimuli, including hyperosmolar stress (e.g., 0.4 M sorbitol), hypertrophic agonists (e.g., AngII and TGF*β*), and oxidative stress (e.g., H_2_O_2_) on the intensity and subcellular localization of pThr182-MK5 immunoreactivity (**Figure 8**). In each case, pT182-MK5 immunoreactivity both increased in intensity and relocated to the cytoplasm and the perinuclear region. Both the increase in intensity and redistribution of pT182-MK5 immunoreactivity was prevented by SB203580. These results suggest p38α and/or p38*β* are the primary mediators of MK5 phosphorylation at Thr182 in cardiac myofibroblasts, whereas ERK3 may be responsible for targeting activated MK5 to specific subcellular locations.

**Figure 6.**
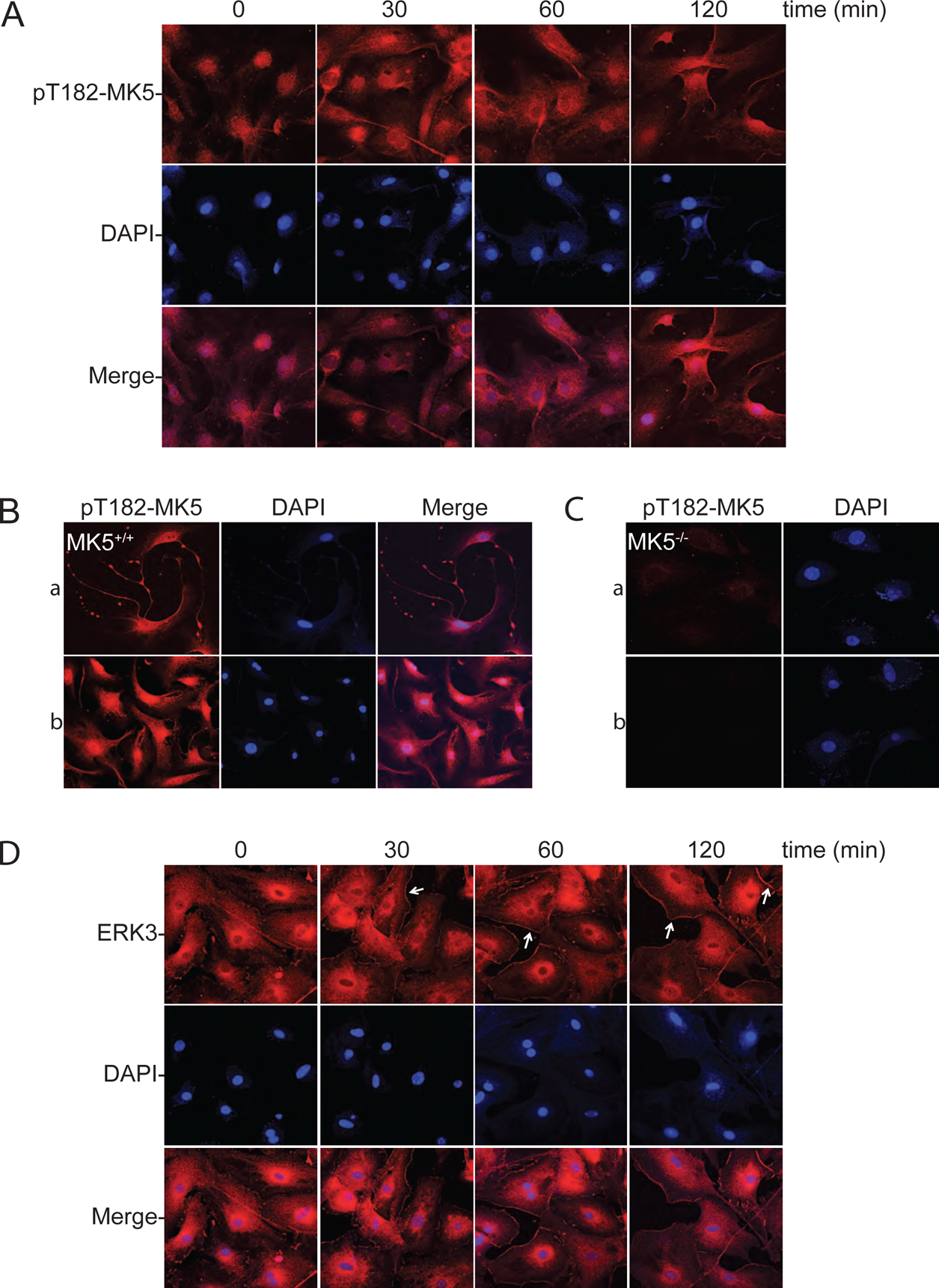
Serum-stimulation alters the subcellular distribution of pT182-MK5 and ERK3 immunoreactivity in myofibroblasts. **(A)** Subconfluent quiescent passage 3 cardiac myofibroblasts in M199 were serum-starved and then stimulated with 10% serum for the indicated times. The subcellular localization of phosphothreonine182-MK5 (pT182-MK5) immunoreactivity was revealed by decorating fibroblasts with an anti-pT182-MK5 antibody (Abcam #ab138668) followed by an Alexa Fluor 555-coupled secondary (rabbit) antibody and visualized by confocal fluorescence microscopy. Nuclei were visualized using the DNA-binding fluorescent dye, DAPI (pseudo colored blue). Merged refers to an overlay of pT182-MK5 and DAPI fluorescence. **(B)** Subconfluent MK5^+/+^ cardiac myofibroblasts from passage numbers 0 (a) and 3 (*b*) were fixed and the subcellular distribution of pT182-MK5 immunofluorescence examined as described for panel A. **(C)** Subconfluent passage 2 MK5^-/-^ cardiac myofibroblasts were fixed and the subcellular distribution of pT182-MK5 immunofluorescence examined by incubating in the presence (a) or absence (b) of an anti-pT182-MK5 antibody (Abcam #ab138668) followed by an Alexa Fluor 555-coupled anti-rabbit secondary antibody and visualized by confocal fluorescence microscopy as described for panel A. **(D)** Subconfluent quiescent cultures of cardiac passage 3 myofibroblasts in M199 were serum-starved and then stimulated with 10% serum for the indicated times. The subcellular distribution of ERK3 immunoreactivity was revealed by decorating fibroblasts with an anti-ERK3 antibody followed by an Alexa Fluor 555-coupled secondary (rabbit) antibody and visualized by confocal fluorescence microscopy. Nuclei were visualized using the DNA-binding fluorescent dye, DAPI (pseudo colored blue). Merged refers to an overlay of ERK3 and DAPI fluorescence. Results shown are representative of three qualitatively similar experiments performed using different fibroblast preparations.

**Figure 7.**
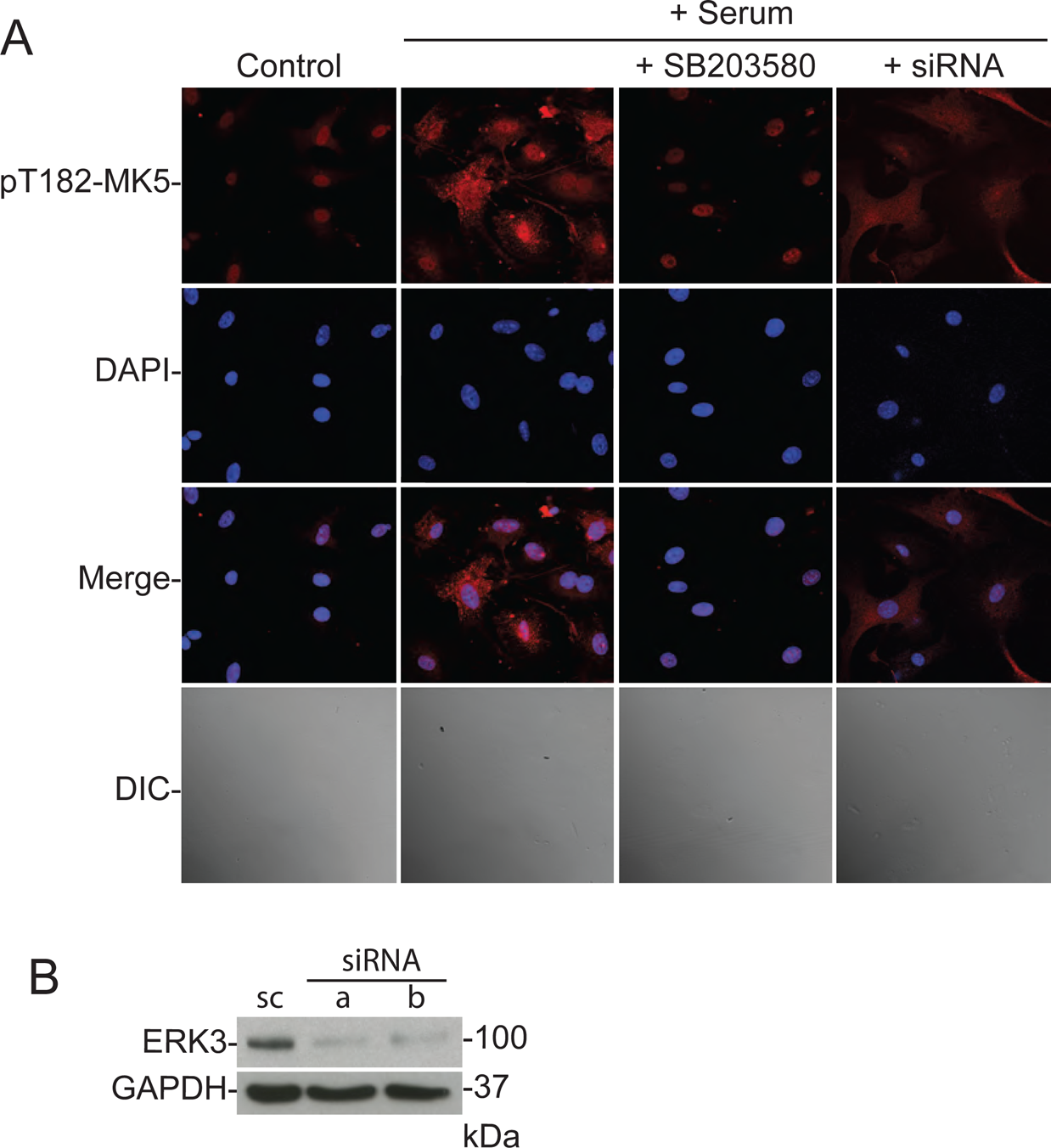
p38α/β inhibition and ERK3 knockdown alter the subcellular distribution of pT182-MK5 immunoreactivity in serum-stimulated cardiac myofibroblasts. **(A)** Myofibroblasts (*passage 2*) were transfected with ERK3-directed siRNA or scrambled siRNA (control), as described in Methods. After 24 h of serum deprivation, subconfluent cultures of cardiac myofibroblasts in M199 were stimulated with 10% serum for 2 h in presence or absence of SB203580 (20 μM). Where indicated, myofibroblasts were preincubated with SB203580 for 15 min prior to the addition of serum. The subcellular localization of pT182-MK5 immunoreactivity was determined by confocal fluorescence microscopy after decorating fibroblasts with a pT182-MK5-specific antibody (Abcam #ab138668) followed by an Alexa Fluor 555-conjugated anti-rabbit secondary antibody. Nuclei were visualized using the DNA-binding fluorescent dye, DAPI (pseudo colored blue). Merged refers to an overlay of pT182-MK5 and DAPI fluorescence. Myofibroblasts were visualized by differential interference contrast (DIC) microscopy. **(B)** Myofibroblasts were transiently transfected with either siRNA for ERK3 (si; a = 5 nM, b = 10 nM) or a scrambled RNA sequence (sc, 5 nM) as described under Methods. The abundance of ERK3 immunoreactivity was examined by immunoblot assay. ERK3 immunoreactive bands were detected in the myofibroblast lysates transfected with scrambled RNA sequence. The intensity of ERK3 bands was reduced, relative to scrambled RNA, upon ERK3 knockdown using a single siRNA. Numbers at the right indicate the position of molecular mass markers (in kDa). Results shown are representative of three qualitatively similar experiments performed using different fibroblast preparations.

**Figure 8.**
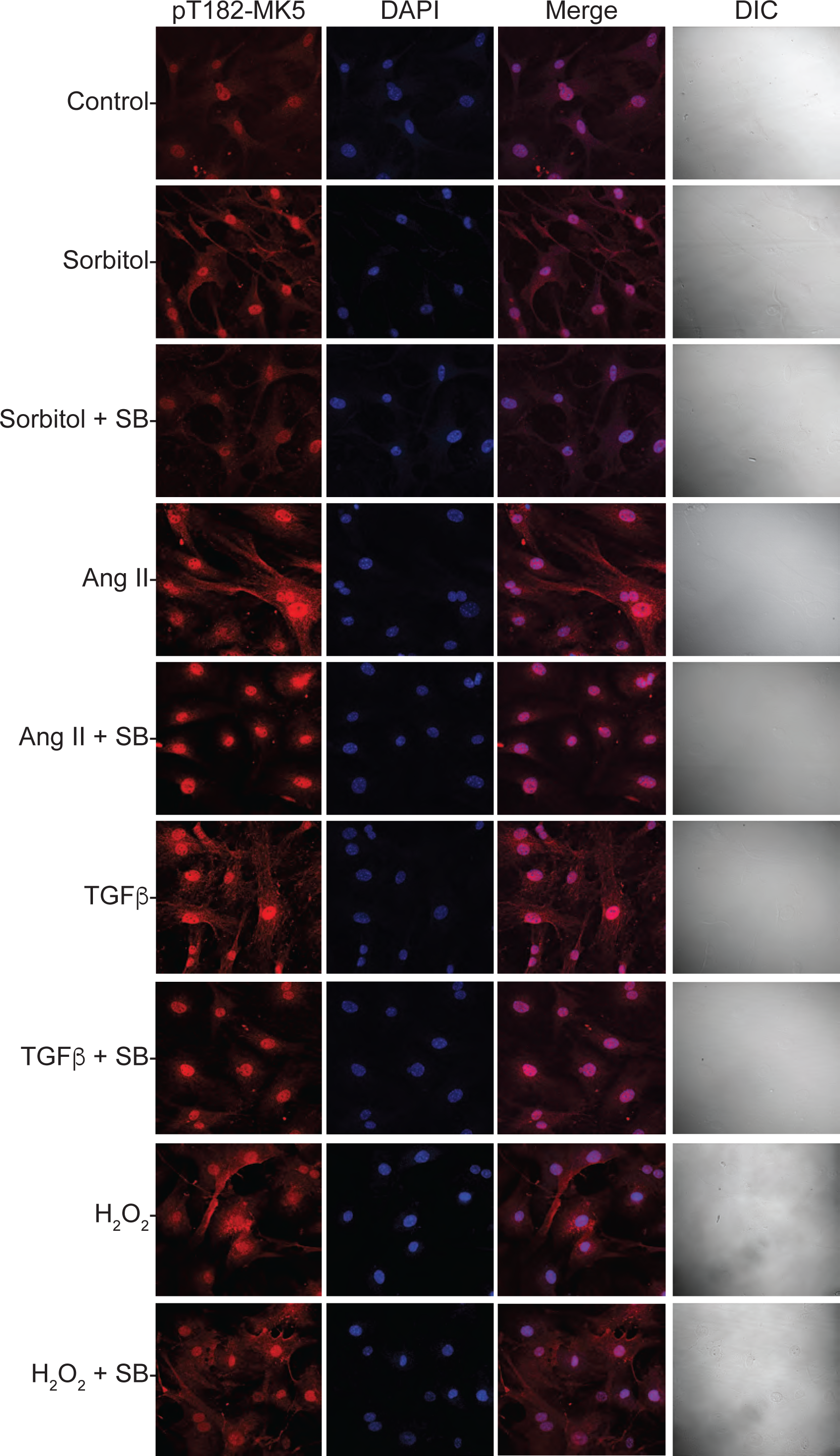
p38α/β inhibition prevents the subcellular relocalization of pT182-MK5 immunoreactivity induced by sorbitol, angiotensin II, TGFβ, and H_2_O_2_. After 24 h of serum deprivation, subconfluent cultures of cardiac myofibroblasts (*passage 2*) in M199 media were stimulated with 0.4 M sorbitol, 1 μM Ang II, 1 μM TGFβ, or 25 μM H_2_O_2_ for 2 h in the presence or absence of SB203580 (20 μM). Where indicated, cells were preincubated with SB203580 for 15 min prior to stimulation. The subcellular localization of pT182-MK5 immunoreactivity was determined by confocal fluorescence microscopy after decorating fibroblasts with a pT182-MK5-specific antibody (Abcam #ab138668) followed by an Alexa Fluor 555-conjugated anti-rabbit secondary antibody. Nuclei were visualized using the DNA-binding fluorescent dye, DAPI (pseudo colored blue). Merged refers to an overlay of pT182-MK5 and DAPI fluorescence. Myofibroblasts were visualized by differential interference contrast (DIC) microscopy. Results shown are representative of three qualitatively similar experiments performed using different fibroblast preparations.

### MK5 Immunoreactive Profile in 2D-PAGE and Phos-tag SDS-PAGE

The available anti-pT182-MK5 antibodies revealed numerous non-specific bands in immunoblot assays using fibroblast lysates (data not shown). Furthermore, changes in pT182-MK5 immunoreactivity would only provide information concerning phosphorylation at a single site. To have a more global view of changes in MK5 phosphorylation, two electrophoretic techniques that will separate phosphorylated forms of MK5, two-dimensional gel electrophoresis (2DE) and Phos-tag SDS-PAGE, were employed. Two-dimensional gel electrophoresis separates proteins based on both their isoelectric point (p*I*) ^28, 29^ and molecular mass. Posttranslational modifications such as phosphorylation can alter both the p*I* and apparently molecular mass of a protein ^28, 29^. As shown above, immunoblot analysis following SDS-PAGE MK5 revealed two bands of MK5 immunoreactivity, 48 and 62 kDa, corresponding to MK5.1 and MK5.6, respectively (**Figure 1C**). To determine if detectable total MK5 comprised 4 immunoreactive spots corresponding to the non-phosphorylated and pT182 forms MK5.1 and MK5.6, total heart lysates were prepared from MK5^+/+^ and MK5^-/-^ mice and resolved on 2-dimensional gel electrophoresis comprising immobilized pH gradient (IPG) strips (pH 3–10) in the first dimension and SDS-PAGE in the second. Gels were then transferred to nitrocellulose and MK5 immunoreactive spots revealed. Surprisingly, MK5 immunoreactivity with an apparent molecular mass corresponding to that of MK5.1 was resolved into a large number of spots by 2DE (**Figure 9A**) which were attenuated or absent in lysates from MK5^-/-^ hearts (**Figure 9B**). Multiple spots of total MK5 immunoreactivity in the 2DE profile suggested MK5 was phosphorylated at sites in addition to threonine-182. No immunoreactive spots were observed that would correspond to MK5.6. In subsequent studies, we employed Phos-tag SDS-PAGE to examine the phosphorylation status of MK5 as it permitted multiple samples to be run on the same gel and MK5.6 immunoreactivity was not lost. In the presence of a divalent cation such as Mn^2+^, the Phos-tag reagent interacts with, and reduces the electrophoretic mobility of, phosphorylated proteins whereas chelating the Mn^2+^ ions using EDTA allows separation based upon apparent molecular mass. Fibroblasts were isolated from mouse and rat left ventricle, lysed, treated with or without lambda and alkaline phosphatases, resolved on Phos-tag-SDS-PAGE gels cast without or with 10 mM EDTA, transferred to nitrocellulose, and total MK5 immunoreactivity revealed. In each case, protein loading was the same. In fibroblast lysates from both mouse and rat heart, Phos-tag SDS-PAGE (-EDTA) resolved total MK5 immunoreactivity into multiple bands that included those identified as MK5.1 and MK5.6 on Figure 1C plus additional diffuse bands migrating slower than MK5.1 and MK5.6 (**Figure 9C, upper panel lanes 1 and 3**). MK5.1 appeared to migrate as a doublet. Pretreating these fibroblast lysates with phosphoprotein phosphatases (lambda phosphatase followed by alkaline phosphatase) eliminated the slower mobility immunoreactive forms, yielding a profile of immunoreactivity resembling that observed following SDS-PAGE (**Figure 1C**) with MK5.1 and MK5.6 clearly visible (**Figure 9C, upper panel lanes 2 and 4**) and mouse MK5.1 migrating as a single band. When the samples shown in the upper panel of Figure 9C were resolved on Phos-tag SDS-PAGE containing 10 mM EDTA (**Figure 9C, middle panel**) no slower-migrating forms of MK5 immunoreactivity were observed: both with or without phosphoprotein phosphatase pretreatment with; only immunoreactive bands corresponding to MK5.1 and MK5.6 were observed. A comparison of the banding patterns in Figure 9C suggests that most of MK5.6 and a large proportion of MK5.1 were phosphorylated with a stoichiometry of at least 1. These observations suggest that, in cardiac ventricular myofibroblasts, MK5 is phosphorylated at multiple sites.

**Figure 9.**
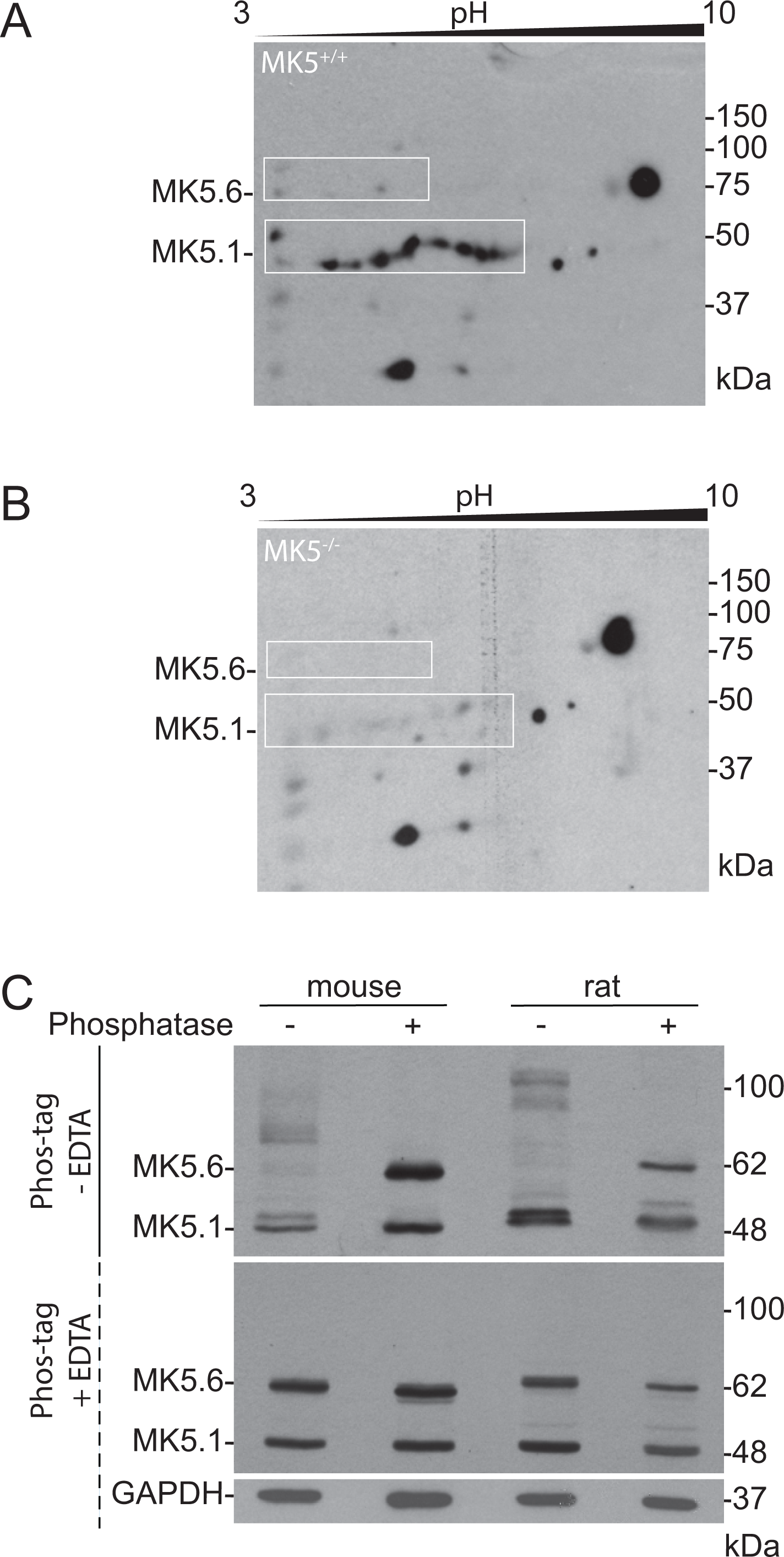
MK5 immunoreactivity migrates as multiple species on 2-dimensional gel electrophoresis and Phos-tag SDS-PAGE. Proteins were extracted from **(A)** MK5^+/+^ and **(B)** MK5^−/−^ hearts, resolved on 2-dimensional gel electrophoresis (3DE; immobilized pH gradient gel electrophoresis, SDS-PAGE), transferred to nitrocellulose, and probed for total MK5 immunoreactivity as described in Methods. Numerous MK5-immunoreactive spots were detected in lysates from MK5^+/+^ mice **(*A*)** that were attenuated or absent in lysates from MK5^-/-^ mice **(B)**. Numbers above indicate the pH gradient in the first dimension and those at the right indicate the position of molecular mass markers in the second dimension (in kDa). **(C)** Resolution of total MK5 immunoreactivity on Phos-tag SDS-PAGE in the presence or absence of EDTA. Lysates were prepared from mouse and rat passage 2 cardiac myofibroblasts lysates. Where indicated, lysates were incubated with phosphoprotein phosphatases (*λ* phosphatase followed by alkaline phosphatase) to dephosphorylate MK5 prior to separation on Phos-tag SDS-PAGE. MK5 immunoreactive bands were detected by immunoblot assay with MK5-specific antibodies. After probing for MK5 immunoreactivity, the Phos-tag + EDTA membrane was stripped and reprobed for GAPDH immunoreactivity, which was used as a loading control. Numbers at the right indicate the position of molecular mass markers (in kDa). A representative immunoblot is shown.

We next undertook a bioinformatic analysis of phosphorylation sites and their cognate protein kinases in mouse MK5.1 (NP_034895.1) using version 5.0 of the Group-based Prediction System (GPS5; http://gps.biocuckoo.cn) with a high threshold ^30^. Using a high threshold, GPS5 predicted MK5 could be phosphorylated at 10 tyrosine, 28 serine, and 19 threonine residues (Table 2) by a large number of protein serine/threonine, protein tyrosine and dual-specificity kinases. Comprehensive list of kinases predicted to phosphorylate MK5 at serine/threonine and tyrosine residues are provided in Supplemental Tables 1 and 2, respectively. GPS-5 has the advantage over other tools for predicting phosphorylation sites and the kinases that may phosphorylate these sites as it predicts for 479 human protein kinases. Although a mouse-specific module is available, GPS-5 failed to predict T182 as a site for p38 when using the mouse-specific module, so the default module was employed herein. To experimentally determine sites phosphorylated in MK5, lysates were prepared from serum-fed myofibroblasts and either MK5 or ERK3 were immunoprecipitated. Immune complexes were submitted to a proteomics core facility for tryptic digest and LC-MS/MS. However, attempts to identify the phosphorylation sites in MK5 by LC-MS/MS have been unsuccessful, to date, as no peptide sequences corresponding to MK5 have been obtained in either MK5 or ERK3 immunoprecipitates.

**Table 2.** Predicted protein kinases and phosphorylation sites in mouse MK5.1

| Amino acid | Site | Cognate protein kinase |
| --- | --- | --- |
| tyrosine (10) | Y-20, Y-188, Y-189, Y-216, Y-218, Y-232, Y-238, Y-242, Y-375, Y-425 | Abl, Ack, Axl, Csk, DDR, EGFR, Eph, FGFR, Fak, InsR, Met, PDGFR, Src, Syk, Tec, Trk, VEGFR, BAZ, DYRK, MAPK, STE7, LRRK, PEK, WEE |
| serine (28)<br>threonine (19) | S-2, S-5, T-14, S-15, S-21, T-25, S-33, S-43, T-44, T-71, S-85, S-92, S-93, S-115, T-121, S-126, T-129, S-160, T-182, T-186, S-206, T-211, S-212, T-214, T-217, S-221, S-226, S-243, S-247, T-249, T-260, S-262, S-271, S-274, T-294, S-306, T-307, S-316, S-348, S-354, T-368, T-383, S-386, S-438, S-440, T-447, T-467 | Akt, DMPK, GRK, NDR, PDK1, PKA, PKC, PKG, RSK, SGK, CAMK, CAMK1, CAMK2, CAMKL, DAPK, MAPKAPK, MLCK, PHK, PIM, PKD, RAD53, TSSK, CK1, VRK, CDK, CDKL, CK2, CLK, DYRK, GSK, MAPK, RCK, STE, STE11, STE20, STE7, IRAK, LISK, LRRK, MLK, RAF, RIPK, STKR, Alpha, BCR, GTF2F1, PDHK, PIKK, RIO, Aur, BUB, Bud32, CAMKK, CDC7, Haspin, IKK, KIS, MOS, NAK, NEK, NKF2, PEK, PLK, TLK, TOPK, TTK, ULK, WEE, WNK, InsR, Lmr, Src, Tec |

Group-based Prediction System (GPS; http://gps.biocuckoo.cn) output for protein tyrosine, protein serine/threonine, and dual-specificity kinase phosphorylation sites and cognate protein kinases in mouse MK5.1 (NCBI NP\_034895.1).

The serum-induced increase in pT182-MK5 immunoreactivity was blocked by SB203580 (20 μM), so we next examined the role of p38*α*/*β* in the mobility shifts in MK5 immunoreactive bands observed on Phos-tag SDS-PAGE in lysates from both serum-starved fibroblasts with or without re-addition of serum (**Figure 10A**). In both serum-starved and serum-stimulated (2 h) fibroblasts, no band was detected for the non-phosphorylated form of MK5.6. Inhibiting p38 did not prevent the phosphorylation of MK5.6 although two diffuse bands were observed to appear or increase in intensity in serum-starved and serum-stimulated fibroblasts in the presence of SB203580 (indicated by asterisk). The phosphorylation state of MK5.1 differs from that of MK5.6. In serum starve fibroblasts, a band corresponding the unphosphorylated form of MK5.1 was detected (**Figure 10A, upper panel**) forming a doubled with a larger band of slightly lower electrophoretic mobility. In the presence of SB203580 (20 μM), the MK5 immunoreactivity appears to shift towards the lower band in this doublet. A similar pattern was observed for MK5.1 immunoreactivity in serum-stimulated fibroblasts; however, the abundance of immunoreactivity in both bands was reduced relative to that from the serum-starved fibroblasts. Immunoreactive bands corresponding to MK5.1 and MK5.6 were evident when the samples were resolved on Phos-tag SDS-PAGE containing 10 mM EDTA (**Figure 10A, middle panel**). Hence, p38*α*/*β* are not the only protein kinases that phosphorylate MK5 in myofibroblasts.

**Figure 10.**
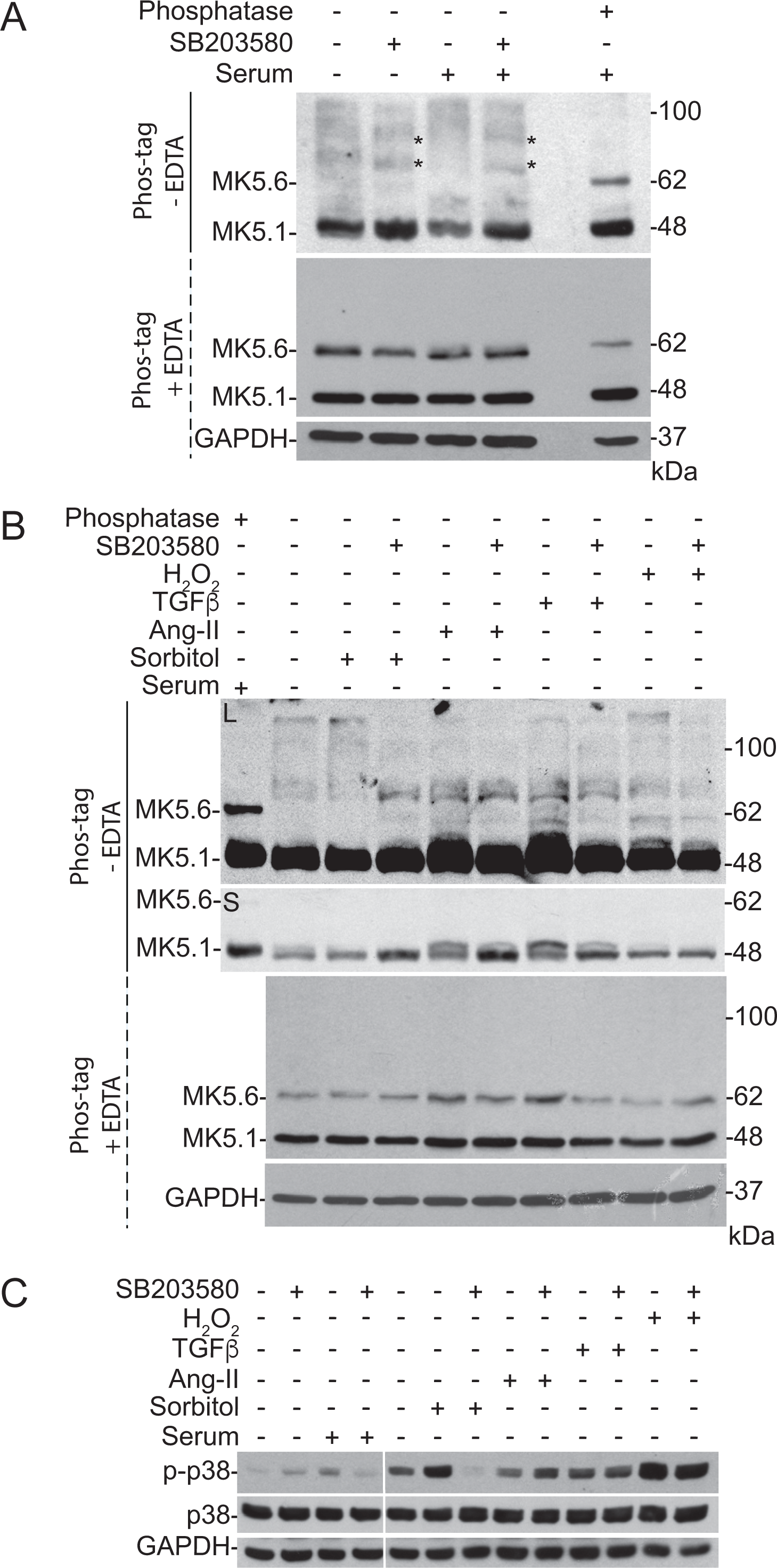
Phos-tag SDS-PAGE reveals lower mobility forms of MK5 immunoreactivity in serum-, sorbitol-, angiotensin II-, TGFβ-, and H_2_O_2_-treated myofibroblasts. After 24 h of serum deprivation, sub confluent passage 2 myofibroblasts were treated with **(A)** 10% serum or **(B)** 0.4 M sorbitol, 1 μM Ang II, 1 μM TGFβ, or 25 μM H_2_O_2_ for 2 h in the presence or absence of SB203580 (20 μM). Where indicated, cells were preincubated with SB203580 for 15 min prior to stimulation. Lysates were prepared and each sample applied to both Phos-tag SDS-PAGE - EDTA and Phos-tag SDS-PAGE +EDTA followed by immunoblot assay for total MK5 immunoreactivity. Phosphatase (*λ* phosphatase followed by alkaline phosphatase) treated rat cardiac myofibroblast lysate was used as a dephosphorylated control sample on Phos-tag SDS-PAGE -EDTA. After probing for MK5 immunoreactivity, the Phos-tag SDS-PAGE +EDTA membrane was stripped and reprobed for GAPDH immunoreactivity, which was used as a loading control. Numbers at the right indicate the position of molecular mass markers (in kDa). Asterisks in Panel A indicate bands of MK5 immunoreactivity that increase in intensity with SB203580 treatment. (**C)** After 24 h of serum deprivation, subconfluent passage 2 cardiac myofibroblasts were treated with 10% serum, 0.4 M sorbitol, 1 μM Ang II, 1 μM TGFβ, or 25 μM H_2_O_2_ for 2 h in the presence or absence of SB203580 (20 μM). Where indicated, cells were preincubated with SB203580 for 15 min prior to the stimulation. Lysates were prepared, resolved on SDS-PAGE followed by immunoblot assay for phospho-TGY-p38 and p38α. GAPDH was used as an internal loading control. Results shown are representative of three qualitatively similar experiments performed using different fibroblast preparations.

To further characterize MK5 phosphorylation, serum-starved mouse LV myofibroblasts were exposed to hyperosmotic stress (0.4 M sorbitol), AngII, TGFβ, or H_2_O_2_, for 2 h, in the absence and presence of SB203580 (20 μM), lysed, and resolved on Phos-tag SDS-PAGE (**Figure 10B**). As described above, a basal level of MK5 phosphorylation was observed in serum-starved myofibroblasts with no band corresponding to MK5.6 detected, diffuse bands of lower electrophoretic mobility, and MK5.1 migrating as a doublet. In Figure 10B, two exposures (long, short) of the Phos-tag SDS-PAGE -EDTA immunoblot are provided to reveal both the faint, slower migrating bands (long exposure) while also providing clear images of the MK5.1 doublet (short exposure). Each of the stimuli employed affected the profile of MK5 immunoreactivity on Phos-tag SDS-PAGE, relative to untreated serum-starved fibroblasts (lane 2), albeit to different extents: they increased the abundance of various slower-migrating forms of MK5 immunoreactivity on Phos-tag PAGE. Although SB203580 affected some of the slower migrating bands (**Figure 10B, upper panel, L**), it was not able to block phosphorylation of MK5.6.

A shorter exposure to film allowed for a better assessment of MK5.1 (**Figure 10B, upper panel, S**). As with actively growing (constantly serum-fed) myofibroblasts (**Figure 9C**), in serum-starved myofibroblasts MK5.1 migrated as a doubled on Phos-tag SDS-PAGE. Looking at the effects of the stimuli examined in this study, hyperosmotic stress (0.4 M sorbitol) cause a reduced intensity of the upper band in the MK5.1 doublet, and sorbitol plus SB203580 increased the intensity of the lower band. Both AngII and TGF*β* increased the intensity of the upper band and reduced that of the lower band and this response was prevented by SB203580. The effects of H_2_O_2_ resembled those of osmotic stress whereas H_2_O_2_ plus SB203580 failed to produce the increase in the non-phosphorylated form of MK5.1 (lower band of the doublet) observed with sorbitol plus SB203580. Hence these stimuli differed in terms of their: 1) effects on the behavior of MK5 on Phos-tag SDS-PAGE and 2) sensitivities to inhibition of p38*α*/*β*, suggesting the involvement of additional protein kinases in phosphorylating MK5. Resolving these same samples on Phos-tag SDS-PAGE containing EDTA confirmed the presence of both MK5.1 and MK5.6 in each sample (**Figure 10B, lower panel**).

Figure 10C confirms p38 was phosphorylated at the T-G-Y motif in the activation loop region; however, as these cells were exposed to stimuli for 2 h, and desensitization may have occurred in certain cases, the levels of phosphorylation shown in this figure do not represent the magnitude of p38 activation. Furthermore, as SB203580 inhibits the activity of p38*α*/*β*, a decrease in phospho-T-G-Y immunoreactivity would indicate a stimulus acting via the non-canonical TAB1-mediated activation of p38*α*, which involves TAB1 binding p38*α*, inducing a conformational change, resulting in p38*α* autophosphorylation within the T-G-Y motif ^31, 32^. Taken together, these results support the involvement of p38α/β in phosphorylating MK5 at T182 but that MK5 is phosphorylated at additional sites by kinases other than p38 in cardiac fibroblasts.

### Phosphorylation of MK5 is altered in mouse hearts exposed to a chronic increase in afterload

We have shown previously that, in hearts exposed to a chronic increase in afterload induced by constriction of the transverse aorta (TAC), MK5 haplodeficiency attenuated both the increase in collagen 1-a1 mRNA and myocyte hypertrophy ^10^. The abundance of total MK5 immunoreactivity in heart lysates was not altered by TAC ^10^. Hence, we next used Phos-tag SDS-PAGE to determine if the phosphorylation status of MK5 was affected by TAC and observed shift towards slower electrophoretic mobility bands in the whole heart lysates from mice sacrificed 8 wk after TAC relative to sham hearts (**Figure 11A**). This included reduced motility of MK5.1. It is also worthy of note that band for MK5.6 was observed in the absence of EDTA in either sham or TAC hearts, but was clearly visible in the presence of EDTA, indicated this MK5 variant was phosphorylated even in healthy hearts. Furthermore, p38α/β in the phospho-T-G-F immunoreactivity was more intense in TAC heart relative to sham heart (**Figure 11B**). These results establish that MK5 is phosphorylated in healthy hearts and the phosphorylation state of MK5 is altered in hearts remodelling in response to increased afterload.

**Figure 11.**
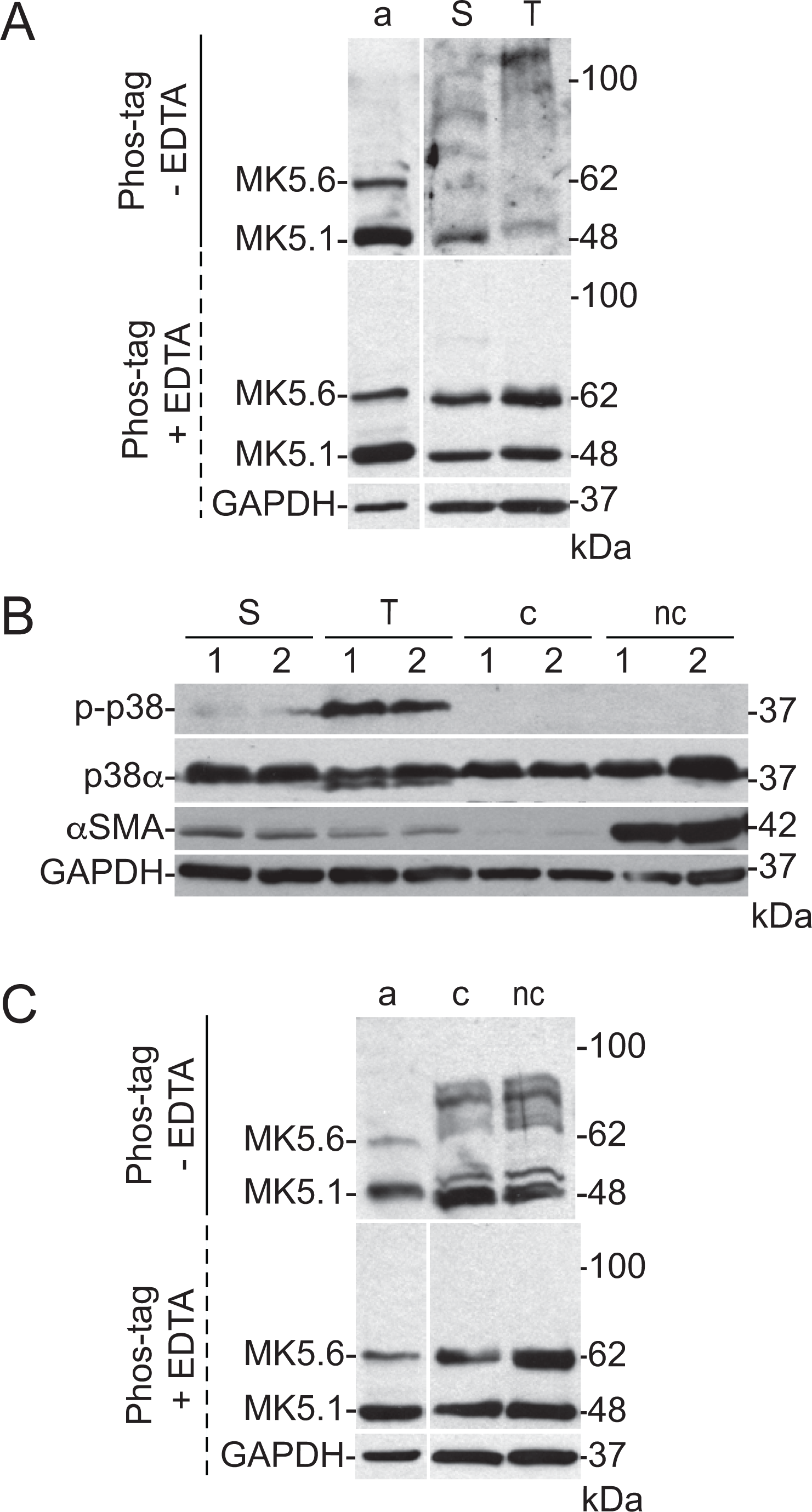
Fibroblast activation and chronic elevations in cardiac afterload alters MK5 mobility on Phos-tag SDS-PAGE. **(A)** Heart lysates (20 g/lane) from mice 8-wk after sham surgery (S) or constriction of the transverse aorta (T) were resolved on Phos-tag SDS-PAGE - EDTA and Phos-tag SDS-PAGE +EDTA followed by immunoblot assay for total MK5 immunoreactivity. Phosphatase (*λ* phosphatase followed by alkaline phosphatase) treated rat cardiac myofibroblast lysate was used as a dephosphorylated control sample on Phos-tag SDS-PAGE -EDTA (lane a). After probing for MK5 immunoreactivity, the Phos-tag SDS-PAGE +EDTA membrane was stripped and reprobed for GAPDH immunoreactivity, which was used as a loading control. **(B)** Cardiac ventricular fibroblasts were cultured on ‘compliant’ collagen-coated, hydrogel-bound (8 kPa) polystyrene plates (c) or ‘non-compliant’ uncoated standard cell culture dishes (nc), resulting in quiescent fibroblasts or myofibroblasts, respectively. Lysates from passage 2 quiescent fibroblasts and myofibroblasts as well as from 8-wk sham (S) and TAC (T) hearts (20 μg/lane) were used for immunoblot assays. Representative immunoblots for phospho-TGY-p38, p38α, *α*-smooth muscle actin (αSMA), and GAPDH are shown. Two biological replicates were loaded per group (1,2). Numbers at the right indicate the position of molecular mass markers (in kDa). **(C)** Cardiac fibroblast and myofibroblast lysates (20 g/lane) were subjected to Phos-tag SDS-PAGE ±EDTA followed by immunoblot assay for total MK5 immunoreactivity. Phosphatase (*λ* phosphatase followed by alkaline phosphatase) treated rat cardiac myofibroblast lysate was used as a dephosphorylated control sample on Phos-tag SDS-PAGE -EDTA (lane a). After probing for MK5 immunoreactivity, the Phos-tag SDS-PAGE +EDTA membrane was stripped and reprobed for GAPDH immunoreactivity, which was used as a loading control. Numbers at the right indicate the position of molecular mass markers (in kDa). Results shown are representative of three qualitatively similar experiments performed.

### MK5 is phosphorylated in both quiescent and activated cardiac fibroblasts

Fibroblasts maintained on a compliant (8 kPa) substrate remain quiescent whereas fibroblasts cultured on a rigid, non-compliant substrate become activated, a state referred to as a myofibroblast, and is associated with increased expression of α-SMA (**Figure 11B**). To examine the effect of fibroblast activation on MK5 phosphorylation, fibroblasts were isolated, cultured on rigid or compliant substrates, lysed, and the lysates resolved on Phos-tag SDS-PAGE. Surprisingly, Phos-tag SDS-PAGE revealed multiple phospho-forms of MK5 in both the quiescent fibroblasts and myofibroblasts with an increased intensity of lower mobility bands in myofibroblasts relative to quiescent fibroblasts. In both cases, no band corresponding to non-phosphorylated MK5.6 detected (**Figure 11C**). Interestingly, p38 MAPK phosphorylation was not observed in either the actively growing fibroblasts or myofibroblasts (**Figure 11B**). Therefore, phosphorylation of MK5 at least some sites is independent of fibroblast activation.

### MK5 and ERK3 form a stable complex in cardiac myofibroblasts

Studies in immortalized cell lines have identified MK5 as both a binding partner and substrate of ERK3 ^6^. Furthermore, MK5 immunoprecipitates from whole mouse heart lysates contain ERK3 immunoreactivity but not that of p38*α* ^9^. Hence, we next examined the possible interaction between MK5 and ERK3 in cardiac myofibroblasts using proximity ligation assays (PLAs). PLAs revealed ERK3-MK5 complexes in the cytoplasm but not in the nucleus (**Figure 12A**). To test the specificity of the assay, PLAs were also performed in myofibroblasts following siRNA-mediated knockdown of MK5 (MK5-kd). The efficiency of MK5 knockdown was assessed by immunoblotting as shown in Figure 12B. The MK5-ERK3 complexes were less abundant in myofibroblasts following knockdown of MK5 (**Figure 12C**), consistent with the presence of cytoplasmic ERK3-MK5 complexes in these cells. To further confirm the ERK3-MK5 interaction in myofibroblasts, we performed co-immunoprecipitation assays. MK5 immunoprecipitates contained ERK3 immunoreactivity and vice-versa (**Figure 13A**). However, whereas ERK3 immunoprecipitates contained abundant amounts of MK5.1 immunoreactivity, that of MK5.6 was less so. As previously reported from whole heart ^9^, p38*α* immunoreactivity was not detected in either MK5 or ERK3 immunoprecipitates. In serum-starved myofibroblasts, MK5 immunoreactivity was less abundant than cells were serum was re-added to the media after starvation (**Figure 13B**). In addition, in both serum-stimulated and serum-starved myofibroblasts, SB203580 reduced the abundance of MK5 immunoreactivity in ERK3 immunoprecipitates. It is also notable that the electrophoretic mobility of MK5.1 and MK5.6 enriched in ERK3 immunoprecipitates on SDS-PAGE was reduced relative to that observed in the total lysate, suggesting ERK3 may be binding a phosphorylated form of MK5. Hence, findings confirm that ERK3 and MK5 form a stable complex in myofibroblasts and suggest that p38α/β-mediated phosphorylation of MK5 may be required for either the formation or stabilization of ERK3-MK5 complexes.

**Figure 12.**
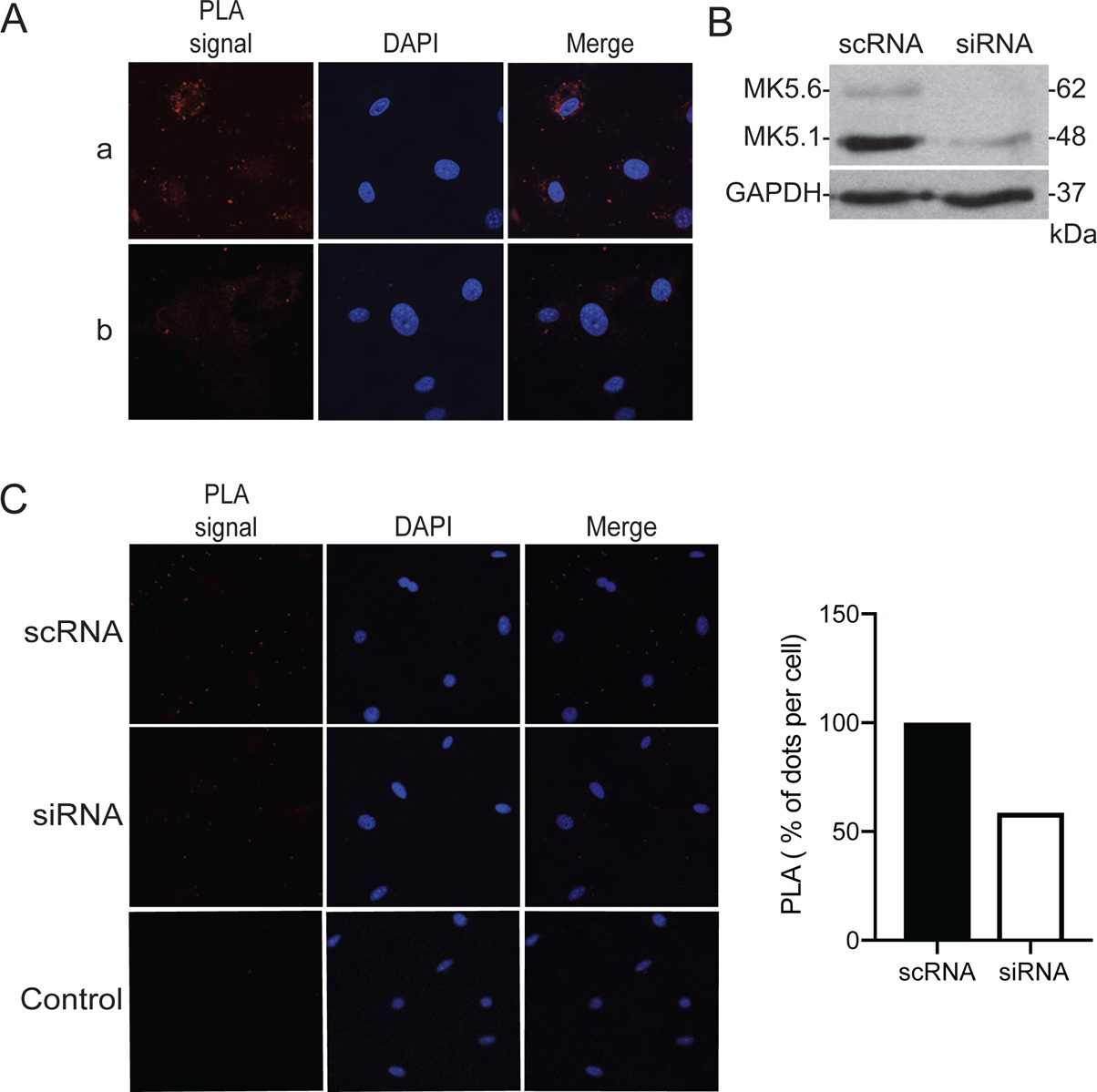
MK5 and ERK3 colocalize in the myofibroblast cytoplasm. **(A)** Passage 3 myofibroblasts were fixed, incubated with (a) or without (b) goat anti-MK5 and rabbit anti-ERK3 antibodies, and proximity revealed using DUOLink In Situ red proximity ligation assay kits and visualized by confocal fluorescence microscopy. Red dots (PLA signal) indicate sites where MK5 and ERK3 are in close proximity. Nuclei were visualized using the DNA-binding fluorescent dye, DAPI (pseudo colored blue). Merged refers to an overlay of the PLA and DAPI fluorescence. Several fields of cells were examined per sample, and the images shown are representative of three separate experiments**. (B)** Cardiac myofibroblasts isolated from MK5^+/+^ mice were transiently transfected with either siRNA for MK5 (si) or a scrambled RNA sequence (sc) as described in Methods and the abundance of MK5 immunoreactivity assessed by immunoblot assay. The intensity of both MK5.1 (48-kDa) and MK5.6 (62-kDa) bands was reduced upon MK5 knockdown. GAPDH immunoreactivity was used as a loading control. Numbers at the *right* indicate the position of molecular mass markers (in kDa). These results are qualitatively similar to those obtained from 3 separate fibroblast preparations. **(C)** Proximity ligation assays (PLA) were performed to determine if MK5 and ERK3 were in close proximity in myofibroblasts and the effect of siRNA-mediated MK5 knockdown on the PLA signal. Cardiac myofibroblasts isolated from MK5^+/+^ mice were transiently transfected with either siRNA for MK5 (si) or a scrambled RNA sequence (sc) as described in Methods. After transfection with MK5 siRNA, cells were incubated for 24 h in M199 containing 10% serum. Red dots indicate sites where MK5 and ERK3 were in close proximity (PLA signal). The number of red sports was reduced by MK5 knockdown. Nuclei were visualized using the DNA-binding fluorescent dye, DAPI (pseudo colored blue). Merged refers to an overlay of the PLA and DAPI fluorescence. Several fields of cells were examined per sample, and the images shown are representative of three separate experiments. The histogram shows the percentage of MK5-ERK3 PLA positive spots in scRNA and siRNA groups. Several fields of cells were examined, and representative images are shown from one of three performed.

**Figure 13.**
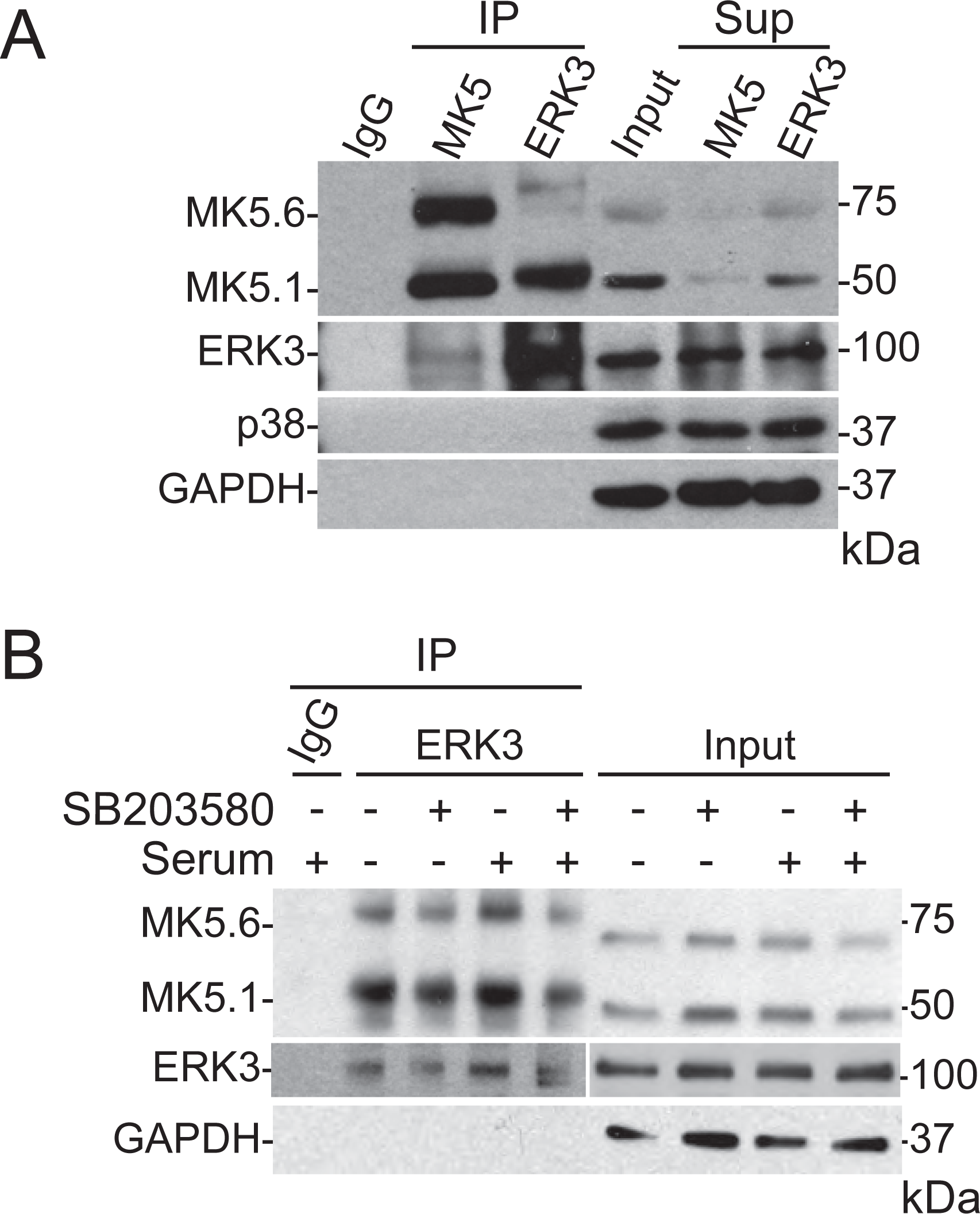
MK5 and ERK3 coimmunoprecipitate from cardiac myofibroblast lysates. **(A)** Immunoprecipitation was from lysates (500 μg/IP) prepared from passage 3 myofibroblasts. MK5 and ERK3 were immunoprecipitated using anti-MK5 antibody (PRAK A-7, # SC46667) and anti-ERK3 antibody (ERK3 5E1, # MA1-101). Immune complexes, lysate input (20 μg), and post-IP supernatants were probed for MK5, ERK3, p38*α*, and GAPDH immunoreactivity by immunoblot assay as indicated. **(B)** Immunoprecipitation was from lysates (500 μg/IP) prepared from passage 3 myofibroblasts following stimulation with 10% serum in the presence and absence of 20 μM SB203580. ERK3 was immunoprecipitated using an ERK3-specific antibody (ERK3 5E1, # MA1-101). Immune complexes and lysate input (20 μg) were probed for MK5, ERK3, and GAPDH immunoreactivity by immunoblot assay as indicated. Purified rabbit IgG was employed as a negative control during immunoprecipitation. Numbers at the *right* indicate the position of molecular mass markers (in kDa). Results shown are representative of three independent experiments.

### ERK3 stability is independent of MK5 in cardiac myofibroblasts

In cultured cell systems, ERK3 has been shown to be an unstable protein that is degraded by the ubiquitin-proteasome pathway with a half-life of 30-45 min ^33, 34^. Furthermore, the interaction with MK5 stabilizes ERK3, whereas the deletion of MK5 leads to a rapid reduction in the endogenous ERK3 levels ^6, 8^. To examine if MK5 stabilizes ERK3 in cardiac myofibroblasts, we assessed ERK3 immunoreactivity in lysates from whole heart (**Figure 14A**) and cardiac myofibroblasts (**Figure 14B**) from MK5^-/-^ and MK5^+/+^ mice. Surprisingly the abundance of ERK3 immunoreactivity was unaltered by the absence of MK5. Furthermore, confocal immunofluorescence microscopy revealed the intensity and localization of ERK3 immunoreactivity were similar in myofibroblasts from MK5^+/+^ and MK5^-/-^ mice (**Figure 14C**). As fibroblasts from MK5^-/-^ mice may have compensated for the absence of MK5, we also examined the stability of ERK3 following an acute knockdown of MK5. To this end, ERK3 immunoreactivity was also examined in myofibroblasts after a siRNA-mediated knockdown of MK5 (**Figure 15**) where we observed ERK3 immunoreactivity was unaffected by the acute knockdown of MK5. Hence, proteins in addition to MK5 may be present in cardiac fibroblasts that serve to stabilize ERK3.

**Figure 14.**
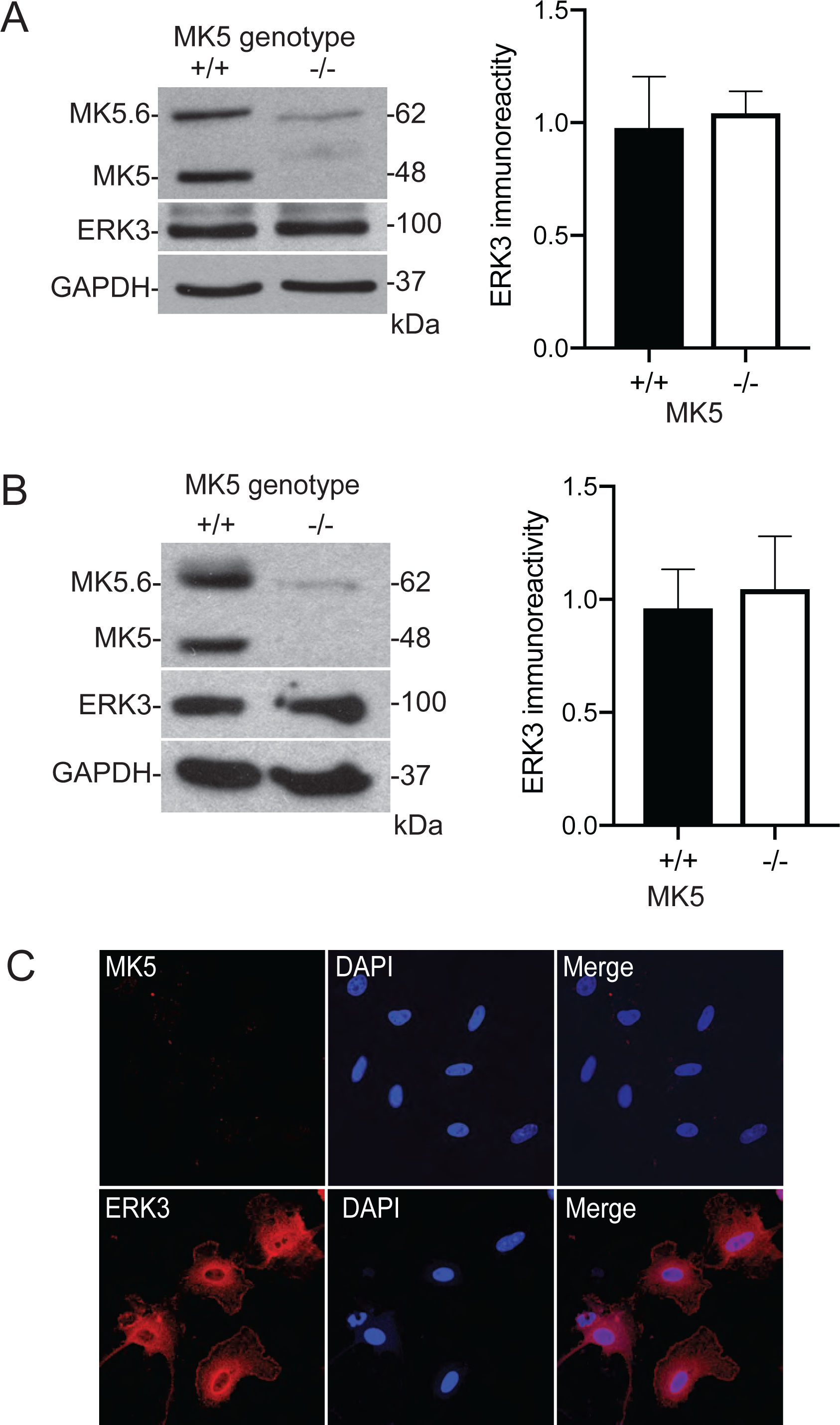
ERK3 immunoreactivity is not lost in the absence of MK5. **(A)** Heart lysates (20 μg/lane) from MK5^+/+^ and MK5^-/-^ mice were subjected to immunoblot assay. An ERK3 immunoreactive band of 100 kDa was detected in the heart lysates from both MK5^+/+^ and MK5^-/-^ mice. The intensity of the ERK3 immunoreactive band was not reduced in lysates from MK5^-/-^ mice whereas MK5.1 and MK5.6 were virtually absent. GAPDH immunoreactivity was used as a loading control. Numbers at the *right* indicate the position of molecular mass markers (in kDa). A representative immunoblot is shown in the *left*, whereas the histogram on the *right* shows the mean ± SEM for ERK3 immunoreactivity from three separate heart lysates from mice of each genotype. **(B)** Cardiac myofibroblasts were isolated from MK5^+/+^ and MK5^-/-^ mice, maintained in culture and lysed at passage 3. MK5 and ERK3 expression were assessed by immunoblot assay. An ERK3 immunoreactive band of 100 kDa was detected in the lysates from both MK5^+/+^ and MK5^-/-^ myofibroblasts. The intensity of the ERK3 immunoreactive band was not reduced in lysates from MK5^-/-^ mice whereas MK5.1 and MK5.6 were virtually absent. GAPDH immunoreactivity was used as a loading control. Numbers at the *right* indicate the position of molecular mass markers (in kDa). A representative immunoblot is shown in the *left*, whereas the *right* shows the histograms of ERK3 immunoreactivity. Data are presented as the means ± SEM of five separate cell isolations. **(C)** Passage 0 myofibroblasts from MK5^-/-^ mice fixed and decorated with antisera against MK5 or ERK3 followed by an Alexa Fluor 455-conjugated anti-rabbit secondary antibody (1:500). Immunofluorescence images showing MK5-negative staining (top panels) and ERK3-positive (bottom panels) staining in cultured cardiac fibroblasts from MK5^-/-^ mice. Nuclei were visualized by staining with DAPI (1:1000). Several fields of cells were examined, and representative images are shown.

**Figure 15.**
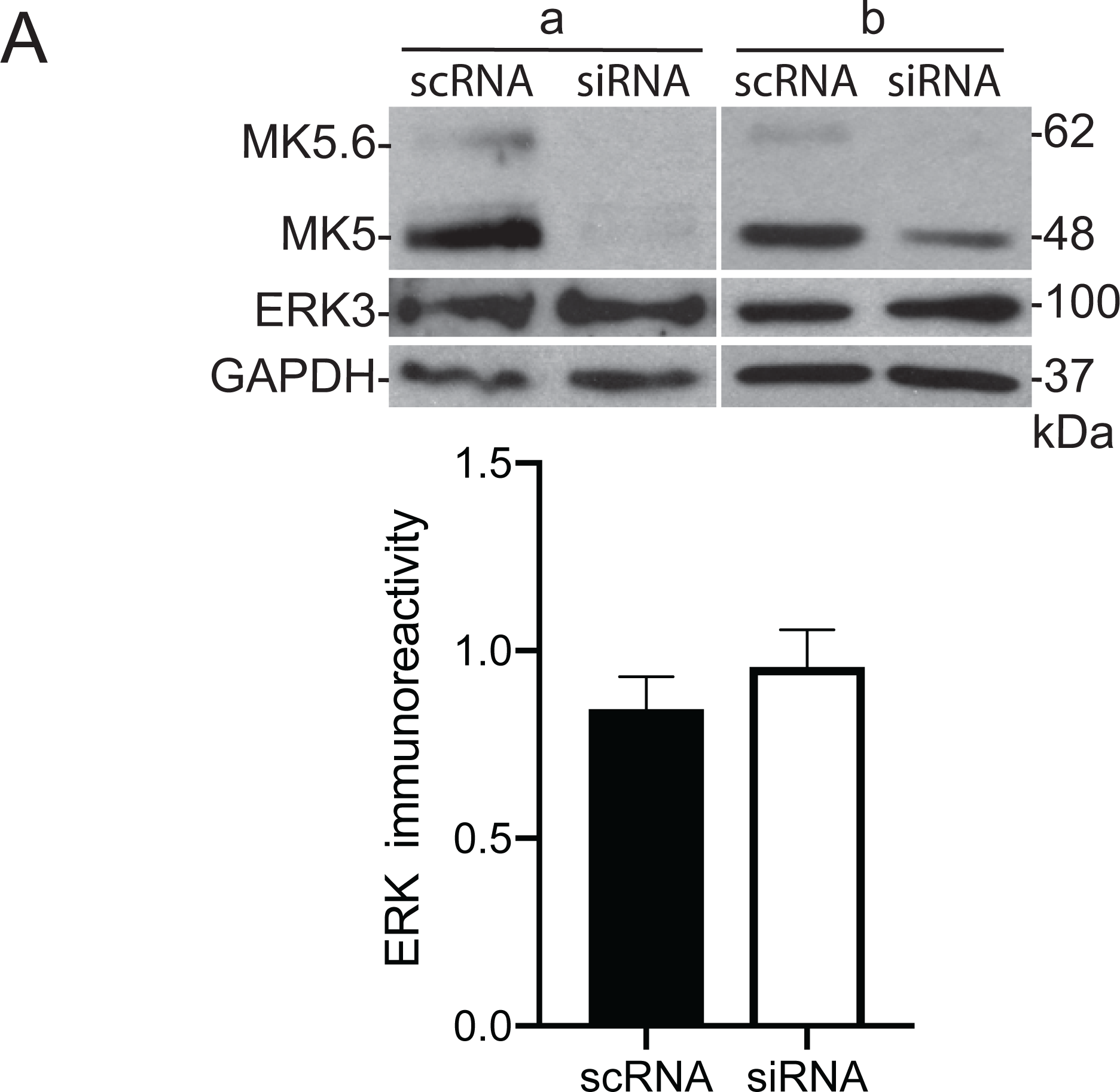
The acute knockdown of MK5 fails to alter the abundance of ERK3 immunoreactivity. **(A)** Cardiac myofibroblasts were transiently transfected with either siRNA for MK5 (si) (two different siRNA sequences: a, b) or a scrambled RNA sequence (sc) as described in Methods. The effect of MK5 knockdown on the abundance of MK5 and ERK3 immunoreactivity was assessed by immunoblot assay. The intensity of ERK3 immunoreactive band was unaffected by the acute knockdown of MK5 whereas MK5.1 and MK6.6 were virtually absent. GAPDH immunoreactivity was used as a loading control. A representative immunoblot is shown on the *left*, whereas the histogram on the *right* shows the mean ± SEM for ERK3 immunoreactivity from six determinations.

## Discussion

MK5 has been implicated in fibroblast function and cardiac remodeling ^11, 12, 35^. Putative activators of MK5 include p38*α*/*β*, ERK3, ERK4, and PKA (reviewed in ^17^). ERK3, ERK4, and MK5 have been demonstrated at the mRNA and protein levels in a variety of tissues and cell lines ^36^; however, the regulation, expression and subcellular distribution of ERK3, ERK4, and MK5 in cardiac cells remain poorly described. MK5 mRNA is ubiquitous in adult mammalian tissues including all the stages of embryogenesis and most abundant in brain, right ventricle, liver, skeletal muscle, and kidney ^4, 37, 38^. Similar to MK5, ERK3 mRNA has been detected in all the tissues and cell lines examined; however, the level of expression varies widely between cells, organs, and tissues based on their differentiation status and are of highest abundance in the brain, skeletal muscle, and gastrointestinal tract. ^36, 39, 40^. In contrast to MK5 and ERK3, ERK4 mRNA shows a more restricted expression profile ^41, 42^. The present study was to better understand the function of these protein kinases in primary cultures of adult cardiac ventricular fibroblasts and myocytes. MK5, ERK3, and ERK4 mRNA was detected in both cardiac fibroblasts and myocytes. However, whereas the copy numbers for MK5 and ERK4 mRNA were comparable between myocytes and myofibroblasts, that of ERK3 was much higher in myofibroblasts. Whereas MK5 and ERK3 immunoreactivity was detected in myofibroblasts but not myocytes the opposite was observed for ERK4. Interestingly, in spite of having comparable amounts of total MK5 mRNA, no MK5 immunoreactivity was detected in myocytes after proteasome inhibition or pro-hypertrophic stimuli. Further study revealed phosphorylation of MK5 at T182 in the activation loop (pT182-MK5) was 1) low in serum-starved fibroblasts, 2) increased sharply in the cytoplasm in response to serum, osmotic stress, AngII, TGF*β*, or H_2_O_2_, and 3) prevented in each case by pharmacological inhibition of p38*α*/*β*. In contrast, knocking down ERK3 did not prevent T182 phosphorylation but altered the cytosolic localization of pT182-MK5 immunoreactivity. In addition, inhibiting p38*α*/*β* reduced the abundance of MK5 immunoreactivity in ERK3 immunoprecipitates. However, Phos-tag SDS-PAGE revealed that MK5 is phosphorylated at multiples sites, some of which are sensitive to inhibition of p38*α*/*β*, in both serum-starved and fed quiescent fibroblasts and myofibroblasts and the electrophoretic pattern changed in response to stimuli including osmotic stress, AngII, TGF*β*, or H_2_O_2_. The phosphorylation status of MK5 in whole heart also differed between sham and 8-week TAC mice. Hence, in cardiac fibroblasts, MK5 is activated by p38*α*/*β*-mediated phosphorylation at T182 resulting in translocation of activated MK5 from the nuclear to the cytosolic compartment where at least a fraction of the activated MK5 forms a complex with ERK3. The significance of other phosphorylation sites within MK5 remains to be determined.

Five splice variants of MK5 mRNA have been identified in mice to date ^9^. An examination of the expression profile of MK5 variants in myofibroblasts and myocytes revealed only modest differences. The abundance of MK5.1, the originally described variant, was comparable between myofibroblasts and myocytes. In both cell types, the most abundant MK5 variant, following MK5.1, was MK5.2, which also was similar in abundance in these two cell types. However, none of the transcripts are translated in cardiac myocytes under conditions examined to date. Being as MK5 immunoreactivity was not detected in myocytes following inhibition of the proteasome, it’s likely the absence of MK5 at the protein level reflects the inability of these cells to translate MK5 mRNA and not rapid post-translational degradation of MK5. Although pro-hypertrophic agonists such as ET-1, AngII and Iso did not induce the translation of MK5 mRNA, it remains to be determined if other conditions may do so in myocytes and, if so, would MK5 be under the control of p38*α*/*β* or ERK4 in these cells.

Studies in cell lines have localized ERK3 to both the nuclear and cytoplasmic compartments^20, 22, 24, 26, 33^. ERK3 has also been detected in the endoplasmic reticulum (ER), ER-Golgi intermediate compartment (ERGIC), and Golgi apparatus ^43^. In cardiac myofibroblasts, ERK3 immunoreactivity was detected in the cytoplasmic compartment. Unlike conventional MAP kinases, the subcellular localization of ERK3 is independent of activation loop phosphorylation or enzymatic activity and unaffected by common mitogenic or stress stimuli such as serum, arsenate, or epidermal growth factors ^6, 20, 22^. Although no functional NES motif has been identified in ERK3, previous studies have demonstrated that the cytoplasmic localization of ERK3 is mediated by chromosome region maintenance 1 (CRM1)-dependent nuclear export mechanism with amino acids 297–542 in ERK3 comprising a CRM1-binding domain ^22, 44^. Alternatively, proteolytic cleavage within the carboxy-terminal of ERK3, a cell cycle-dependent event in HeLa cells, results in its translocation into the nucleus ^43^. Upon stimulation of cardiac myofibroblasts with serum, ERK3 immunoreactivity was observed at the periphery of the cells, suggesting a possible interaction with the membrane or cytoskeleton. The cytoplasmic retention of ERK3 in cardiac myofibroblasts may, at least in part, be due to cell type-specific differences in its post-translational modification.

Subcellular localization may be a determinant of ERK3 and MK5 function ^17, 36, 45^. In cell lines, both endogenous and heterologously expressed MK5 localized primarily to the nucleus. However, the presence of both nuclear export signal (NES) and nuclear localization signal (NLS) motifs enables a continuous shuttling of MK5 between the nucleus and cytoplasm ^7, 8, 21, 23, 25, 46–48^. Similarly, in cardiac myofibroblasts, MK5 immunoreactivity was located predominantly in the nucleus with only a weak signal in the cytoplasm. Phosphorylation plays a critical role in the regulation of MK5 activity and subcellular localization with threonine-182 being the major regulatory phosphorylation site ^48–51^. p38*α*/*β*, ERK3, and ERK4 phosphorylate MK5 at threonine-182 *in vitro* and *in vivo* whereas PKA has been reported to activate MK5 via phosphorylation at serine-115 ^4–6, 8, 52, 53^. In cardiac myofibroblasts, serum stimulation increased pT182-MK5 immunoreactivity in the cytoplasm, where it appeared to localize to the cytoskeleton and pseudopodia. Similarly, in response to sorbitol, AngII, TGFβ, or H_2_O_2_, the overall abundance of pT182-MK5 immunoreactivity increased and localized primarily to the cytoplasm in myofibroblasts. Interestingly, the distribution of pT182-MK5 immunoreactivity within the cytoplasm depended on the nature of the stimuli. In serum and H_2_O_2_-stimulated myofibroblasts, pT182-MK5 immunoreactivity was concentrated in the perinuclear region. On the other hand, pT182-MK5 immunoreactivity was associated with the cytoskeleton in response to AngII or TGFβ and showed a diffuse cytoplasmic distribution following sorbitol-treatment. However, these studies examined the effects of a 2 h exposure to stimuli and hence may differ in the time-course of their effects on the subcellular localization of pT182-MK5 immunoreactivity. In each case, inhibition of p38*α*/*β* activity with SB203580 attenuated both the increase in intensity and redistribution of pT182-MK5 immunoreactivity, suggesting activation of p38*α*/*β* MAPK may be necessary for the translocation of MK5. When co-expressed, the cellular location of the p38α-MK5 complex was solely nuclear while the p38β-MK5 complex resides in the cytoplasm, suggesting that p38α and p38β may have distinct effects on the subcellular localization of MK5^54^. Additionally, expression studies in HEK293 cells revealed that the variants MK5.1, MK5.2 and MK5.3 are in the nucleus and upon the activation of the p38 pathway, those variants relocated to the cytoplasm. The two variants lacking exons 2 – 6 inclusive (MK5.4 and MK5.5) are exclusively in the cytoplasm. However, upon p38 activation, small amounts of MK5.4 and MK5.5 were detected in the nucleus ^9^. Hence, it seems that, in cardiac fibroblasts, activated p38*α*/*β* can activate MK5 through direct phosphorylation at threonine-182 in its activation loop or bind directly to MK5 at its p38 docking site, both of which promote the subcellular translocation of MK5 effectively. On the other hand, direct binding of ERK3 and ERK4 to MK5 facilitates the nuclear export of MK5 ^7, 48, 49, 51^. In myofibroblasts, ERK3 knockdown resulted in a diffuse cytosolic distribution of pThr182-MK5 immunoreactivity but failed to attenuate the increase in intensity evoked by serum stimulation. This observation suggests that ERK3 may be responsible for targeting pThr182-MK5 to specific cytosolic sites in cardiac fibroblasts. Interestingly, the dual-specificity phosphatase Cdc14A has also been implicated in the cytoplasmic export of MK5 ^55^. Further work will be necessary to understand the role of other phosphorylation sites and interacting partners in the activation and subcellular localization of MK5 variants in cardiac fibroblasts.

In addition to threonine-182 in the MK5 activation loop, phosphorylation of MK5 at threonine-142, serine-93, serine-115, serine-212, serine-472 as well as tyrosine-188 and tyrosine-216 has been shown ^4, 25, 56, 57^. Bioinformatic analysis predicted a large number of phosphorylation sites within MK5 **(Table 2, Supplemental Tables 1 and 2)**. Phos-tag SDS-PAGE revealed multiple slower migrating bands of MK5 immunoreactivity in actively dividing myofibroblasts, which were sensitive to phosphatase treatment or the presence of EDTA (chelation of Ca^2+^) in the Phos-tag gels. These controls confirm the slower migrating bands as resulting from phosphorylation. Phos-tag SDS-PAGE revealed the presence of the slower migrating forms of MK5 immunoreactivity even in serum-deprived myofibroblasts, whereas treatment of serum-starved myofibroblasts with serum, sorbitol, AngII, TGFβ, or H_2_O_2_ increased in the abundance of slow migrating MK5 immunoreactive bands. While various treatments reduced the MK5 mobility on Phos-tag SDS-PAGE, stimulus-specific differences in the profiles were observed. Inhibition of p38α/β reduced the abundance of lower-mobility forms of MK5 to some extent but failed to abolish the mobility shift, implicating other protein kinases and phosphorylation sites in cardiac myofibroblasts. Furthermore, when compared with MK5.1, the substantial decrease in unphosphorylated MK5.6 on Phos-tag SDS-PAGE under almost all conditions examined indicates a high stoichiometry of MK5.6 phosphorylation in cardiac fibroblasts.

Phos-tag SDS-PAGE demonstrated MK5 was phosphorylated in serum-starved myofibroblasts where pT182-MK5 immunoreactivity was virtually absent. Changes in mobility observed following SB203580 treatment in serum-starved myofibroblasts indicates phosphorylation by p38*α*/*β* at sites other than threonine-182 that may be less rapidly dephosphorylated. Reducing the expression of MK5 (MK5^-/-^ or MK5-kd) in cardiac fibroblasts affects both the abundance of transcripts for numerous proteins involved in ECM homeostasis and the secretion of type 1 collagen in a manner that is dependent upon both the expression level of MK5 and the activation status of these cells ^12^. In this study, we used Phos-tag SDS-PAGE to compare the phosphorylation of MK5 in actively quiescent fibroblasts (grown on a compliant substrate) and myofibroblasts (grown on a rigid substrate). Interestingly, MK5 was phosphorylated in both fibroblasts and myofibroblasts, but the migration patterns differed, indicating MK5 phosphorylation varies depending on the activation status of the fibroblasts.

Although ERK3 displays high sequence similarity with the conventional MAPK ERK1, ERK3 is structurally distinct from its conventional counterpart. In addition, ERK3 possesses an activation motif that has only one serine as the phosphoacceptor (S189 ^58, 59^. MK5 and the steroid receptor coactivator 2 (SRC-2) are well studied interacting partners/substrates of ERK3 ^6, 8, 60^. Similarly, ERK3 and ERK4 well-characterized interacting partners of MK5 ^6, 7^. MK5 binds ERK3 and ERK4 via an unusual motif located in their C-terminals called D domain and FRIEDE motif, respectively ^49^. In the case of ERK3, phosphorylation at serine-189, but not its actual catalytic activity, is necessary for its binding to MK5 ^48, 61, 62^. We observed, co-immunoprecipitation of ERK3 and MK5 from fibroblasts suggesting they form a relatively stable complex. Furthermore, proximity ligation assays indicated ERK3 and MK5 were in close proximity only in the cytosolic compartment. Interestingly, the present study revealed that, in serum-stimulated myofibroblasts, SB203580 reduced the abundance of MK5 immunoreactivity in ERK3 immunoprecipitates. This observation suggests that p38α/β-mediated MK5 nuclear translocation of MK5 may facilitate the formation of ERK3-MK5 in cardiac fibroblasts.

MK5 haplodeficiency reduced the increase in collagen 1-*α*_1_ mRNA and hypertrophy induced by a chronic increase in cardiac afterload and MK5-deficiency alters fibroblast function, suggesting that MK5 signaling plays an important role in cardiac remodeling ^10–12^. The present study employed Phos-tag SDS-PAGE to determine if MK5 phosphorylation was altered in response to increased afterload. The transverse aortae were constricted (TAC) in MK5^+/+^ mice to induce a chronic increase in afterload and mice were sacrificed 8 weeks later. Phos-tag SDS-PAGE revealed MK5 was phosphorylated in both sham and TAC hearts eight-weeks post-surgery. However, a substantial decrease in the unphosphorylated form of MK5.1 and MK5.6 was observed in TAC hearts compared to sham hearts, suggesting that the MK5 phosphorylation was altered when the myocardium was exposed to hemodynamic overload. However, the upstream kinase(s) that activate MK5 remain controversial ^17^. Although p38α/β, ERK3 and ERK4 have been proposed as activators of MK5, only ERK3 immunoreactivity is detected in MK5 immunoprecipitates from TAC and sham hearts ^9^ and cardiac fibroblast lysates, suggesting they interact to form a stable complex. Furthermore, in the heart, endogenous MK5 or ERK3 cannot be pulled down by GST-ERK3 or GST-MK5, respectively, suggesting neither exists in a monomeric state in cardiac cells ^9^. Based upon the findings of the present study, we propose that in cardiac fibroblasts, stimuli such as stress-activated p38*α*/*β*, which enter the nucleus and phosphorylated MK5 at threonine-182 causing MK5 to relocate to the cytosol where it can form a complex with ERK3. As MK5 and ERK3 immunoreactivity was detected in fibroblasts, but not myocytes, and stimulating myocytes with pro-hypertrophic hormones did not induce the appearance of MK5 immunoreactivity, the attenuated in hypertrophy observed in MK5-haplodeficient mice likely reflects a role for MK5 in fibroblast-myocyte communication and ERK4 may function in a novel signaling pathway in myocytes that does not involve MK5.

In spite of having a close relation with ERK4, ERK4 is stable whereas ERK3 has been shown to be a highly unstable protein with a half-life of about 1 h in immortalized cell lines ^7, 33, 51^. These studies found that MK5 stabilized ERK3 ^8^. siRNA-mediated knockdown of MK5 drastically reduced the level of endogenous ERK3. Similarly, ERK3 levels were changed during embryogenesis in MK5 null mice. These reports strengthen the assumption that MK5 may act as a chaperone of ERK3 and stabilize ERK3 by direct interaction. However, in contrast to other cell systems, ERK3 was not destabilized by the absence of MK5 in cardiac myofibroblasts, suggesting the presence of an as-of-yet identified binding partner in cardiac fibroblasts that stabilizes the ERK3 protein. Chen *et.al.* recently reported an oncogenic BRaf-mediated upregulation of ERK3 expression where BRaf increased the abundance of ERK3 mRNA and stabilized ERK3 protein ^63^. Further studies are needed to elucidate the mechanism of MK5-independent stabilization of ERK3 in cardiac myofibroblasts.

## Conclusions

This study provides important insights into the fundamental understanding of the expression, cell localization and molecular interaction of MK5, ERK3, and ERK4 in cardiac fibroblasts and myocytes. MK5 is transcribed but not translated in myocytes: MK5 immunoreactivity was not rescued by inhibition of the proteasome and translation of MK5 mRNA was not induced in response to the pro-hypertrophic agonists. Conditions examined, phosphorylation of MK5 at threonine-182 in the activation loop was attenuated by inhibition of p38*α*/*β* as was the ability of MK5 to form a complex with ERK3. In contrast, Phos-tag SDS-PAGE demonstrated that MK5 was highly phosphorylated, in the absence or presence of stimuli, and even in the absence of threonine-182 phosphorylation. Further work is needed to determine the significance or MK5 phosphorylation at sites other than threonine-182.

## Funding

This work is supported by grants from the Heart and Stroke Foundation of Canada (Grant Numbers G-14-0006060 and G-18-0022227).

## Acknowledgements

We thank Dr. Robert Parent for animal care and breeding. The Authors would also like to thank Cindy Sutherland and Mike Walsh for their assistance in setting up the Phos-tag SDS-PAGE system in the Allen lab.

## Notes

### Competing Interest Statement

The authors have declared no competing interest.

