## Supplemental Table 1 for "Expression, subcellular localization, and phosphorylation of MK5 in adult cardiac ventricular fibroblasts"

**Supplemental Table 1. Predicted serine/threonine phosphorylation sites and their cognate protein kinases in mouse MK5.1.**

| position | residue | kinase | Peptide sequence | score | cutoff |
| --- | --- | --- | --- | --- | --- |
| 2 | S | VRK | ******MSEDSDMEK | 440.814 | 303.187 |
| 2 | S | CK2 | ******MSEDSDMEK | 4.486 | 2.665 |
| 2 | S | CAMKK | ******MSEDSDMEK | 3.617 | 3.05 |
| 2 | S | Haspin | ******MSEDSDMEK | 12.182 | 0.998 |
| 2 | S | WEE | ******MSEDSDMEK | 3.527 | 2.306 |
| 2 | S | CRIK | ******MSEDSDMEK | 2.315 | 0.919 |
| 2 | S | PRKG2 | ******MSEDSDMEK | 62.727 | 55.934 |
| 2 | S | PHKG1 | ******MSEDSDMEK | 2.575 | 2.484 |
| 2 | S | VRK2 | ******MSEDSDMEK | 68.436 | 48.397 |
| 2 | S | CK2a1 | ******MSEDSDMEK | 6.202 | 4.483 |
| 2 | S | MSN | ******MSEDSDMEK | 6.214 | 3.852 |
| 2 | S | LIMK | ******MSEDSDMEK | 19.114 | -0.331 |
| 2 | S | CAMKK-Meta | ******MSEDSDMEK | 0.001 | 0.001 |
| 2 | S | Myt1 | ******MSEDSDMEK | 3.971 | 2.486 |
| 2 | S | CIT | ******MSEDSDMEK | 2.315 | 0.919 |
| 2 | S | DMPK | ******MSEDSDMEK | 6.128 | 3.509 |
| 2 | S | GRK1 | ******MSEDSDMEK | 13.353 | 10.931 |
| 2 | S | STK39 | ******MSEDSDMEK | 88.029 | 86.489 |
| 2 | S | MAP4K3 | ******MSEDSDMEK | 5.705 | 3.801 |
| 2 | S | STE20 | ******MSEDSDMEK | 2.663 | 2.462 |
| 2 | S | LIMK1 | ******MSEDSDMEK | 19.114 | -0.331 |
| 2 | S | TRPM6 | ******MSEDSDMEK | 4.92 | 1.899 |
| 2 | S | CAMKK1 | ******MSEDSDMEK | 2.042 | 1.714 |
| 2 | S | PKMYT1 | ******MSEDSDMEK | 3.971 | 2.486 |
| 5 | S | CK1 | ***MSEDSDMEKAIK | 7.94 | 6.306 |
| 5 | S | CK2 | ***MSEDSDMEKAIK | 7.918 | 2.665 |
| 5 | S | CAMKK | ***MSEDSDMEKAIK | 3.174 | 3.05 |
| 5 | S | Haspin | ***MSEDSDMEKAIK | 10.556 | 0.998 |
| 5 | S | WEE | ***MSEDSDMEKAIK | 4.384 | 2.306 |
| 5 | S | CRIK | ***MSEDSDMEKAIK | 2.438 | 0.919 |
| 5 | S | DRAK | ***MSEDSDMEKAIK | 5.134 | 4.561 |
| 5 | S | PIM3 | ***MSEDSDMEKAIK | 5.708 | 5.258 |
| 5 | S | CK1 | ***MSEDSDMEKAIK | 11.28 | 10.881 |
| 5 | S | VRK1 | ***MSEDSDMEKAIK | 0.399 | 0.394 |
| 5 | S | CK2a1 | ***MSEDSDMEKAIK | 7.121 | 4.483 |
| 5 | S | CSNK2A2 | ***MSEDSDMEKAIK | 0.001 | 0.001 |
| 5 | S | KHS | ***MSEDSDMEKAIK | 0.789 | 0.771 |
| 5 | S | LIMK | ***MSEDSDMEKAIK | 15.023 | -0.331 |
| 5 | S | STKR1 | ***MSEDSDMEKAIK | 12.835 | 11.666 |
| 5 | S | CAMKK-Meta | ***MSEDSDMEKAIK | 0.001 | 0.001 |
| 5 | S | PLK1 | ***MSEDSDMEKAIK | 13.666 | 13.525 |
| 5 | S | TLK1 | ***MSEDSDMEKAIK | 3.178 | 3.166 |
| 5 | S | TLK2 | ***MSEDSDMEKAIK | 2.463 | 2.396 |
| 5 | S | Myt1 | ***MSEDSDMEKAIK | 4.558 | 2.486 |
| 5 | S | CIT | ***MSEDSDMEKAIK | 2.438 | 0.919 |
| 5 | S | DMPK | ***MSEDSDMEKAIK | 4.563 | 3.509 |
| 5 | S | GRK1 | ***MSEDSDMEKAIK | 13.415 | 10.931 |
| 5 | S | PRKCI | ***MSEDSDMEKAIK | 9.982 | 9.121 |
| 5 | S | CSNK1A1 | ***MSEDSDMEKAIK | 26.053 | 24.196 |
| 5 | S | LIMK1 | ***MSEDSDMEKAIK | 15.023 | -0.331 |
| 5 | S | TRPM6 | ***MSEDSDMEKAIK | 5.934 | 1.899 |
| 5 | S | CAMKK2 | ***MSEDSDMEKAIK | 22.268 | 20.956 |
| 5 | S | PLK1 | ***MSEDSDMEKAIK | 13.067 | 12.929 |
| 5 | S | ULK2 | ***MSEDSDMEKAIK | 4.264 | 4.057 |
| 5 | S | PKMYT1 | ***MSEDSDMEKAIK | 4.558 | 2.486 |
| 5 | S | SrcA | ***MSEDSDMEKAIK | 3.726 | 3.458 |
| 5 | S | SRC | ***MSEDSDMEKAIK | 4.184 | 3.708 |
| 14 | T | CAMK1 | MEKAIKETSILEEYS | 12.68 | 11.583 |
| 14 | T | DAPK | MEKAIKETSILEEYS | 12.019 | 9.755 |
| 14 | T | PHK | MEKAIKETSILEEYS | 3.498 | 3.018 |
| 14 | T | VRK | MEKAIKETSILEEYS | 354.141 | 303.187 |
| 14 | T | OCDC7 | MEKAIKETSILEEYS | 79.251 | 77.782 |
| 14 | T | Haspin | MEKAIKETSILEEYS | 4.42 | 0.998 |
| 14 | T | TOPK | MEKAIKETSILEEYS | 3.517 | 3.19 |
| 14 | T | WEE | MEKAIKETSILEEYS | 3.001 | 2.306 |
| 14 | T | CRIK | MEKAIKETSILEEYS | 3.879 | 0.919 |
| 14 | T | DAPK | MEKAIKETSILEEYS | 21.86 | 17.52 |
| 14 | T | LIMK | MEKAIKETSILEEYS | 6.56 | -0.331 |
| 14 | T | BUB1 | MEKAIKETSILEEYS | 1.97E-5 | 0E0 |
| 14 | T | CDC7 | MEKAIKETSILEEYS | 79.251 | 77.782 |
| 14 | T | HASPIN | MEKAIKETSILEEYS | 4.42 | 0.998 |
| 14 | T | PBK | MEKAIKETSILEEYS | 3.517 | 3.19 |
| 14 | T | Myt1 | MEKAIKETSILEEYS | 4.103 | 2.486 |
| 14 | T | CIT | MEKAIKETSILEEYS | 3.879 | 0.919 |
| 14 | T | DMPK | MEKAIKETSILEEYS | 4.398 | 3.509 |
| 14 | T | STK39 | MEKAIKETSILEEYS | 88.576 | 86.489 |
| 14 | T | LIMK1 | MEKAIKETSILEEYS | 6.56 | -0.331 |
| 14 | T | TOR2 | MEKAIKETSILEEYS | 2.272 | 2.201 |
| 14 | T | CAMKK1 | MEKAIKETSILEEYS | 1.819 | 1.714 |
| 14 | T | ULK2 | MEKAIKETSILEEYS | 5.436 | 4.057 |
| 14 | T | PKMYT1 | MEKAIKETSILEEYS | 4.103 | 2.486 |
| 15 | S | STE | EKAIKETSILEEYSI | 7.375 | 6.594 |
| 15 | S | RSK | EKAIKETSILEEYSI | 11.083 | 10.901 |
| 15 | S | CAMK1 | EKAIKETSILEEYSI | 12.393 | 11.583 |
| 15 | S | DAPK | EKAIKETSILEEYSI | 11.695 | 9.755 |
| 15 | S | CK2 | EKAIKETSILEEYSI | 3.098 | 2.665 |
| 15 | S | Bud32 | EKAIKETSILEEYSI | 3.224 | 2.937 |
| 15 | S | Haspin | EKAIKETSILEEYSI | 3.332 | 0.998 |
| 15 | S | PLK | EKAIKETSILEEYSI | 17.202 | 14.476 |
| 15 | S | TLK | EKAIKETSILEEYSI | 3.282 | 2.277 |
| 15 | S | WEE | EKAIKETSILEEYSI | 2.985 | 2.306 |
| 15 | S | TK | EKAIKETSILEEYSI | 73.15 | 47.151 |
| 15 | S | CRIK | EKAIKETSILEEYSI | 4.253 | 0.919 |
| 15 | S | CAMK2A | EKAIKETSILEEYSI | 8.134 | 7.322 |
| 15 | S | CAMK2B | EKAIKETSILEEYSI | 159.516 | 147.247 |
| 15 | S | NuaK | EKAIKETSILEEYSI | 6.051 | 5.434 |
| 15 | S | DAPK | EKAIKETSILEEYSI | 18.004 | 17.52 |
| 15 | S | DRAK | EKAIKETSILEEYSI | 5.701 | 4.561 |
| 15 | S | PHKG1 | EKAIKETSILEEYSI | 2.973 | 2.484 |
| 15 | S | VRK2 | EKAIKETSILEEYSI | 52.434 | 48.397 |
| 15 | S | CSNK2A2 | EKAIKETSILEEYSI | 0.001 | 0.001 |
| 15 | S | CLK2 | EKAIKETSILEEYSI | 0.006 | 0.006 |
| 15 | S | LIMK | EKAIKETSILEEYSI | 6.028 | -0.331 |
| 15 | S | TP53RK | EKAIKETSILEEYSI | 3.224 | 2.937 |
| 15 | S | CAMKK-Meta | EKAIKETSILEEYSI | 0.001 | 0.001 |
| 15 | S | HASPIN | EKAIKETSILEEYSI | 3.332 | 0.998 |
| 15 | S | NEK11 | EKAIKETSILEEYSI | 4.306 | 3.839 |
| 15 | S | PLK1 | EKAIKETSILEEYSI | 16.702 | 13.525 |
| 15 | S | TLK1 | EKAIKETSILEEYSI | 3.819 | 3.166 |
| 15 | S | Myt1 | EKAIKETSILEEYSI | 3.006 | 2.486 |
| 15 | S | Lmr | EKAIKETSILEEYSI | 6.066 | 5.599 |
| 15 | S | Src | EKAIKETSILEEYSI | 4.892 | 4.017 |
| 15 | S | Tec | EKAIKETSILEEYSI | 3.713 | 3.05 |
| 15 | S | CIT | EKAIKETSILEEYSI | 4.253 | 0.919 |
| 15 | S | DMPK | EKAIKETSILEEYSI | 5.129 | 3.509 |
| 15 | S | PRKCB | EKAIKETSILEEYSI | 5.748 | 4.392 |
| 15 | S | NUAK1 | EKAIKETSILEEYSI | 6.051 | 5.434 |
| 15 | S | STK17B | EKAIKETSILEEYSI | 3.371 | 3.169 |
| 15 | S | MAP2K1 | EKAIKETSILEEYSI | 95.608 | 91.159 |
| 15 | S | LIMK1 | EKAIKETSILEEYSI | 6.028 | -0.331 |
| 15 | S | TRPM6 | EKAIKETSILEEYSI | 3.203 | 1.899 |
| 15 | S | MTOR | EKAIKETSILEEYSI | 14.964 | 10.962 |
| 15 | S | NEK11 | EKAIKETSILEEYSI | 4.306 | 3.839 |
| 15 | S | PLK1 | EKAIKETSILEEYSI | 15.681 | 12.929 |
| 15 | S | ULK2 | EKAIKETSILEEYSI | 5.466 | 4.057 |
| 15 | S | PKMYT1 | EKAIKETSILEEYSI | 3.006 | 2.486 |
| 15 | S | LMTK2 | EKAIKETSILEEYSI | 6.066 | 5.599 |
| 15 | S | SrcA | EKAIKETSILEEYSI | 4.084 | 3.458 |
| 15 | S | BTK | EKAIKETSILEEYSI | 3.713 | 3.05 |
| 21 | S | VRK | TSILEEYSINWTQKL | 326.999 | 303.187 |
| 21 | S | IRAK | TSILEEYSINWTQKL | 3.395 | 3.391 |
| 21 | S | Haspin | TSILEEYSINWTQKL | 3.28 | 0.998 |
| 21 | S | PLK | TSILEEYSINWTQKL | 14.905 | 14.476 |
| 21 | S | CRIK | TSILEEYSINWTQKL | 5.058 | 0.919 |
| 21 | S | CAMK2B | TSILEEYSINWTQKL | 153.793 | 147.247 |
| 21 | S | CAMK2G | TSILEEYSINWTQKL | 4.026 | 3.87 |
| 21 | S | MYLK2 | TSILEEYSINWTQKL | 4.712 | 4.132 |
| 21 | S | RK2 | TSILEEYSINWTQKL | 69.591 | 48.397 |
| 21 | S | LIMK | TSILEEYSINWTQKL | 2.677 | -0.331 |
| 21 | S | BUB1 | TSILEEYSINWTQKL | 3.5E-6 | 0E0 |
| 21 | S | PLK1 | TSILEEYSINWTQKL | 14.138 | 13.525 |
| 21 | S | PLK2 | TSILEEYSINWTQKL | 22.341 | 19.736 |
| 21 | S | TLK2 | TSILEEYSINWTQKL | 2.581 | 2.396 |
| 21 | S | CIT | TSILEEYSINWTQKL | 5.058 | 0.919 |
| 21 | S | STE20 | TSILEEYSINWTQKL | 2.72 | 2.462 |
| 21 | S | LIMK1 | TSILEEYSINWTQKL | 2.677 | -0.331 |
| 21 | S | CAMKK1 | TSILEEYSINWTQKL | 2.185 | 1.714 |
| 21 | S | PLK1 | TSILEEYSINWTQKL | 13.982 | 12.929 |
| 21 | S | PLK2 | TSILEEYSINWTQKL | 19.846 | 18.362 |
| 21 | S | ULK2 | TSILEEYSINWTQKL | 4.771 | 4.057 |
| 21 | S | SrcA | TSILEEYSINWTQKL | 3.857 | 3.458 |
| 25 | T | STE7 | EEYSINWTQKLGAGI | 35.43 | 31.362 |
| 25 | T | PIKK | EEYSINWTQKLGAGI | 12.166 | 12.124 |
| 25 | T | CAMKK | EEYSINWTQKLGAGI | 3.214 | 3.05 |
| 25 | T | CRIK | EEYSINWTQKLGAGI | 2.832 | 0.919 |
| 25 | T | VRK2 | EEYSINWTQKLGAGI | 49.957 | 48.397 |
| 25 | T | KHS | EEYSINWTQKLGAGI | 0.81 | 0.771 |
| 25 | T | LIMK | EEYSINWTQKLGAGI | 0.93 | -0.331 |
| 25 | T | ATM | EEYSINWTQKLGAGI | 16.479 | 13.727 |
| 25 | T | BUB1 | EEYSINWTQKLGAGI | 2.16E-5 | 0E0 |
| 25 | T | CAMKK-Meta | EEYSINWTQKLGAGI | 0.001 | 0.001 |
| 25 | T | Lmr | EEYSINWTQKLGAGI | 5.782 | 5.599 |
| 25 | T | Src | EEYSINWTQKLGAGI | 4.762 | 4.017 |
| 25 | T | CIT | EEYSINWTQKLGAGI | 2.832 | 0.919 |
| 25 | T | MAP2K3 | EEYSINWTQKLGAGI | 5.68E-5 | 0E0 |
| 25 | T | LIMK1 | EEYSINWTQKLGAGI | 0.93 | -0.331 |
| 25 | T | ATM | EEYSINWTQKLGAGI | 16.479 | 13.727 |
| 25 | T | CAMKK1 | EEYSINWTQKLGAGI | 2.429 | 1.714 |
| 25 | T | LMTK2 | EEYSINWTQKLGAGI | 5.782 | 5.599 |
| 33 | S | MNK | QKLGAGISGPVRVCV | 19.08 | 18.111 |
| 33 | S | PRKD2 | QKLGAGISGPVRVCV | 2.364 | 2.35 |
| 33 | S | BUB1 | QKLGAGISGPVRVCV | 4.67E-5 | 0E0 |
| 33 | S | Src | QKLGAGISGPVRVCV | 4.819 | 4.017 |
| 33 | S | ARAF | QKLGAGISGPVRVCV | 3.882 | 3.868 |
| 33 | S | PDK4 | QKLGAGISGPVRVCV | 3.201 | 2.463 |
| 43 | S | CK2 | VRVCVKKSTQERFAL | 2.729 | 2.665 |
| 43 | S | MLKA | VRVCVKKSTQERFAL | 1.445 | 1.396 |
| 43 | S | CSNK2A2 | VRVCVKKSTQERFAL | 0.001 | 0.001 |
| 43 | S | AURKC | VRVCVKKSTQERFAL | 23.429 | 18.34 |
| 43 | S | BUB1 | VRVCVKKSTQERFAL | 4.03E-6 | 0E0 |
| 43 | S | MKNK1 | VRVCVKKSTQERFAL | 10.583 | 10.57 |
| 43 | S | MAP3K10 | VRVCVKKSTQERFAL | 3.673 | 3.409 |
| 44 | T | KHS | RVCVKKSTQERFALK | 0.833 | 0.771 |
| 44 | T | BUB1 | RVCVKKSTQERFALK | 7.82E-5 | 0E0 |
| 71 | T | RAF | RLHMMCATHPNIVQI | 2.697 | 2.477 |
| 71 | T | NDR | RLHMMCATHPNIVQI | 0.492 | 0.407 |
| 71 | T | BUB1 | RLHMMCATHPNIVQI | 7.21E-6 | 0E0 |
| 71 | T | STK38 | RLHMMCATHPNIVQI | 1.57 | 0.393 |
| 71 | T | STK39 | RLHMMCATHPNIVQI | 90.568 | 86.489 |
| 71 | T | STE20 | RLHMMCATHPNIVQI | 2.536 | 2.462 |
| 85 | S | BUB1 | IIEVFANSVQFPHES | 3.35E-5 | 0E0 |
| 85 | S | NEK2 | IIEVFANSVQFPHES | 3.099 | 2.335 |
| 85 | S | MAP2K3 | IIEVFANSVQFPHES | 4.22E-5 | 0E0 |
| 85 | S | NEK2 | IIEVFANSVQFPHES | 3.099 | 2.335 |
| 92 | S | VRK | SVQFPHESSPRARLL | 329.601 | 303.187 |
| 92 | S | BRSK | SVQFPHESSPRARLL | 1.413 | 1.335 |
| 92 | S | VRK2 | SVQFPHESSPRARLL | 51.781 | 48.397 |
| 92 | S | NEK11 | SVQFPHESSPRARLL | 4.327 | 3.839 |
| 92 | S | GRK3 | SVQFPHESSPRARLL | 13.399 | 13.381 |
| 92 | S | NEK11 | SVQFPHESSPRARLL | 4.327 | 3.839 |
| 93 | S | IRAK | VQFPHESSPRARLLI | 3.392 | 3.391 |
| 93 | S | KIS | VQFPHESSPRARLLI | 3.143 | 2.45 |
| 93 | S | PKCa | VQFPHESSPRARLLI | 9.042 | 9.03 |
| 93 | S | CCRK | VQFPHESSPRARLLI | 3.342 | 3.11 |
| 93 | S | CRK7 | VQFPHESSPRARLLI | 4.99 | 4.506 |
| 93 | S | ERK1 | VQFPHESSPRARLLI | 11.477 | 11.368 |
| 93 | S | nmo | VQFPHESSPRARLLI | 0.326 | 0.259 |
| 93 | S | IRAK1 | VQFPHESSPRARLLI | 91.854 | 89.042 |
| 93 | S | UHMK1 | VQFPHESSPRARLLI | 3.143 | 2.45 |
| 93 | S | TLK2 | VQFPHESSPRARLLI | 3.156 | 2.396 |
| 93 | S | PRKCB | VQFPHESSPRARLLI | 5.704 | 4.392 |
| 93 | S | CDK20 | VQFPHESSPRARLLI | 3.342 | 3.11 |
| 93 | S | CDK3 | VQFPHESSPRARLLI | 10.486 | 9.024 |
| 93 | S | CDK12 | VQFPHESSPRARLLI | 4.99 | 4.506 |
| 93 | S | HIPK1 | VQFPHESSPRARLLI | 3.734 | 3.685 |
| 93 | S | ERK2 (MAPK1) | VQFPHESSPRARLLI | 15.154 | 14.663 |
| 93 | S | JNK3 (MAPK10) | VQFPHESSPRARLLI | 0.024 | 0.023 |
| 93 | S | NLK | VQFPHESSPRARLLI | 0.326 | 0.259 |
| 93 | S | STE20 | VQFPHESSPRARLLI | 2.645 | 2.462 |
| 115 | S | PKA | GELFHRISQHRHFTE | 23.157 | 22.884 |
| 115 | S | Aur | GELFHRISQHRHFTE | 9.145 | 8.819 |
| 115 | S | PKACA | GELFHRISQHRHFTE | 4.143 | 3.21 |
| 115 | S | PKCi | GELFHRISQHRHFTE | 6.096 | 3.394 |
| 115 | S | MSK | GELFHRISQHRHFTE | 2.779 | 2.665 |
| 115 | S | MLKA | GELFHRISQHRHFTE | 1.435 | 1.396 |
| 115 | S | PHKG1 | GELFHRISQHRHFTE | 2.552 | 2.484 |
| 115 | S | PRKD1 | GELFHRISQHRHFTE | 6.981 | 6.813 |
| 115 | S | ILK | GELFHRISQHRHFTE | 23.462 | 21.936 |
| 115 | S | Lmr | GELFHRISQHRHFTE | 5.724 | 5.599 |
| 115 | S | PRKCA | GELFHRISQHRHFTE | 19.585 | 19.505 |
| 115 | S | PRKCZ | GELFHRISQHRHFTE | 51.845 | 41.793 |
| 115 | S | RPS6KA5 | GELFHRISQHRHFTE | 2.673 | 2.558 |
| 115 | S | CLA4 | GELFHRISQHRHFTE | 4.288 | 3.489 |
| 115 | S | ILK | GELFHRISQHRHFTE | 23.462 | 21.936 |
| 115 | S | ULK2 | GELFHRISQHRHFTE | 4.573 | 4.057 |
| 115 | S | LMTK2 | GELFHRISQHRHFTE | 5.724 | 5.599 |
| 121 | T | AKT3 | ISQHRHFTEKQASQV | 4.415 | 4.201 |
| 121 | T | CAMK1 | ISQHRHFTEKQASQV | 45.482 | 40.948 |
| 121 | T | DAPK | ISQHRHFTEKQASQV | 17.698 | 17.52 |
| 121 | T | PIM1 | ISQHRHFTEKQASQV | 0.001 | 0.001 |
| 121 | T | PIM3 | ISQHRHFTEKQASQV | 5.573 | 5.258 |
| 121 | T | CDK7 | ISQHRHFTEKQASQV | 21.415 | 20.158 |
| 121 | T | FRAY | ISQHRHFTEKQASQV | 102.454 | 86.942 |
| 121 | T | BUB1 | ISQHRHFTEKQASQV | 1.87E-5 | 0E0 |
| 121 | T | Lmr | ISQHRHFTEKQASQV | 5.699 | 5.599 |
| 121 | T | GRK7 | ISQHRHFTEKQASQV | 1.695 | 1.646 |
| 121 | T | STK38 | ISQHRHFTEKQASQV | 0.409 | 0.393 |
| 121 | T | RPS6KA2 | ISQHRHFTEKQASQV | 12.871 | 12.185 |
| 121 | T | DAPK1 | ISQHRHFTEKQASQV | 18.418 | 12.301 |
| 121 | T | DAPK3 | ISQHRHFTEKQASQV | 9.746 | 9.069 |
| 121 | T | CDK7 | ISQHRHFTEKQASQV | 22.274 | 20.865 |
| 121 | T | LMTK2 | ISQHRHFTEKQASQV | 5.699 | 5.599 |
| 126 | S | PIKK | HFTEKQASQVTKQIA | 12.419 | 12.124 |
| 126 | S | TLK | HFTEKQASQVTKQIA | 2.993 | 2.277 |
| 126 | S | CHK1 | HFTEKQASQVTKQIA | 5.674 | 5.673 |
| 126 | S | MEK3 | HFTEKQASQVTKQIA | 0.029 | 0.025 |
| 126 | S | DNAPK | HFTEKQASQVTKQIA | 11.966 | 11.766 |
| 126 | S | SMG1 | HFTEKQASQVTKQIA | 3.926 | 3.251 |
| 126 | S | BUB1 | HFTEKQASQVTKQIA | 2.71E-5 | 0E0 |
| 126 | S | NEK2 | HFTEKQASQVTKQIA | 2.418 | 2.335 |
| 126 | S | TLK1 | HFTEKQASQVTKQIA | 4.639 | 3.166 |
| 126 | S | PRKCB | HFTEKQASQVTKQIA | 5.443 | 4.392 |
| 126 | S | BMPR1B | HFTEKQASQVTKQIA | -0.955 | -1.013 |
| 126 | S | PRKDC | HFTEKQASQVTKQIA | 11.966 | 11.766 |
| 126 | S | SMG1 | HFTEKQASQVTKQIA | 3.926 | 3.251 |
| 126 | S | NEK2 | HFTEKQASQVTKQIA | 2.418 | 2.335 |
| 129 | T | TK | EKQASQVTKQIALAL | 52.09 | 47.151 |
| 129 | T | DYRK1 | EKQASQVTKQIALAL | 22.973 | 21.744 |
| 160 | S | CAMK-Unique | NLLFKDNSLDAPVKL | 5.997 | 4.241 |
| 160 | S | STE20 | NLLFKDNSLDAPVKL | 6.885 | 6.147 |
| 160 | S | TTK | NLLFKDNSLDAPVKL | 21.481 | 21.21 |
| 160 | S | SRK2E | NLLFKDNSLDAPVKL | 4.03 | 3.558 |
| 160 | S | MELK | NLLFKDNSLDAPVKL | 214.678 | 195.594 |
| 160 | S | BUB1 | NLLFKDNSLDAPVKL | 1.04E-6 | 0E0 |
| 160 | S | TTK | NLLFKDNSLDAPVKL | 24.179 | 22.818 |
| 160 | S | MELK | NLLFKDNSLDAPVKL | 214.678 | 195.594 |
| 160 | S | STK3 | NLLFKDNSLDAPVKL | 101.89 | 73.61 |
| 160 | S | CDC5 | NLLFKDNSLDAPVKL | 278.002 | 245.053 |
| 182 | T | GRK | VDQGDLMTPQFTPYY | 3.396 | 2.938 |
| 182 | T | GSK | VDQGDLMTPQFTPYY | 10.671 | 8.674 |
| 182 | T | GTF2F1 | VDQGDLMTPQFTPYY | 3.195 | 2.841 |
| 182 | T | CAMKK | VDQGDLMTPQFTPYY | 3.986 | 3.05 |
| 182 | T | PKH2 | VDQGDLMTPQFTPYY | 5.458 | 3.289 |
| 182 | T | PRKD3 | VDQGDLMTPQFTPYY | 3.176 | 2.139 |
| 182 | T | CCRK | VDQGDLMTPQFTPYY | 3.143 | 3.11 |
| 182 | T | GSK3B | VDQGDLMTPQFTPYY | 9.568 | 7.881 |
| 182 | T | p38 | VDQGDLMTPQFTPYY | 7.91 | 7.426 |
| 182 | T | MEKK2 | VDQGDLMTPQFTPYY | 22.931 | 21.636 |
| 182 | T | KHS | VDQGDLMTPQFTPYY | 0.83 | 0.771 |
| 182 | T | NinaC | VDQGDLMTPQFTPYY | 4.954 | 4.159 |
| 182 | T | YSK | VDQGDLMTPQFTPYY | 151.926 | 133.749 |
| 182 | T | GTF2F1 | VDQGDLMTPQFTPYY | 3.195 | 2.841 |
| 182 | T | BUB1 | VDQGDLMTPQFTPYY | 2.01E-5 | 0E0 |
| 182 | T | CAMKK-Meta | VDQGDLMTPQFTPYY | 0.001 | 0.001 |
| 182 | T | BIKE | VDQGDLMTPQFTPYY | 3.625 | 3.624 |
| 182 | T | GRK1 | VDQGDLMTPQFTPYY | 15.584 | 10.931 |
| 182 | T | CDK20 | VDQGDLMTPQFTPYY | 3.143 | 3.11 |
| 182 | T | CDK2 | VDQGDLMTPQFTPYY | 14.865 | 14.749 |
| 182 | T | CDK6 | VDQGDLMTPQFTPYY | 84.439 | 84.402 |
| 182 | T | PHO85 | VDQGDLMTPQFTPYY | 3.336 | 3.101 |
| 182 | T | HIPK1 | VDQGDLMTPQFTPYY | 3.835 | 3.685 |
| 182 | T | ERK4 (MAPK4) | VDQGDLMTPQFTPYY | 9.957 | 2.921 |
| 182 | T | ERK3 (MAPK6) | VDQGDLMTPQFTPYY | 7.549 | 2.9 |
| 182 | T | p38 beta (MAPK11) | VDQGDLMTPQFTPYY | 48.087 | 42.826 |
| 182 | T | p38 gamma (MAPK12) | VDQGDLMTPQFTPYY | 129.007 | 127.287 |
| 182 | T | p38 delta (MAPK13) | VDQGDLMTPQFTPYY | 47.16 | 45.539 |
| 182 | T | MYO3A | VDQGDLMTPQFTPYY | 4.954 | 4.159 |
| 182 | T | TAOK1 | VDQGDLMTPQFTPYY | 4.957 | 3.53 |
| 182 | T | STK24 | VDQGDLMTPQFTPYY | 4.466 | 3.424 |
| 182 | T | MAP3K10 | VDQGDLMTPQFTPYY | 4.337 | 3.409 |
| 182 | T | TRPM6 | VDQGDLMTPQFTPYY | 2.025 | 1.899 |
| 182 | T | AAK1 | VDQGDLMTPQFTPYY | 3.625 | 3.624 |
| 186 | T | TKL | DLMTPQFTPYYVAPQ | 6.379 | 5.299 |
| 186 | T | MAPK | DLMTPQFTPYYVAPQ | 13.824 | 12.837 |
| 186 | T | LISK | DLMTPQFTPYYVAPQ | 4.989 | 3.139 |
| 186 | T | LKB | DLMTPQFTPYYVAPQ | 32.229 | 24.571 |
| 186 | T | CDK2 | DLMTPQFTPYYVAPQ | 10.78 | 10.318 |
| 186 | T | CRK7 | DLMTPQFTPYYVAPQ | 4.647 | 4.506 |
| 186 | T | ERK1 | DLMTPQFTPYYVAPQ | 11.859 | 11.368 |
| 186 | T | p38 | DLMTPQFTPYYVAPQ | 7.541 | 7.426 |
| 186 | T | TESK | DLMTPQFTPYYVAPQ | 5.084 | 3.286 |
| 186 | T | ILK | DLMTPQFTPYYVAPQ | 23.234 | 21.936 |
| 186 | T | BUB1 | DLMTPQFTPYYVAPQ | 2.52E-5 | 0E0 |
| 186 | T | STK11 | DLMTPQFTPYYVAPQ | 32.327 | 24.553 |
| 186 | T | PHO85 | DLMTPQFTPYYVAPQ | 3.411 | 3.101 |
| 186 | T | CDK12 | DLMTPQFTPYYVAPQ | 4.647 | 4.506 |
| 186 | T | HIPK1 | DLMTPQFTPYYVAPQ | 5.754 | 3.685 |
| 186 | T | ERK2 (MAPK1) | DLMTPQFTPYYVAPQ | 15.903 | 14.663 |
| 186 | T | JNK3 (MAPK10) | DLMTPQFTPYYVAPQ | 0.023 | 0.023 |
| 186 | T | p38 delta (MAPK13) | DLMTPQFTPYYVAPQ | 50.409 | 45.539 |
| 186 | T | p38 alpha (MAPK14) | DLMTPQFTPYYVAPQ | 12.399 | 11.017 |
| 186 | T | MAP2K6 | DLMTPQFTPYYVAPQ | 3.535 | 3.308 |
| 186 | T | TESK1 | DLMTPQFTPYYVAPQ | 5.074 | 3.271 |
| 186 | T | ILK | DLMTPQFTPYYVAPQ | 23.234 | 21.936 |
| 206 | S | NEK | RRHQKEKSGIIPTSP | 122.577 | 118.615 |
| 206 | S | NDR | RRHQKEKSGIIPTSP | 0.606 | 0.407 |
| 206 | S | CAMK2G | RRHQKEKSGIIPTSP | 4.022 | 3.87 |
| 206 | S | PIM1 | RRHQKEKSGIIPTSP | 0.001 | 0.001 |
| 206 | S | BUB1 | RRHQKEKSGIIPTSP | 3.8E-5 | 0E0 |
| 206 | S | WNK3 | RRHQKEKSGIIPTSP | 3.305 | 2.995 |
| 206 | S | GRK7 | RRHQKEKSGIIPTSP | 1.741 | 1.646 |
| 206 | S | TK38 | RRHQKEKSGIIPTSP | 1.618 | 0.393 |
| 211 | T | PDHK | EKSGIIPTSPTPYTY | 24.755 | 24.593 |
| 211 | T | MYLK | EKSGIIPTSPTPYTY | 1.457 | 1.449 |
| 211 | T | BUB1 | EKSGIIPTSPTPYTY | 2.11E-5 | 0E0 |
| 212 | S | CDK | KSGIIPTSPTPYTYN | 12.36 | 10.587 |
| 212 | S | Alpha | KSGIIPTSPTPYTYN | 12.343 | 12.011 |
| 212 | S | IKK | KSGIIPTSPTPYTYN | 9.699 | 8.821 |
| 212 | S | CCRK | KSGIIPTSPTPYTYN | 3.69 | 3.11 |
| 212 | S | CDC2 | KSGIIPTSPTPYTYN | 10.385 | 9.122 |
| 212 | S | CDK2 | KSGIIPTSPTPYTYN | 10.69 | 10.318 |
| 212 | S | CDK4 | KSGIIPTSPTPYTYN | 9.391 | 8.423 |
| 212 | S | CDK5 | KSGIIPTSPTPYTYN | 29.615 | 27.329 |
| 212 | S | CDK7 | KSGIIPTSPTPYTYN | 22.523 | 20.158 |
| 212 | S | CDK8 | KSGIIPTSPTPYTYN | 9.318 | 9.04 |
| 212 | S | HIPK | KSGIIPTSPTPYTYN | 7.239 | 7.023 |
| 212 | S | GSK3A | KSGIIPTSPTPYTYN | 97.424 | 82.191 |
| 212 | S | ERK | KSGIIPTSPTPYTYN | 28.267 | 25.065 |
| 212 | S | ERK1 | KSGIIPTSPTPYTYN | 12.232 | 11.368 |
| 212 | S | JNK | KSGIIPTSPTPYTYN | 11.709 | 10.69 |
| 212 | S | nmo | KSGIIPTSPTPYTYN | 0.308 | 0.259 |
| 212 | S | BUB1 | KSGIIPTSPTPYTYN | 5.19E-5 | 0E0 |
| 212 | S | TBK1 | KSGIIPTSPTPYTYN | 184.464 | 166.364 |
| 212 | S | CDK20 | KSGIIPTSPTPYTYN | 3.69 | 3.11 |
| 212 | S | CDK2 | KSGIIPTSPTPYTYN | 15.156 | 14.749 |
| 212 | S | CDK3 | KSGIIPTSPTPYTYN | 9.136 | 9.024 |
| 212 | S | CDK4 | KSGIIPTSPTPYTYN | 8.897 | 7.735 |
| 212 | S | CDK6 | KSGIIPTSPTPYTYN | 85.801 | 84.402 |
| 212 | S | CDK5 | KSGIIPTSPTPYTYN | 18.99 | 17.626 |
| 212 | S | CDK7 | KSGIIPTSPTPYTYN | 21.903 | 20.865 |
| 212 | S | CDK8 | KSGIIPTSPTPYTYN | 49.327 | 35.984 |
| 212 | S | CDK9 | KSGIIPTSPTPYTYN | 5.692 | 5.019 |
| 212 | S | HIPK2 | KSGIIPTSPTPYTYN | 113.803 | 109.385 |
| 212 | S | HIPK3 | KSGIIPTSPTPYTYN | 4.77 | 4.386 |
| 212 | S | ERK2 (MAPK1) | KSGIIPTSPTPYTYN | 14.838 | 14.663 |
| 212 | S | ERK1 (MAPK3) | KSGIIPTSPTPYTYN | 18.368 | 16.01 |
| 212 | S | JNK3 (MAPK10) | KSGIIPTSPTPYTYN | 0.024 | 0.023 |
| 212 | S | JNK2 (MAPK9) | KSGIIPTSPTPYTYN | 161.944 | 155.771 |
| 212 | S | NLK | KSGIIPTSPTPYTYN | 0.308 | 0.259 |
| 212 | S | p38 gamma (MAPK12) | KSGIIPTSPTPYTYN | 131.772 | 127.287 |
| 212 | S | p38 alpha (MAPK14) | KSGIIPTSPTPYTYN | 11.797 | 11.017 |
| 212 | S | MAP2K3 | KSGIIPTSPTPYTYN | 2.71E-5 | 0E0 |
| 214 | T | CDK | GIIPTSPTPYTYNKS | 11.158 | 10.587 |
| 214 | T | DYRK | GIIPTSPTPYTYNKS | 19.853 | 17.356 |
| 214 | T | BUB | GIIPTSPTPYTYNKS | 0.342 | 0.314 |
| 214 | T | KIS | GIIPTSPTPYTYNKS | 2.951 | 2.45 |
| 214 | T | CCRK | GIIPTSPTPYTYNKS | 3.424 | 3.11 |
| 214 | T | CDK2 | GIIPTSPTPYTYNKS | 11.498 | 10.318 |
| 214 | T | CDK5 | GIIPTSPTPYTYNKS | 28.609 | 27.329 |
| 214 | T | CDK9 | GIIPTSPTPYTYNKS | 40.501 | 36.669 |
| 214 | T | CRK7 | GIIPTSPTPYTYNKS | 4.95 | 4.506 |
| 214 | T | DYRK1 | GIIPTSPTPYTYNKS | 27.581 | 21.744 |
| 214 | T | DYRK2 | GIIPTSPTPYTYNKS | 11.963 | 10.537 |
| 214 | T | FRAP | GIIPTSPTPYTYNKS | 11.86 | 9.635 |
| 214 | T | BUB1 | GIIPTSPTPYTYNKS | 7.39E-5 | 0E0 |
| 214 | T | UHMK1 | GIIPTSPTPYTYNKS | 2.951 | 2.45 |
| 214 | T | CDK20 | GIIPTSPTPYTYNKS | 3.424 | 3.11 |
| 214 | T | CDC28 | GIIPTSPTPYTYNKS | 16.669 | 15.641 |
| 214 | T | CDK6 | GIIPTSPTPYTYNKS | 84.951 | 84.402 |
| 214 | T | CDK5 | GIIPTSPTPYTYNKS | 18.493 | 17.626 |
| 214 | T | PHO85 | GIIPTSPTPYTYNKS | 3.419 | 3.101 |
| 214 | T | CDK9 | GIIPTSPTPYTYNKS | 5.685 | 5.019 |
| 214 | T | CDK12 | GIIPTSPTPYTYNKS | 4.95 | 4.506 |
| 214 | T | DYRK1B | GIIPTSPTPYTYNKS | 0.104 | 0.087 |
| 214 | T | FUS3 | GIIPTSPTPYTYNKS | 3.955 | 2.834 |
| 214 | T | JNK3 (MAPK10) | GIIPTSPTPYTYNKS | 0.023 | 0.023 |
| 214 | T | JNK2 (MAPK9) | GIIPTSPTPYTYNKS | 156.012 | 155.771 |
| 214 | T | p38 gamma (MAPK12) | GIIPTSPTPYTYNKS | 127.828 | 127.287 |
| 214 | T | p38 delta (MAPK13) | GIIPTSPTPYTYNKS | 45.665 | 45.539 |
| 214 | T | MTOR | GIIPTSPTPYTYNKS | 12.123 | 10.962 |
| 217 | T | FRAY | PTSPTPYTYNKSCDL | 90.696 | 86.942 |
| 217 | T | BUB1 | PTSPTPYTYNKSCDL | 1.11E-5 | 0E0 |
| 217 | T | Tec | PTSPTPYTYNKSCDL | 3.189 | 3.05 |
| 217 | T | ACVRL1 | PTSPTPYTYNKSCDL | 1.204 | 1.091 |
| 217 | T | PDK2 | PTSPTPYTYNKSCDL | 2.13 | 1.828 |
| 217 | T | BTK | PTSPTPYTYNKSCDL | 3.189 | 3.05 |
| 221 | S | BUB | TPYTYNKSCDLWSLG | 0.331 | 0.314 |
| 221 | S | BUB1 | TPYTYNKSCDLWSLG | 6.01E-5 | 0E0 |
| 221 | S | TGFBR1 | TPYTYNKSCDLWSLG | 22.172 | 17.6 |
| 226 | S | BUB1 | NKSCDLWSLGVIIYV | 2.78E-5 | 0E0 |
| 243 | S | BUB1 | CGYPPFYSKHHSRTI | 1.31E-5 | 0E0 |
| 243 | S | GRK1 | CGYPPFYSKHHSRTI | 18.391 | 10.931 |
| 243 | S | CAMKK1 | CGYPPFYSKHHSRTI | 1.722 | 1.714 |
| 243 | S | ULK3 | CGYPPFYSKHHSRTI | 3.094 | 2.286 |
| 247 | S | SLK | PFYSKHHSRTIPKDM | 4.478 | 4.036 |
| 247 | S | BUB1 | PFYSKHHSRTIPKDM | 2.27E-5 | 0E0 |
| 247 | S | GRK1 | PFYSKHHSRTIPKDM | 18.202 | 10.931 |
| 247 | S | PRKCG | PFYSKHHSRTIPKDM | 136.61 | 116.631 |
| 247 | S | PAK4 | PFYSKHHSRTIPKDM | 22.22 | 19.402 |
| 247 | S | ULK1 | PFYSKHHSRTIPKDM | 38.955 | 38.389 |
| 249 | T | STE-Unique | YSKHHSRTIPKDMRK | 11.874 | 10.171 |
| 249 | T | PIM2 | YSKHHSRTIPKDMRK | 6.652 | 6.373 |
| 249 | T | MAP3K8 | YSKHHSRTIPKDMRK | 22.156 | 20.344 |
| 249 | T | FRAY | YSKHHSRTIPKDMRK | 138.775 | 86.942 |
| 249 | T | KSR | YSKHHSRTIPKDMRK | 2.703 | 2.391 |
| 249 | T | BUB1 | YSKHHSRTIPKDMRK | 4.57E-5 | 0E0 |
| 249 | T | STK39 | YSKHHSRTIPKDMRK | 115.823 | 86.489 |
| 260 | T | AKT2 | DMRKKIMTGSFEFPE | 24.953 | 24.834 |
| 260 | T | CRIK | DMRKKIMTGSFEFPE | 1.033 | 0.919 |
| 260 | T | PIM1 | DMRKKIMTGSFEFPE | 0.002 | 0.001 |
| 260 | T | PIM2 | DMRKKIMTGSFEFPE | 6.943 | 6.373 |
| 260 | T | ILK | DMRKKIMTGSFEFPE | 23.244 | 21.936 |
| 260 | T | BUB1 | DMRKKIMTGSFEFPE | 3.52E-5 | 0E0 |
| 260 | T | InsR | DMRKKIMTGSFEFPE | 5.215 | 4.34 |
| 260 | T | CIT | DMRKKIMTGSFEFPE | 1.033 | 0.919 |
| 260 | T | DAPK2 | DMRKKIMTGSFEFPE | 4.871 | 3.067 |
| 260 | T | STK10 | DMRKKIMTGSFEFPE | 3.645 | 3.565 |
| 260 | T | MAP2K2 | DMRKKIMTGSFEFPE | 4.951 | 3.157 |
| 260 | T | ILK | DMRKKIMTGSFEFPE | 23.244 | 21.936 |
| 260 | T | BRAF | DMRKKIMTGSFEFPE | 3.272 | 2.945 |
| 262 | S | PIM | RKKIMTGSFEFPEEE | 24.6 | 15.765 |
| 262 | S | BUB | RKKIMTGSFEFPEEE | 0.348 | 0.314 |
| 262 | S | CAMK2D | RKKIMTGSFEFPEEE | 55.852 | 50.397 |
| 262 | S | CK1-D | RKKIMTGSFEFPEEE | 11.716 | 8.479 |
| 262 | S | PDHK | RKKIMTGSFEFPEEE | 6.233 | 5.468 |
| 262 | S | BUB1 | RKKIMTGSFEFPEEE | 5.41E-5 | 0E0 |
| 262 | S | PLK2 | RKKIMTGSFEFPEEE | 23.743 | 19.736 |
| 262 | S | Myt1 | RKKIMTGSFEFPEEE | 3.408 | 2.486 |
| 262 | S | DAPK3 | RKKIMTGSFEFPEEE | 9.412 | 9.069 |
| 262 | S | ULK2 | RKKIMTGSFEFPEEE | 4.098 | 4.057 |
| 262 | S | PKMYT1 | RKKIMTGSFEFPEEE | 3.408 | 2.486 |
| 271 | S | CDKL | EFPEEEWSQISEMAK | 5.013 | 4.358 |
| 271 | S | PIKK | EFPEEEWSQISEMAK | 13.605 | 12.124 |
| 271 | S | CK1-D | EFPEEEWSQISEMAK | 8.704 | 8.479 |
| 271 | S | CDKL5 | EFPEEEWSQISEMAK | 5.013 | 4.358 |
| 271 | S | ATM | EFPEEEWSQISEMAK | 13.893 | 13.727 |
| 271 | S | SMG1 | EFPEEEWSQISEMAK | 3.627 | 3.251 |
| 271 | S | BUB1 | EFPEEEWSQISEMAK | 8.19E-5 | 0E0 |
| 271 | S | CSNK1D | EFPEEEWSQISEMAK | 18.197 | 15.529 |
| 271 | S | STK4 | EFPEEEWSQISEMAK | 5.027 | 4.959 |
| 271 | S | ATM | EFPEEEWSQISEMAK | 13.893 | 13.727 |
| 271 | S | SMG1 | EFPEEEWSQISEMAK | 3.627 | 3.251 |
| 274 | S | IRAK | EEEWSQISEMAKDVV | 3.397 | 3.391 |
| 274 | S | MEK1 | EEEWSQISEMAKDVV | 156.777 | 150.481 |
| 274 | S | BUB1 | EEEWSQISEMAKDVV | 3.73E-5 | 0E0 |
| 274 | S | MPSK | EEEWSQISEMAKDVV | 2.423 | 2.409 |
| 274 | S | GRK3 | EEEWSQISEMAKDVV | 13.496 | 13.381 |
| 274 | S | DAPK1 | EEEWSQISEMAKDVV | 14.451 | 12.301 |
| 274 | S | MAPKAPK5 | EEEWSQISEMAKDVV | 33.901 | 31.873 |
| 274 | S | CSNK1A1 | EEEWSQISEMAKDVV | 25.044 | 24.196 |
| 274 | S | STK16 | EEEWSQISEMAKDVV | 2.423 | 2.409 |
| 274 | S | SRC | EEEWSQISEMAKDVV | 3.822 | 3.708 |
| 294 | T | MYLK | VKPEERLTIEGVLDH | 1.762 | 1.449 |
| 294 | T | IPL1 | VKPEERLTIEGVLDH | 4.639 | 3.33 |
| 294 | T | BUB1 | VKPEERLTIEGVLDH | 7.58E-6 | 0E0 |
| 294 | T | STK11 | VKPEERLTIEGVLDH | 24.628 | 24.553 |
| 294 | T | DAPK2 | VKPEERLTIEGVLDH | 4.099 | 3.067 |
| 294 | T | CSNK1D | VKPEERLTIEGVLDH | 18.047 | 15.529 |
| 294 | T | CLA4 | VKPEERLTIEGVLDH | 3.68 | 3.489 |
| 306 | S | RIO | LDHPWLNSTEALDNV | 3.061 | 2.731 |
| 306 | S | BUB1 | LDHPWLNSTEALDNV | 2.55E-5 | 0E0 |
| 306 | S | STK38 | LDHPWLNSTEALDNV | 1.626 | 0.393 |
| 306 | S | SIK2 | LDHPWLNSTEALDNV | 1.668 | 1.586 |
| 307 | T | MLK | DHPWLNSTEALDNVL | 3.544 | 3.15 |
| 307 | T | BUB1 | DHPWLNSTEALDNVL | 2.6E-5 | 0E0 |
| 307 | T | WNK4 | DHPWLNSTEALDNVL | 3.384 | 3.101 |
| 307 | T | MAP2K3 | DHPWLNSTEALDNVL | 1.7E-5 | 0E0 |
| 307 | T | MAP2K6 | DHPWLNSTEALDNVL | 4.085 | 3.308 |
| 316 | S | TSSK4 | ALDNVLPSAQLMMDK | 3.507 | 2.917 |
| 316 | S | BUB1 | ALDNVLPSAQLMMDK | 3.97E-5 | 0E0 |
| 316 | S | BMPR1B | ALDNVLPSAQLMMDK | -0.964 | -1.013 |
| 348 | S | STE11 | RIQDLKVSLKPLHSV | 5.204 | 5.196 |
| 348 | S | NEK | RIQDLKVSLKPLHSV | 132.849 | 118.615 |
| 348 | S | ULK | RIQDLKVSLKPLHSV | 38.936 | 36.383 |
| 348 | S | PKCd | RIQDLKVSLKPLHSV | -9.168 | -9.497 |
| 348 | S | MELK | RIQDLKVSLKPLHSV | 201.925 | 195.594 |
| 348 | S | IRAK1 | RIQDLKVSLKPLHSV | 115.359 | 89.042 |
| 348 | S | ULK | RIQDLKVSLKPLHSV | 34.187 | 31.912 |
| 348 | S | PRKCD | RIQDLKVSLKPLHSV | 12.604 | 11.142 |
| 348 | S | MELK | RIQDLKVSLKPLHSV | 201.925 | 195.594 |
| 348 | S | SIK2 | RIQDLKVSLKPLHSV | 1.772 | 1.586 |
| 348 | S | MAP2K3 | RIQDLKVSLKPLHSV | 5.63E-5 | 0E0 |
| 348 | S | ULK1 | RIQDLKVSLKPLHSV | 41.919 | 38.389 |
| 348 | S | ULK2 | RIQDLKVSLKPLHSV | 4.306 | 4.057 |
| 354 | S | IKK | VSLKPLHSVNNPILR | 9.225 | 8.821 |
| 354 | S | PEK | VSLKPLHSVNNPILR | 1.823 | 1.697 |
| 354 | S | PIM3 | VSLKPLHSVNNPILR | 5.457 | 5.258 |
| 354 | S | BUB1 | VSLKPLHSVNNPILR | 1.7E-5 | 0E0 |
| 354 | S | NEK6 | VSLKPLHSVNNPILR | 39.702 | 39.149 |
| 354 | S | BRSK2 | VSLKPLHSVNNPILR | 4.64 | 4.325 |
| 354 | S | DAPK2 | VSLKPLHSVNNPILR | 3.2 | 3.067 |
| 368 | T | RAD53 | RKRKLLGTKPKDGIY | 6.613 | 6.3 |
| 368 | T | PIM1 | RKRKLLGTKPKDGIY | 0.002 | 0.001 |
| 368 | T | VRK1 | RKRKLLGTKPKDGIY | 0.412 | 0.394 |
| 383 | T | NKF2 | IHDHENGTEDSNVAL | 18.848 | 15.808 |
| 383 | T | TTK | IHDHENGTEDSNVAL | 25.511 | 21.21 |
| 383 | T | BARK | IHDHENGTEDSNVAL | 181.432 | 165.626 |
| 383 | T | FRAY | IHDHENGTEDSNVAL | 122.293 | 86.942 |
| 383 | T | MEK4 | IHDHENGTEDSNVAL | 0.009 | 0.009 |
| 383 | T | MLK | IHDHENGTEDSNVAL | 3.413 | 3.15 |
| 383 | T | BUB1 | IHDHENGTEDSNVAL | 1.3E-5 | 0E0 |
| 383 | T | PINK1 | IHDHENGTEDSNVAL | 18.848 | 15.808 |
| 383 | T | TTK | IHDHENGTEDSNVAL | 30.306 | 22.818 |
| 383 | T | GRK2 | IHDHENGTEDSNVAL | 160.353 | 147.674 |
| 383 | T | STK39 | IHDHENGTEDSNVAL | 114.454 | 86.489 |
| 383 | T | MAP2K4 | IHDHENGTEDSNVAL | 0.009 | 0.009 |
| 386 | S | GRK | HENGTEDSNVALEKL | 4.198 | 2.938 |
| 386 | S | CLK2 | HENGTEDSNVALEKL | 0.007 | 0.006 |
| 386 | S | RIPK1 | HENGTEDSNVALEKL | 4.192 | 3.773 |
| 386 | S | BUB1 | HENGTEDSNVALEKL | 3.01E-5 | 0E0 |
| 386 | S | TLK2 | HENGTEDSNVALEKL | 2.536 | 2.396 |
| 386 | S | Tec | HENGTEDSNVALEKL | 3.147 | 3.05 |
| 386 | S | CDC42BPA | HENGTEDSNVALEKL | 3.523 | 2.821 |
| 386 | S | PRKCG | HENGTEDSNVALEKL | 118.843 | 116.631 |
| 386 | S | SIK2 | HENGTEDSNVALEKL | 1.696 | 1.586 |
| 386 | S | DAPK1 | HENGTEDSNVALEKL | 13.02 | 12.301 |
| 386 | S | CSNK1G2 | HENGTEDSNVALEKL | 3.938 | 3.273 |
| 386 | S | BTK | HENGTEDSNVALEKL | 3.147 | 3.05 |
| 438 | S | SGK | LLRDALQSFSWNGRG | 15.385 | 14.019 |
| 438 | S | PLK | LLRDALQSFSWNGRG | 14.808 | 14.476 |
| 438 | S | SGK1 | LLRDALQSFSWNGRG | 15.939 | 13.277 |
| 438 | S | PIM1 | LLRDALQSFSWNGRG | 0.001 | 0.001 |
| 438 | S | BUB1 | LLRDALQSFSWNGRG | 2.91E-5 | 0E0 |
| 438 | S | WNK2 | LLRDALQSFSWNGRG | 3.859 | 3.369 |
| 438 | S | PRKCI | LLRDALQSFSWNGRG | 12.625 | 9.121 |
| 438 | S | DAPK2 | LLRDALQSFSWNGRG | 4.074 | 3.067 |
| 438 | S | TRPM6 | LLRDALQSFSWNGRG | 3.166 | 1.899 |
| 440 | S | MEKK2 | RDALQSFSWNGRGFT | 24.15 | 21.636 |
| 440 | S | BUB1 | RDALQSFSWNGRGFT | 5.82E-5 | 0E0 |
| 440 | S | PRKAA1 | RDALQSFSWNGRGFT | 69.828 | 63.488 |
| 440 | S | CDK19 | RDALQSFSWNGRGFT | 3.467 | 3.342 |
| 440 | S | MAP3K3 | RDALQSFSWNGRGFT | 15.963 | 15.283 |
| 440 | S | MAP2K3 | RDALQSFSWNGRGFT | 6.08E-6 | 0E0 |
| 447 | T | AGC | SWNGRGFTDKVDRLK | 3.778 | 1.925 |
| 447 | T | PKC | SWNGRGFTDKVDRLK | -2.817 | -3.397 |
| 447 | T | PKD | SWNGRGFTDKVDRLK | 21.458 | 21.331 |
| 447 | T | RCK | SWNGRGFTDKVDRLK | 4.489 | 4.171 |
| 447 | T | BCR | SWNGRGFTDKVDRLK | 3.811 | 3.684 |
| 447 | T | PEK | SWNGRGFTDKVDRLK | 1.757 | 1.697 |
| 447 | T | AKT1 | SWNGRGFTDKVDRLK | 12.082 | 11.65 |
| 447 | T | ROCK | SWNGRGFTDKVDRLK | 14.172 | 13.925 |
| 447 | T | PKCa | SWNGRGFTDKVDRLK | 10.106 | 9.03 |
| 447 | T | PKCd | SWNGRGFTDKVDRLK | -6.28 | -9.497 |
| 447 | T | PKCh | SWNGRGFTDKVDRLK | 14.046 | 10.374 |
| 447 | T | PKCi | SWNGRGFTDKVDRLK | 7.449 | 3.394 |
| 447 | T | PIM1 | SWNGRGFTDKVDRLK | 0.001 | 0.001 |
| 447 | T | CHEK2 | SWNGRGFTDKVDRLK | 13.713 | 12.371 |
| 447 | T | VRK2 | SWNGRGFTDKVDRLK | 57.868 | 48.397 |
| 447 | T | MAK | SWNGRGFTDKVDRLK | 4.489 | 4.171 |
| 447 | T | STKR2 | SWNGRGFTDKVDRLK | 1.503 | 1.301 |
| 447 | T | BCR | SWNGRGFTDKVDRLK | 3.811 | 3.684 |
| 447 | T | BUB1 | SWNGRGFTDKVDRLK | 6.39E-5 | 0E0 |
| 447 | T | GRK3 | SWNGRGFTDKVDRLK | 18.372 | 13.381 |
| 447 | T | GRK1 | SWNGRGFTDKVDRLK | 13.197 | 10.931 |
| 447 | T | LATS2 | SWNGRGFTDKVDRLK | 0.051 | 0.05 |
| 447 | T | PRKCD | SWNGRGFTDKVDRLK | 13.465 | 11.142 |
| 447 | T | PRKCE | SWNGRGFTDKVDRLK | 24.583 | 18.443 |
| 447 | T | PRKCI | SWNGRGFTDKVDRLK | 9.528 | 9.121 |
| 447 | T | DAPK1 | SWNGRGFTDKVDRLK | 12.461 | 12.301 |
| 447 | T | DYRK3 | SWNGRGFTDKVDRLK | 3.013 | 2.653 |
| 447 | T | ICK | SWNGRGFTDKVDRLK | 4.489 | 4.171 |
| 447 | T | PAK6 | SWNGRGFTDKVDRLK | 2.509 | 2.163 |
| 447 | T | ACVRL1 | SWNGRGFTDKVDRLK | 1.196 | 1.091 |
| 447 | T | BMPR1B | SWNGRGFTDKVDRLK | 0.66 | -1.013 |
| 447 | T | TGFBR2 | SWNGRGFTDKVDRLK | 1.503 | 1.301 |
| 467 | T | CDKL | KQVIEEQTLPHEPQ* | 5.649 | 4.358 |
| 467 | T | LRRK | KQVIEEQTLPHEPQ* | 16.576 | 13.084 |
| 467 | T | STKR | KQVIEEQTLPHEPQ* | 23.24 | 21.849 |
| 467 | T | MOS | KQVIEEQTLPHEPQ* | 3.323 | 2.885 |
| 467 | T | PEK | KQVIEEQTLPHEPQ* | 4.955 | 1.697 |
| 467 | T | PLK | KQVIEEQTLPHEPQ* | 16.936 | 14.476 |
| 467 | T | PRKD2 | KQVIEEQTLPHEPQ* | 2.71 | 2.35 |
| 467 | T | CDKL5 | KQVIEEQTLPHEPQ* | 5.649 | 4.358 |
| 467 | T | SLK | KQVIEEQTLPHEPQ* | 4.317 | 4.036 |
| 467 | T | TAO | KQVIEEQTLPHEPQ* | 4.514 | 3.655 |
| 467 | T | LRRK2 | KQVIEEQTLPHEPQ* | 21.683 | 17.83 |
| 467 | T | BUB1 | KQVIEEQTLPHEPQ* | 5.07E-5 | 0E0 |
| 467 | T | MOS | KQVIEEQTLPHEPQ* | 3.323 | 2.885 |
| 467 | T | HRI | KQVIEEQTLPHEPQ* | 4.152 | 3.713 |
| 467 | T | PLK1 | KQVIEEQTLPHEPQ* | 13.612 | 13.525 |
| 467 | T | TLK2 | KQVIEEQTLPHEPQ* | 2.51 | 2.396 |
| 467 | T | TTK | KQVIEEQTLPHEPQ* | 25.217 | 22.818 |
| 467 | T | GRK3 | KQVIEEQTLPHEPQ* | 16.467 | 13.381 |
| 467 | T | GRK1 | KQVIEEQTLPHEPQ* | 14.152 | 10.931 |
| 467 | T | GRK5 | KQVIEEQTLPHEPQ* | 2.738 | 2.677 |
| 467 | T | GRK7 | KQVIEEQTLPHEPQ* | 2.819 | 1.646 |
| 467 | T | PRKCH | KQVIEEQTLPHEPQ* | 28.913 | 28.536 |
| 467 | T | STK17B | KQVIEEQTLPHEPQ* | 3.268 | 3.169 |
| 467 | T | CSNK1D | KQVIEEQTLPHEPQ* | 15.621 | 15.529 |
| 467 | T | DYRK1B | KQVIEEQTLPHEPQ* | 0.166 | 0.087 |
| 467 | T | STK3 | KQVIEEQTLPHEPQ* | 92.135 | 73.61 |
| 467 | T | STK4 | KQVIEEQTLPHEPQ* | 4.968 | 4.959 |
| 467 | T | STK10 | KQVIEEQTLPHEPQ* | 4.2 | 3.565 |
| 467 | T | MAP2K3 | KQVIEEQTLPHEPQ* | 1.64E-4 | 0E0 |
| 467 | T | ACVRL1 | KQVIEEQTLPHEPQ* | 5.678 | 1.091 |
| 467 | T | BMPR1B | KQVIEEQTLPHEPQ* | 15.079 | -1.013 |
| 467 | T | PDK1 | KQVIEEQTLPHEPQ* | 0.209 | 0.182 |
| 467 | T | EIF2AK1 | KQVIEEQTLPHEPQ* | 4.152 | 3.713 |

Group-based Prediction System (GPS; http://gps.biocuckoo.cn) output predicting serine and threonine residues phosphorylated in mouse MK5.1 (NCBI NP_034895.1) and their cognate protein kinases. The peptide sequence shows the predicted phosphorylated amino acid with 7 flanking amino acids. The score refers to the value calculated by GPS5 evaluate the theoretical potential a protein kinase phosphorylating the indicated residue ( higher value = higher potential. The cut-off value for a predicted phosphorylation site using e high threshold indicates a value resulting in a 2% false positive rate.
