## Supplemental Table 2 for "Expression, subcellular localization, and phosphorylation of MK5 in adult cardiac ventricular fibroblasts"

**Supplemental Table 2. Predicted tyrosine phosphorylation sites and their cognate protein kinases in mouse MK5.1.**

| position | residue | kinase | Peptide sequence | score | cutoff |
| --- | --- | --- | --- | --- | --- |
| 20 | Y | EGFR | ETSILEEYSINWTQK | 9.527 | 8.652 |
| 20 | Y | Tec | ETSILEEYSINWTQK | 21.654 | 14.555 |
| 20 | Y | AGC | ETSILEEYSINWTQK | 2.323 | 2.065 |
| 20 | Y | CAMK | ETSILEEYSINWTQK | 1.861 | 1.637 |
| 20 | Y | ABL1 | ETSILEEYSINWTQK | 9.248 | 9.035 |
| 20 | Y | ERBB4 | ETSILEEYSINWTQK | 5.46E-7 | 0E0 |
| 20 | Y | FGFR3 | ETSILEEYSINWTQK | 9.91E-8 | 0E0 |
| 20 | Y | PTK2 | ETSILEEYSINWTQK | 5.53E-5 | 0E0 |
| 20 | Y | INSR | ETSILEEYSINWTQK | 14.568 | 13.552 |
| 20 | Y | CSF1R | ETSILEEYSINWTQK | 1.46E-4 | 0E0 |
| 20 | Y | SRM | ETSILEEYSINWTQK | 2.542 | 1.726 |
| 20 | Y | NTRK1 | ETSILEEYSINWTQK | 3.95E-7 | 0E0 |
| 20 | Y | PEK | ETSILEEYSINWTQK | 2.107 | 1.815 |
| 20 | Y | WEE1B | ETSILEEYSINWTQK | 6.23 | 2.072 |
| 188 | Y | Ack | MTPQFTPYYVAPQVL | 2.49E-6 | 0E0 |
| 188 | Y | TTrk | MTPQFTPYYVAPQVL | 45.32 | 41.671 |
| 188 | Y | Dual/STE | MTPQFTPYYVAPQVL | 0.436 | 0.402 |
| 188 | Y | TNK2 | MTPQFTPYYVAPQVL | 2.49E-6 | 0E0 |
| 188 | Y | ERBB4 | MTPQFTPYYVAPQVL | 5.93E-7 | 0E0 |
| 188 | Y | FGFR3 | MTPQFTPYYVAPQVL | 3.31E-7 | 0E0 |
| 188 | Y | PTK2 | MTPQFTPYYVAPQVL | 1.8E-5 | 0E0 |
| 188 | Y | NTRK1 | MTPQFTPYYVAPQVL | 3.08E-7 | 0E0 |
| 188 | Y | NTRK2 | MTPQFTPYYVAPQVL | 3.58E-6 | 0E0 |
| 189 | Y | Abl | TPQFTPYYVAPQVLE | 12.492 | 10.599 |
| 189 | Y | Ack | TPQFTPYYVAPQVLE | 2.4E-7 | 0E0 |
| 189 | Y | EGFR | TPQFTPYYVAPQVLE | 9.856 | 8.652 |
| 189 | Y | Eph | TPQFTPYYVAPQVLE | 1.697 | 1.41 |
| 189 | Y | ABL1 | TPQFTPYYVAPQVLE | 10.649 | 9.035 |
| 189 | Y | TNK2 | TPQFTPYYVAPQVLE | 2.4E-7 | 0E0 |
| 189 | Y | MERTK | TPQFTPYYVAPQVLE | 2.609 | 2.6 |
| 189 | Y | ERBB4 | TPQFTPYYVAPQVLE | 1.87E-6 | 0E0 |
| 189 | Y | EPHB2 | TPQFTPYYVAPQVLE | 0.64 | 0.554 |
| 189 | Y | FGFR3 | TPQFTPYYVAPQVLE | 1.09E-6 | 0E0 |
| 189 | Y | PTK2 | TPQFTPYYVAPQVLE | 1.88E-5 | 0E0 |
| 189 | Y | ZAP70 | TPQFTPYYVAPQVLE | 9.046 | 7.404 |
| 189 | Y | NTRK1 | TPQFTPYYVAPQVLE | 2.17E-7 | 0E0 |
| 189 | Y | NTRK2 | TPQFTPYYVAPQVLE | 1.6E-6 | 0E0 |
| 189 | Y | LRRK | TPQFTPYYVAPQVLE | 2.196 | 1.645 |
| 189 | Y | PTK6 | TPQFTPYYVAPQVLE | 22.218 | 21.741 |
| 189 | Y | LCK | TPQFTPYYVAPQVLE | 6.843 | 6.736 |
| 189 | Y | MEK4 | TPQFTPYYVAPQVLE | 3.204 | 2.777 |
| 189 | Y | LRRK2 | TPQFTPYYVAPQVLE | 2.196 | 1.645 |
| 189 | Y | MAPK1 | TPQFTPYYVAPQVLE | 1.743 | 1.707 |
| 189 | Y | MAP2K4 | TPQFTPYYVAPQVLE | 3.204 | 2.777 |
| 216 | Y | Ack | IPTSPTPYTYNKSCD | 1.68E-6 | 0E0 |
| 216 | Y | TNK2 | IPTSPTPYTYNKSCD | 1.68E-6 | 0E0 |
| 216 | Y | ERBB4 | IPTSPTPYTYNKSCD | 7.06E-7 | 0E0 |
| 216 | Y | PTK2 | IPTSPTPYTYNKSCD | 4.6E-5 | 0E0 |
| 216 | Y | NTRK1 | IPTSPTPYTYNKSCD | 4.53E-7 | 0E0 |
| 216 | Y | NTRK2 | IPTSPTPYTYNKSCD | 3.16E-6 | 0E0 |
| 216 | Y | BAZ | IPTSPTPYTYNKSCD | -1.528 | -1.564 |
| 216 | Y | BAZ1B | IPTSPTPYTYNKSCD | -1.528 | -1.564 |
| 218 | Y | Ack | TSPTPYTYNKSCDLW | 5.71E-6 | 0E0 |
| 218 | Y | Eph | TSPTPYTYNKSCDLW | 1.508 | 1.41 |
| 218 | Y | FGFR | TSPTPYTYNKSCDLW | 391.447 | 379.897 |
| 218 | Y | TNK2 | TSPTPYTYNKSCDLW | 5.71E-6 | 0E0 |
| 218 | Y | MATK | TSPTPYTYNKSCDLW | 2.08 | 2.01 |
| 218 | Y | ERBB4 | TSPTPYTYNKSCDLW | 2.11E-7 | 0E0 |
| 218 | Y | FGFR3 | TSPTPYTYNKSCDLW | 1.55E-6 | 0E0 |
| 218 | Y | PTK2 | TSPTPYTYNKSCDLW | 2.46E-5 | 0E0 |
| 218 | Y | NTRK1 | TSPTPYTYNKSCDLW | 1.66E-7 | 0E0 |
| 218 | Y | NTRK2 | TSPTPYTYNKSCDLW | 1.29E-6 | 0E0 |
| 218 | Y | BAZ | TSPTPYTYNKSCDLW | -1.559 | -1.564 |
| 218 | Y | BAZ1B | TSPTPYTYNKSCDLW | -1.559 | -1.564 |
| 232 | Y | Ack | WSLGVIIYVMLCGYP | 1.09E-8 | 0E0 |
| 232 | Y | TNK2 | WSLGVIIYVMLCGYP | 1.09E-8 | 0E0 |
| 232 | Y | ERBB4 | WSLGVIIYVMLCGYP | 1.22E-7 | 0E0 |
| 232 | Y | FGFR3 | WSLGVIIYVMLCGYP | 6.99E-7 | 0E0 |
| 232 | Y | NTRK1 | WSLGVIIYVMLCGYP | 6.68E-8 | 0E0 |
| 232 | Y | NTRK2 | WSLGVIIYVMLCGYP | 2.55E-6 | 0E0 |
| 232 | Y | BAZ | WSLGVIIYVMLCGYP | -1.313 | -1.564 |
| 232 | Y | BAZ1B | WSLGVIIYVMLCGYP | -1.313 | -1.564 |
| 238 | Y | DDR | IYVMLCGYPPFYSKH | 1.8 | 1.635 |
| 238 | Y | DDR2 | IYVMLCGYPPFYSKH | 1.8 | 1.635 |
| 238 | Y | EPHB5 | IYVMLCGYPPFYSKH | 1.395 | 1.207 |
| 238 | Y | FGFR3 | IYVMLCGYPPFYSKH | 9.96E-8 | 0E0 |
| 238 | Y | PTK2 | IYVMLCGYPPFYSKH | 2.09E-5 | 0E0 |
| 238 | Y | CSF1R | IYVMLCGYPPFYSKH | 1.91E-4 | 0E0 |
| 238 | Y | NTRK1 | IYVMLCGYPPFYSKH | 6.47E-8 | 0E0 |
| 238 | Y | NTRK2 | IYVMLCGYPPFYSKH | 3.5E-6 | 0E0 |
| 238 | Y | BAZ | IYVMLCGYPPFYSKH | -1.27 | -1.564 |
| 238 | Y | BAZ1B | IYVMLCGYPPFYSKH | -1.27 | -1.564 |
| 242 | Y | Ack | LCGYPPFYSKHHSRT | 4.34E-6 | 0E0 |
| 242 | Y | Trk | LCGYPPFYSKHHSRT | 67.227 | 41.671 |
| 242 | Y | TNK2 | LCGYPPFYSKHHSRT | 4.34E-6 | 0E0 |
| 242 | Y | ERBB4 | LCGYPPFYSKHHSRT | 2.12E-6 | 0E0 |
| 242 | Y | EPHA8 | LCGYPPFYSKHHSRT | 2.211 | 2.005 |
| 242 | Y | FGFR3 | LCGYPPFYSKHHSRT | 1.97E-7 | 0E0 |
| 242 | Y | PTK2 | LCGYPPFYSKHHSRT | 4.33E-5 | 0E0 |
| 242 | Y | CSF1R | LCGYPPFYSKHHSRT | 1.66E-4 | 0E0 |
| 242 | Y | NTRK1 | LCGYPPFYSKHHSRT | 2.02E-7 | 0E0 |
| 242 | Y | NTRK2 | LCGYPPFYSKHHSRT | 5.92E-6 | 0E0 |
| 242 | Y | BAZ | LCGYPPFYSKHHSRT | -1.401 | -1.564 |
| 242 | Y | MAPK | LCGYPPFYSKHHSRT | 2.044 | 1.861 |
| 242 | Y | BAZ1B | LCGYPPFYSKHHSRT | -1.401 | -1.564 |
| 242 | Y | MEK4 | LCGYPPFYSKHHSRT | 3.042 | 2.777 |
| 242 | Y | MAP2K3 | LCGYPPFYSKHHSRT | 1.458 | 0.818 |
| 242 | Y | MAP2K4 | LCGYPPFYSKHHSRT | 3.042 | 2.777 |
| 375 | Y | Ack | TKPKDGIYIHDHENG | 4.15E-6 | 0E0 |
| 375 | Y | TNK2 | TKPKDGIYIHDHENG | 4.15E-6 | 0E0 |
| 375 | Y | FGFR2 | TKPKDGIYIHDHENG | 2.277 | 1.727 |
| 375 | Y | FGFR3 | TKPKDGIYIHDHENG | 9.47E-7 | 0E0 |
| 375 | Y | PTK2 | TKPKDGIYIHDHENG | 1.88E-5 | 0E0 |
| 375 | Y | MST1R | TKPKDGIYIHDHENG | 0.001 | 0.001 |
| 375 | Y | TRK2 | TKPKDGIYIHDHENG | 6.35E-7 | 0E0 |
| 375 | Y | FLT1 | TKPKDGIYIHDHENG | 2.591 | 2.36 |
| 375 | Y | DYRK1 | TKPKDGIYIHDHENG | 1.74 | 1.399 |
| 375 | Y | DYRK2 | TKPKDGIYIHDHENG | 1.317 | 1.304 |
| 425 | Y | Ack | VMQEAWKYNRECKLL | 1.85E-6 | 0E0 |
| 425 | Y | Axl | VMQEAWKYNRECKLL | 0.016 | 0.015 |
| 425 | Y | Tec | VMQEAWKYNRECKLL | 14.927 | 14.555 |
| 425 | Y | TNK2 | VMQEAWKYNRECKLL | 1.85E-6 | 0E0 |
| 425 | Y | ERBB4 | VMQEAWKYNRECKLL | 6.48E-7 | 0E0 |
| 425 | Y | FGFR3 | VMQEAWKYNRECKLL | 7.22E-7 | 0E0 |
| 425 | Y | PTK2 | VMQEAWKYNRECKLL | 4.9E-5 | 0E0 |
| 425 | Y | CSF1R | VMQEAWKYNRECKLL | 2.71E-4 | 0E0 |
| 425 | Y | NTRK1 | VMQEAWKYNRECKLL | 1.2E-7 | 0E0 |
| 425 | Y | NTRK2 | VMQEAWKYNRECKLL | 1.98E-6 | 0E0 |
| 425 | Y | FLT4 | VMQEAWKYNRECKLL | 39.878 | 36.853 |

Group-based Prediction System (GPS; http://gps.biocuckoo.cn) output predicting tyrosine residues phosphorylated in mouse MK5.1 (NCBI NP_034895.1) and their cognate protein kinases. The peptide sequence shows the predicted phosphorylated amino acid with 7 flanking amino acids. The score refers to the value calculated by GPS5 evaluate the theoretical potential a protein kinase phosphorylating the indicated residue ( higher value = higher potential. The cut-off value for a predicted phosphorylation site using e high threshold indicates a value resulting in a 4% false positive rate.
